# Expanding the Genetic Architecture of Nicotine Dependence and its Shared Genetics with Multiple Traits: Findings from the Nicotine Dependence GenOmics (iNDiGO) Consortium

**DOI:** 10.1101/2020.01.15.898858

**Authors:** Bryan C. Quach, Michael J. Bray, Nathan C. Gaddis, Mengzhen Liu, Teemu Palviainen, Camelia C. Minica, Stephanie Zellers, Richard Sherva, Fazil Aliev, Michael Nothnagel, Kendra A. Young, Jesse A. Marks, Hannah Young, Megan U. Carnes, Yuelong Guo, Alex Waldrop, Nancy Y.A. Sey, Maria T. Landi, Daniel W. McNeil, Dmitriy Drichel, Lindsay A. Farrer, Christina A. Markunas, Jacqueline M. Vink, Jouke-Jan Hottenga, William G. Iacono, Henry R. Kranzler, Nancy L. Saccone, Michael C. Neale, Pamela Madden, Marcella Rietschel, Mary L. Marazita, Matthew McGue, Hyejung Won, Georg Winterer and the German Nicotine Cohort Study, Richard Grucza, Danielle M. Dick, Joel Gelernter, Neil E. Caporaso, Timothy B. Baker, Dorret I. Boomsma, Jaakko Kaprio, John E. Hokanson, Scott Vrieze, Laura J. Bierut, Eric O. Johnson, Dana B. Hancock

**Affiliations:** GenOmics, Bioinformatics, and Translational Research Center, Biostatistics and Epidemiology Division, RTI International, Research Triangle Park, North Carolina, USA; Department of Psychiatry, Washington University, St. Louis, Missouri, USA; Department of Psychology, University of Minnesota Twin Cities, Minneapolis, Minnesota, USA; Institute for Molecular Medicine Finland (FIMM), University of Helsinki, Helsinki, Finland; Department of Biological Psychology, Vrije Universiteit, Amsterdam, The Netherlands; Department of Medicine (Biomedical Genetics), Boston University School of Medicine, Boston, Massachusetts, USA; Department of Psychology, Virginia Commonwealth University, Richmond, Virginia, USA; Faculty of Business, Karabuk University, Turkey; Cologne Center for Genomics, University of Cologne, Cologne, Germany; University Hospital Cologne, Cologne, Germany; Department of Epidemiology, University of Colorado Anschutz Medical Campus, Aurora, Colorado, USA; GeneCentric Therapeutics, Research Triangle Park, North Carolina, USA; Department of Genetics, University of North Carolina at Chapel Hill, Chapel Hill, North Carolina, USA; Genetic Epidemiology Branch, Division of Cancer Epidemiology and Genetics, National Cancer Institute, National Institutes of Health, United States department of Health and Human Services, Bethesda, Maryland, USA; Department of Psychology, West Virginia University, Morgantown, West Virginia, USA; Department of Dental Practice and Rural Health, West Virginia University, Morgantown, West Virginia, USA; Department of Neurology, Boston University School of Medicine, Boston, Massachusetts, USA; Department of Ophthalmology, Boston University School of Medicine, Boston, Massachusetts, USA; Department of Epidemiology, Boston University School of Public Health, Boston, Massachusetts, USA; Department of Biostatistics, Boston University School of Public Health, Boston, Massachusetts, USA; Behavioural Science Institute, Radboud University, Nijmegen, The Netherlands; Department of Psychiatry, University of Pennsylvania Perelman School of Medicine, Philadelphia, Pennsylvania, USA; VISN 4 MIRECC, Crescenz VA Medical Center, Philadelphia, Pennsylvania, USA; Department of Genetics, Washington University, St. Louis, Missouri, USA; Division of Biostatistics, Washington University, St. Louis, Missouri, USA; Virginia Institute for Psychiatric and Behavioral Genetics, Virginia Commonwealth University, Richmond, Virginia, USA; Department of Psychiatry, Virginia Commonwealth University, Richmond, Virginia, USA; Department of Genetic Epidemiology in Psychiatry, Central Institute of Mental Health, Medical Faculty Mannheim, University of Heidelberg, Mannheim, Germany; Center for Craniofacial and Dental Genetics, Department of Oral Biology, University of Pittsburgh, Pittsburgh, Pennsylvania, USA; Experimental & Clinical Research Center, Department of Anesthesiology and Operative Intensive Care Medicine, Charité - University Medicine Berlin, Berlin, Germany; Departments of Family and Community Medicine and Health and Clinical Outcomes Research, Saint Louis University, St. Louis, Missouri, USA; College Behavioral and Emotional Health Institute, Virginia Commonwealth University, Richmond, Virginia, USA; Department of Human & Molecular Genetics, Virginia Commonwealth University, Richmond, Virginia, USA; Department of Psychiatry, Yale University School of Medicine, New Haven, Connecticut, USA; Department of Genetics, Yale University School of Medicine, New Haven, Connecticut, USA; Department of Neuroscience, Yale University School of Medicine, New Haven, Connecticut, USA; Department of Psychiatry, VA CT Healthcare Center, West Haven, Connecticut, USA; Occupation and Environmental Epidemiology Branch, Division of Cancer Epidemiology and Genetics, National Cancer Institute, National Institutes of Health, United States Department of Health and Human Services, Bethesda, Maryland, USA; Center for Tobacco Research and Intervention, Department of Medicine, University of Wisconsin School of Medicine and Public Health, Madison, Wisconsin, USA; Department of Public Health, Faculty of Medicine, University of Helsinki, Helsinki, Finland; Fellow Program, Behavioral Health Research Division, RTI International, Research Triangle Park, North Carolina, USA

## Abstract

Cigarette smoking is the leading cause of preventable morbidity and mortality. Knowledge is evolving on genetics underlying initiation, regular smoking, nicotine dependence (ND), and cessation. We performed a genome-wide association study using the Fagerström Test for ND (FTND) in 58,000 smokers of European or African ancestry. Five genome-wide significant loci, including two novel loci *MAGI2/GNAI1* (rs2714700) and *TENM2* (rs1862416) were identified, and loci reported for other smoking traits were extended to ND. Using the heaviness of smoking index (HSI) in the UK Biobank (N=33,791), rs2714700 was consistently associated, but rs1862416 was not associated, likely reflecting ND features not captured by the HSI. Both variants were *cis*-eQTLs (rs2714700 for *MAGI2-AS3* in hippocampus, rs1862416 for *TENM2* in lung), and expression of genes spanning ND-associated variants was enriched in cerebellum. SNP-based heritability of ND was 8.6%, and ND was genetically correlated with 17 other smoking traits (r_g_=0.40–0.95) and co-morbidities. Our results emphasize the FTND as a composite phenotype that expands genetic knowledge of smoking, including loci specific to ND.

## Introduction

Cigarette smoking remains the leading cause of preventable death worldwide^1^ despite the well-known adverse health effects. Smoking causes more than 7 million deaths annually from a multitude of diseases including cancer, chronic obstructive pulmonary disease (COPD), and heart disease.^1,2^ Cigarette smoking is a multi-stage process consisting of initiation, regular smoking, nicotine dependence (ND), and cessation. Each step has a strong genetic component (for example, twin-based heritability estimates up to 70% for the transition from regular smoking to ND^3,4^), and partial overlaps are expected among the sets of sequence variants correlating with the different stages,^3^ as evidenced by findings of the GWAS and Sequencing Consortium of Alcohol and Nicotine use (GSCAN) with sample sizes up to 1.2 million individuals.^5^ GSCAN identified 298 genome-wide significant loci associated with initiation (ever vs. never smoking), age at initiation, cigarettes per day (CPD), and/or cessation (current vs. former smoking); 259 of the loci harbored significant associations with initiation.^1^

In comparison to other stages of smoking, known loci for ND are limited. Only six reproducible, genome-wide significant loci have been identified: *CHRNB3-CHRNA6* (chr8p11), *DBH* (chr9q34), *CHRNA5-CHRNA3-CHRNB4* (chr15q25), *DNMT3B* and *NOL4L* (chr20q11), and *CHRNA4* (chr20q13).^6^ A more complete understanding of the genetics underlying ND is needed, as it could help to predict the likelihood of quitting smoking, withdrawal severity, response to treatment, and health-related consequences.^7-10^ The Fagerström Test for ND (FTND), also called the Fagerström Test for Cigarette Dependence,^11^provides a composite phenotype that captures multiple behavioral and psychological features of ND among smokers.^12^ While CPD is associated with key markers of ND, such as cessation likelihood^13^, the FTND conveys additional valuable information by including 5 items in addition to CPD. FTND is meaningfully associated with tobacco use diagnostic criteria from the *Diagnostic and Statistical Manual of Mental Disorders*^14,15^ and is more highly associated with withdrawal severity than is CPD^7^ Its validity may be due to the inclusion of the time-to-first-cigarette in the morning (TTFC) item, which appears to be especially strongly associated with relapse likelihood^16-18^ and may be an especially informative measure of heritability of ND.^19^Thus, the FTND provides somewhat different information than CPD alone and has been relatively understudied from a genetic perspective because of its more limited availability across datasets.

The FTND score, based on totaling responses to the 6 items that constitute the FTND, ranges from 0 (no dependence) to 10 (highest dependence level).^12,20^ In the present study, we categorized FTND scores as mild (scores 0–3), moderate (scores 4–6), or severe (scores 7–10), as done before in studies comprising our Nicotine Dependence GenOmics (iNDiGO) Consortium.^21,22^ We expand upon our prior analyses and report findings from the largest GWAS meta-analysis for ND (N=58,000; 46,213 European [EUR] and 11,787 African American [AA] ancestry participants from 23 studies) to identify novel genetic loci associated with ND, assess genetic correlations between ND and other phenotypes and gene expression patterns, and test GSCAN-identified loci5 for effects on ND.

## Results

### Cross-ancestry GWAS meta-analysis finds two novel SNP associations with ND

Our cross-ancestry ND GWAS meta-analysis (λ=1.034, **Supplementary Figure 1A**) identified five genome-wide significant loci (**Figure 1**). Associations of the lead SNPs (single nucleotide polymorphisms) from each of these five loci are shown in **Table 1**. All genome-wide significant SNP/indel associations from the cross-ancestry meta-analysis are provided in **Supplementary Table 1**.

**Table 1.**
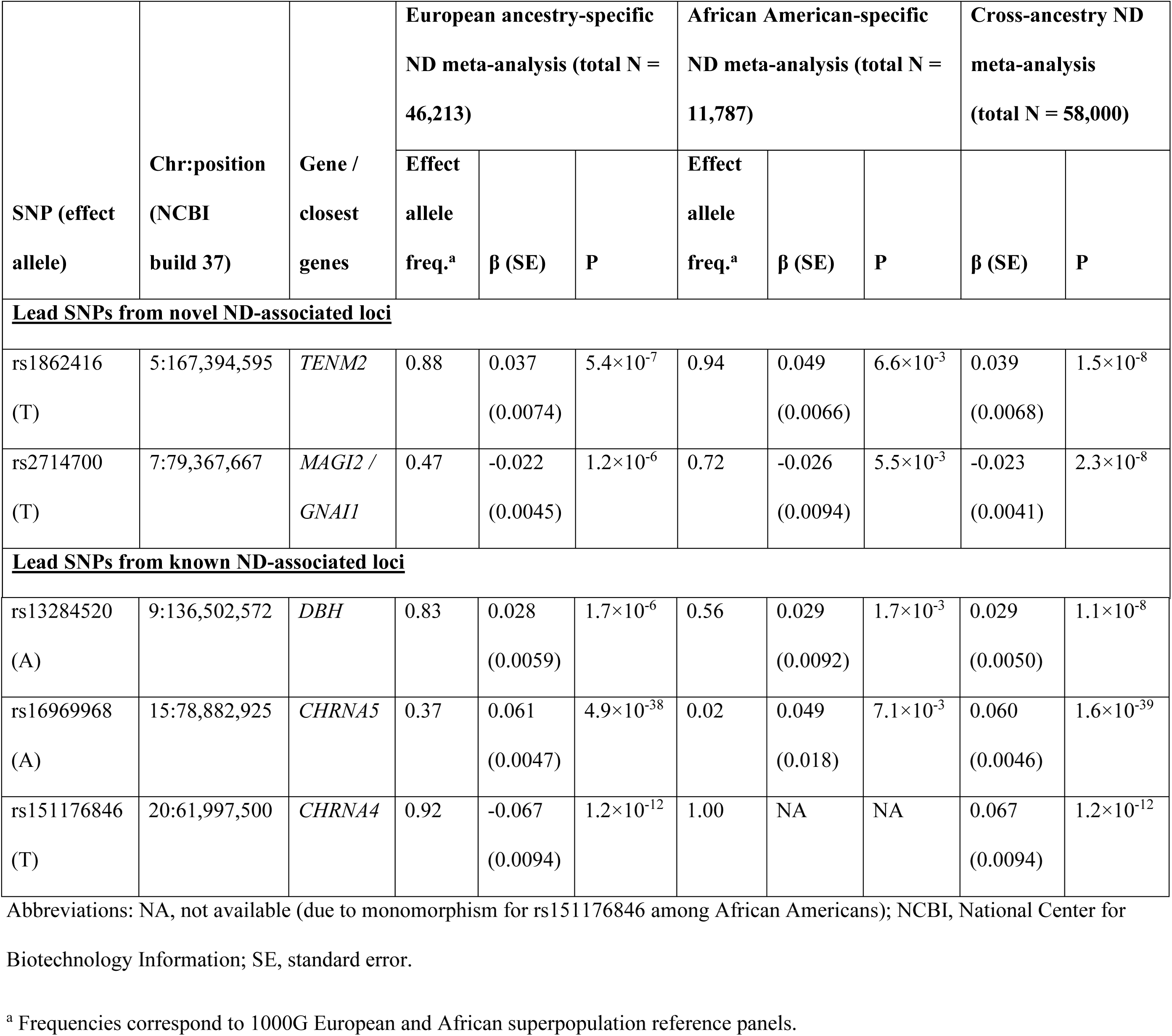
Lead single nucleotide polymorphism (SNP) associations from the five genome-wide significant loci in the Nicotine Dependence GenOmics (iNDiGO) consortium cross-ancestry meta-analysis for nicotine dependence (ND). Ancestry-specific association results are also presented.

**Figure 1.**
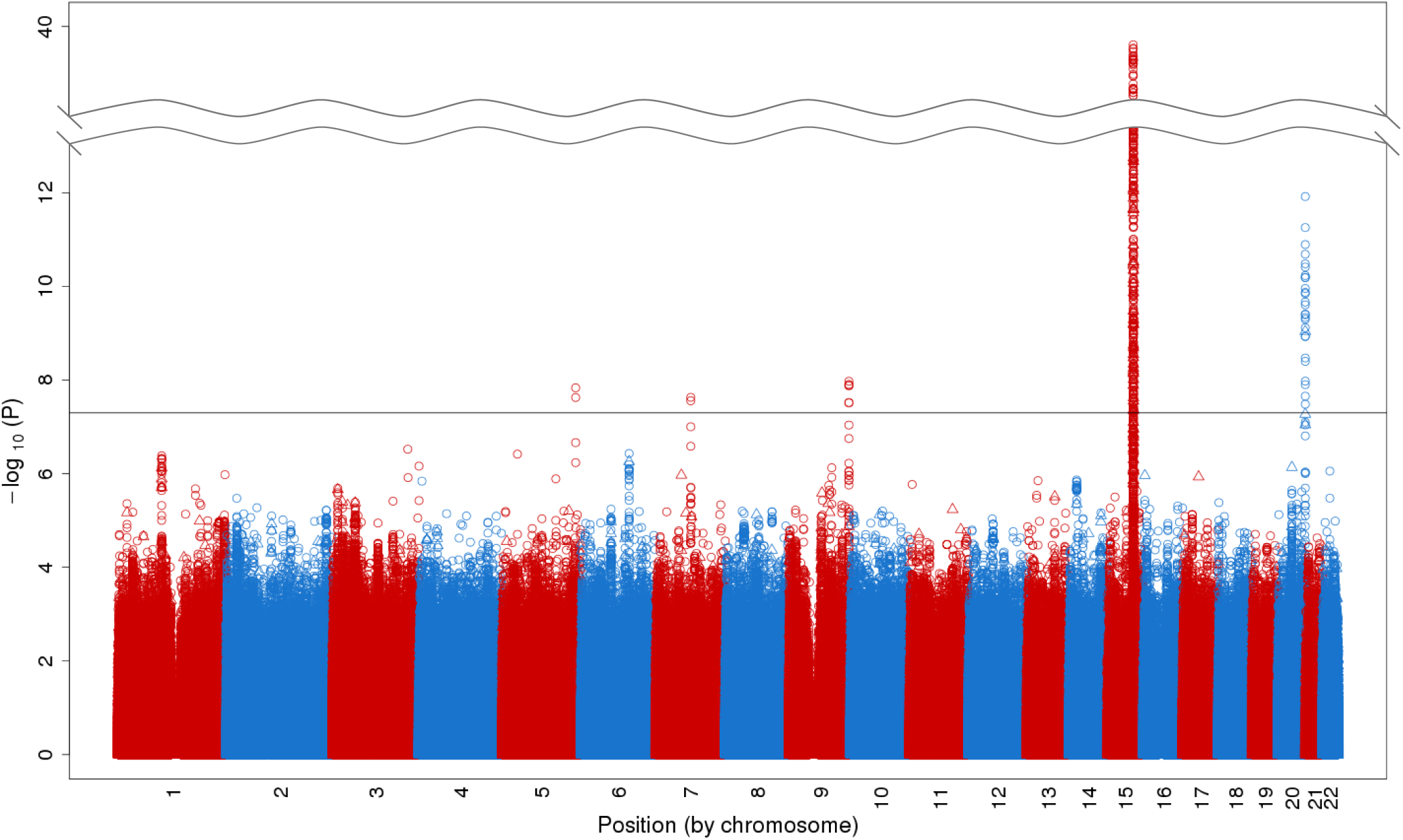
Cross-ancestry nicotine dependence genome-wide association meta-analysis results, comprising 23 iNDiGO studies with total N = 58,000 European and African American ancestry ever smokers. The –log_10_ meta-analysis p-values of single nucleotide polymorphisms (SNPs; depicted as circles) and insertions/deletions (indels; depicted as triangles) are plotted by chromosomal position. Five loci surpassed the genome-wide statistical significance threshold (P<5×10^−8^, as marked by the solid horizontal black line).

Three of the genome-wide significant loci have known associations with ND from our prior GWAS and others^6^: chr15q25^21-23^ (smallest P=1.6×10^−39^ for rs16969968, a well-established functional missense [D398N] *CHRNA5* SNP^24^), chr20q13^21^ (smallest P=1.2×10^−12^ for rs151176846, an intronic *CHRNA4* SNP), and chr9q34^22^ (smallest P=1.1×10^−8^ for rs13284520, an intronic *DBH* SNP). In the EUR-specific GWAS meta-analysis, the loci spanning nicotinic acetylcholine receptor genes (*CHRNA5-A3-B4* and *CHRNA4*), but no novel loci, were identified at genome-wide significance (λ=1.036, **Supplementary Figures 1B** and **2A**). No genome-wide significant loci were found in the GWAS meta-analysis among AAs (λ=1.032, **Supplementary Figures 1C** and **2B**).

Two genome-wide significant loci from the cross-ancestry meta-analysis represent novel associations with ND. On chr7q21, the most significant SNP (P=2.3×10^−9^) was rs2714700, a SNP between the *MAGI2* and *GNAI1* genes (**Supplementary Figures 3A–B**). The most significant SNP on chr5q34, rs1862416 (P=1.5×10^−8^), sits within an intron for *TENM2* (**Supplementary Figures 3C–D**). Both SNPs imputed well: sample size-weighted mean estimated r^2^ values were 0.97 for rs2714700 and 0.92 for rs1862416. Further, both SNPs were common, and their associations with ND were observed across EURs and AAs (**Table 1**) and were largely consistent across studies (**Supplementary Figure 4A–B**): rs2714700-T being associated with reduced risk (meta-analysis odds ratio [OR] and 95% confidence interval [CI]=0.96 [0.94–0.97]) and rs1862416-T being associated with increased risk (meta-analysis OR [95% CI]=1.08 [1.05– 1.11]) for severe vs. mild ND. These comparisons of dissimilar categories were derived from the GWAS regression coefficients (i.e., OR=exp[2×β] for severe vs. mild ND, with OR>1 corresponding to an increased risk of severe ND) to contextualize the magnitude of the observed effect sizes. Neither SNP showed evidence for heterogeneity, based on the I^2^index^25^, across studies (P=0.83 for rs2714700 and 0.75 for rs1862416). Leave-one-study-out analyses (**Supplementary Table 2**) revealed some variability in p-values (P=3.1×10^−7^–7.4×10^−9^ for rs2714700 and P=5.6×10^−9^–3.9×10^−6^ for rs1862416), likely due to fluctuating statistical power given the significant correlation between N and p-value across iterations: r=-0.65, P=8.6×10^−5^. However, there was little variation in the effect sizes (range of β values corresponding to the OR for severe vs. mild ND = 0.95–0.96 for rs2714700-T and 1.07–1.08 for rs1862416-T).

**Table 2.**
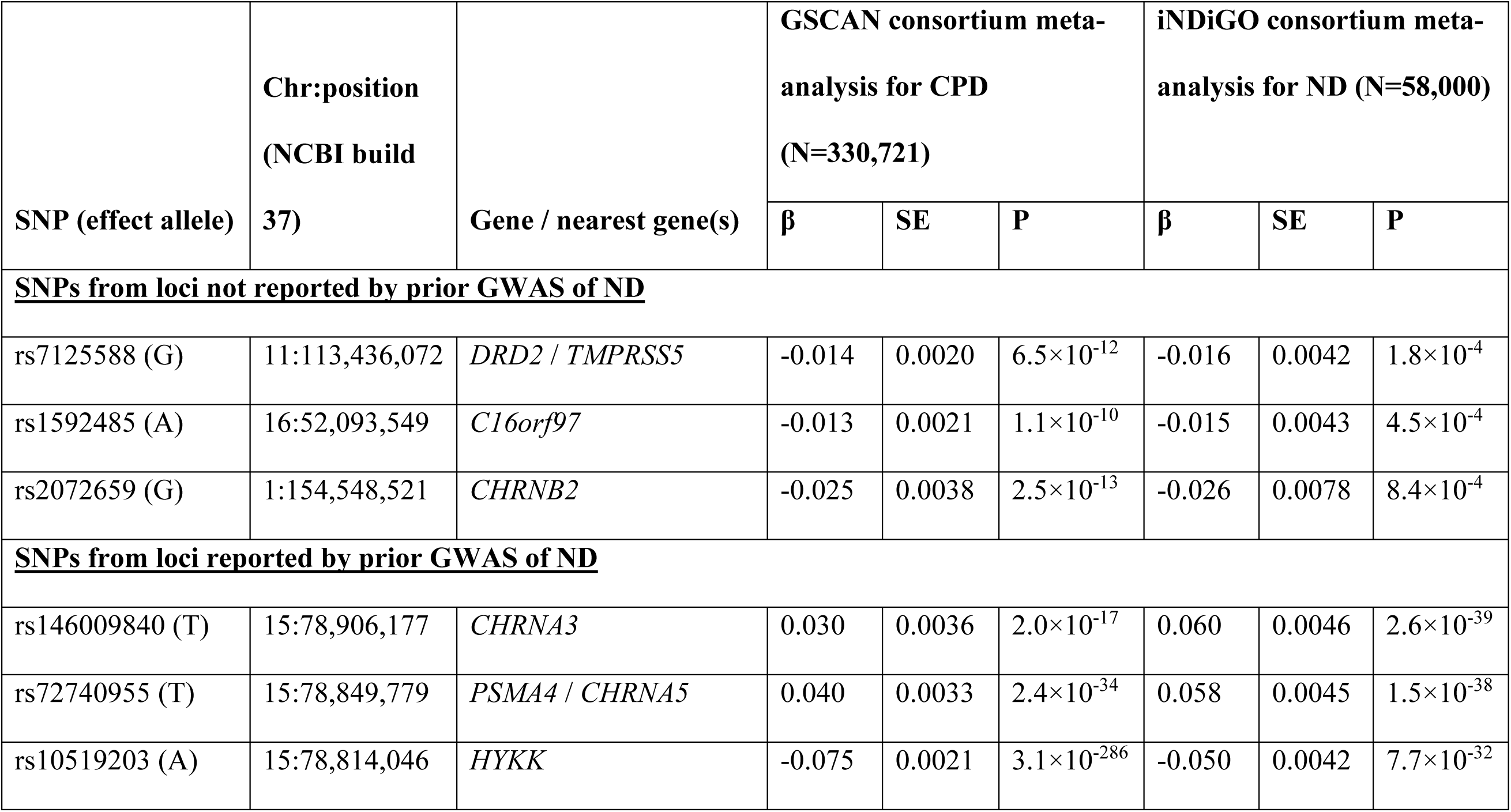
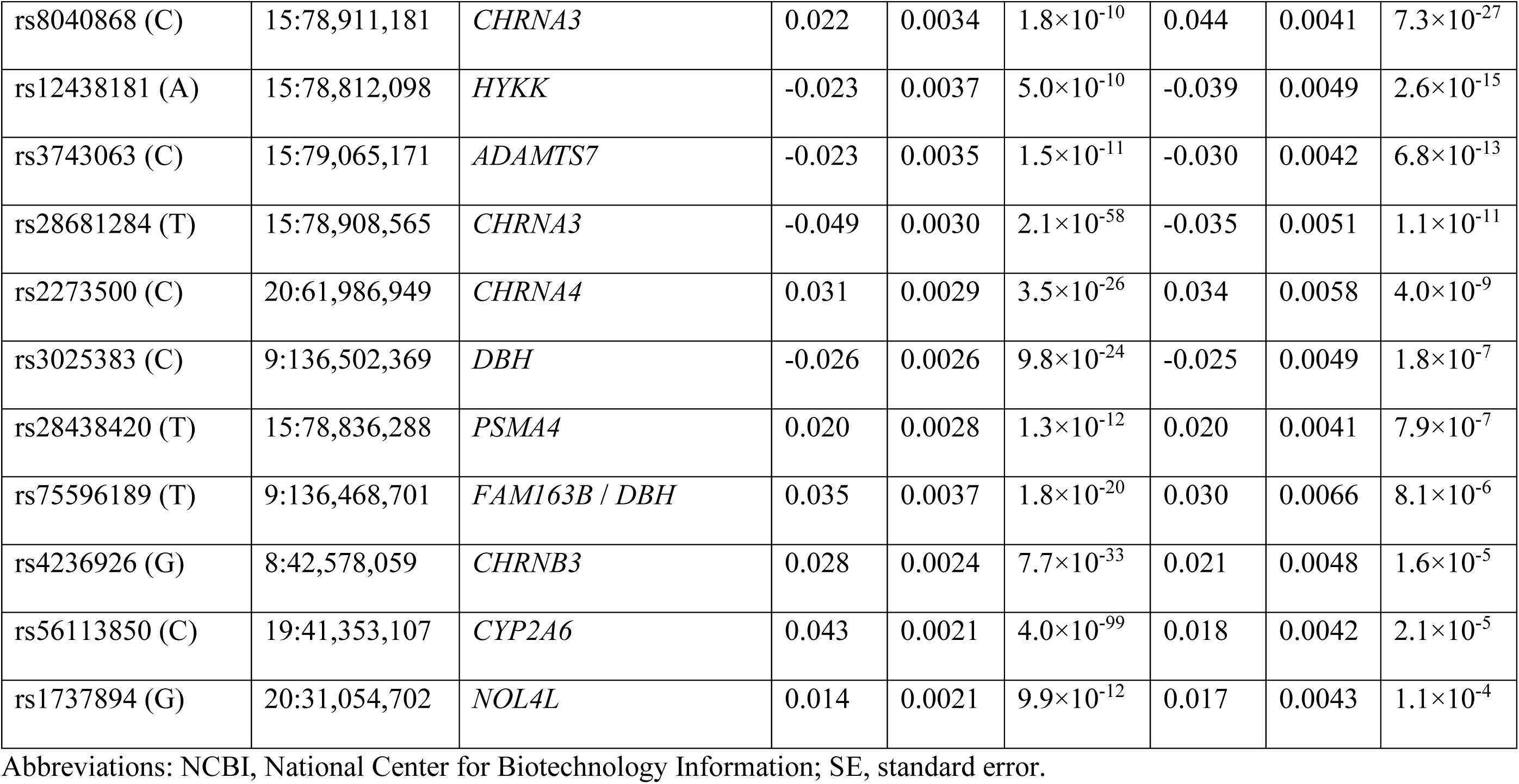
Single nucleotide polymorphisms (SNPs) identified as genome-wide significant for cigarettes per day (CPD) by the GWAS and Sequencing Consortium of Alcohol and Nicotine use (GSCAN) consortium and associated with nicotine dependence (ND) at P<9.1×10^−4^ (α=0.05/55 tests) in the cross-ancestry meta-analysis by the Nicotine Dependence GenOmics (iNDiGO) consortium. Results are sorted by novelty and then by iNDiGO p-values, and β values correspond to direction of association for the effect alleles.

**Figure 2.**
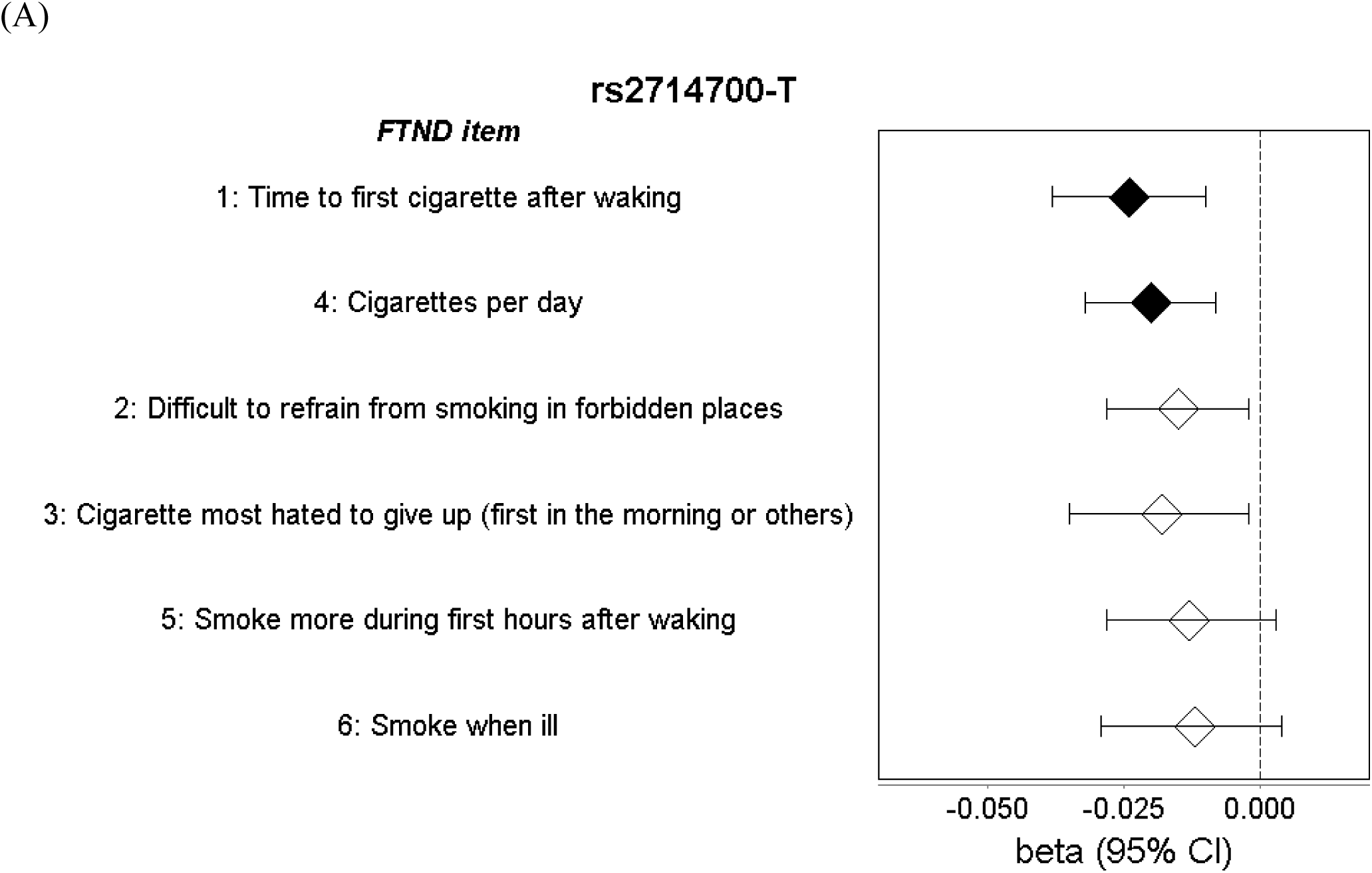
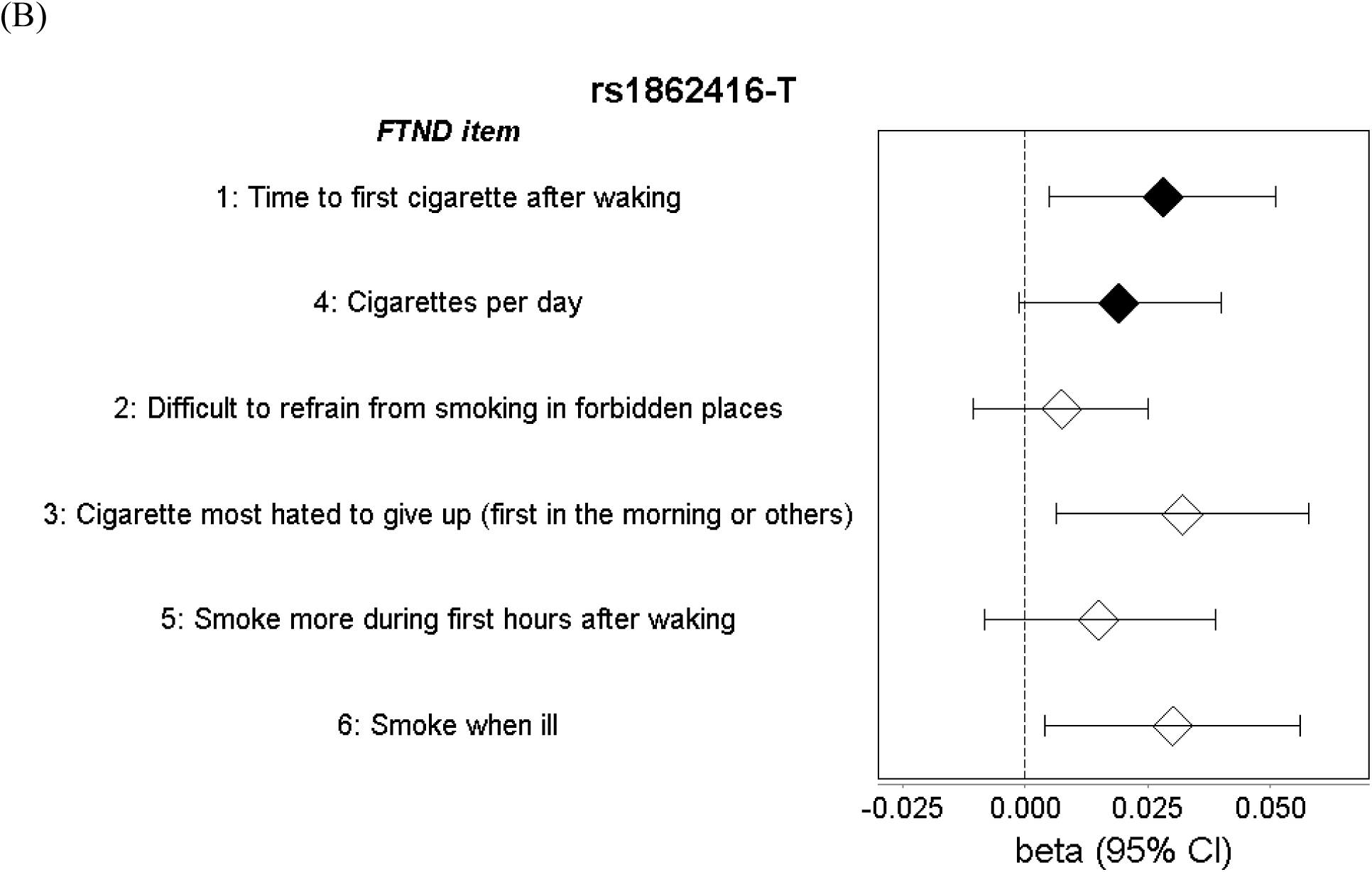
Associations of novel single nucleotide polymorphisms (SNPs) with specific items of the Fagerström Test for Nicotine Dependence (FTND) across the iNDiGO studies. Associations are presented from cross-ancestry meta-analyses of the (A) *MAGI2/GNAI1* SNP allele rs2714700-T and (B) *TENM2* SNP allele rs1862416-T. Beta (β) and corresponding 95% confidence interval (CI) estimates were taken from linear regression models for categorical FTND item responses (1 and 4, closed diamonds) or logistic regression models for binary FTND item responses (2, 3, 5, and 6, open diamonds).

**Figure 3.**
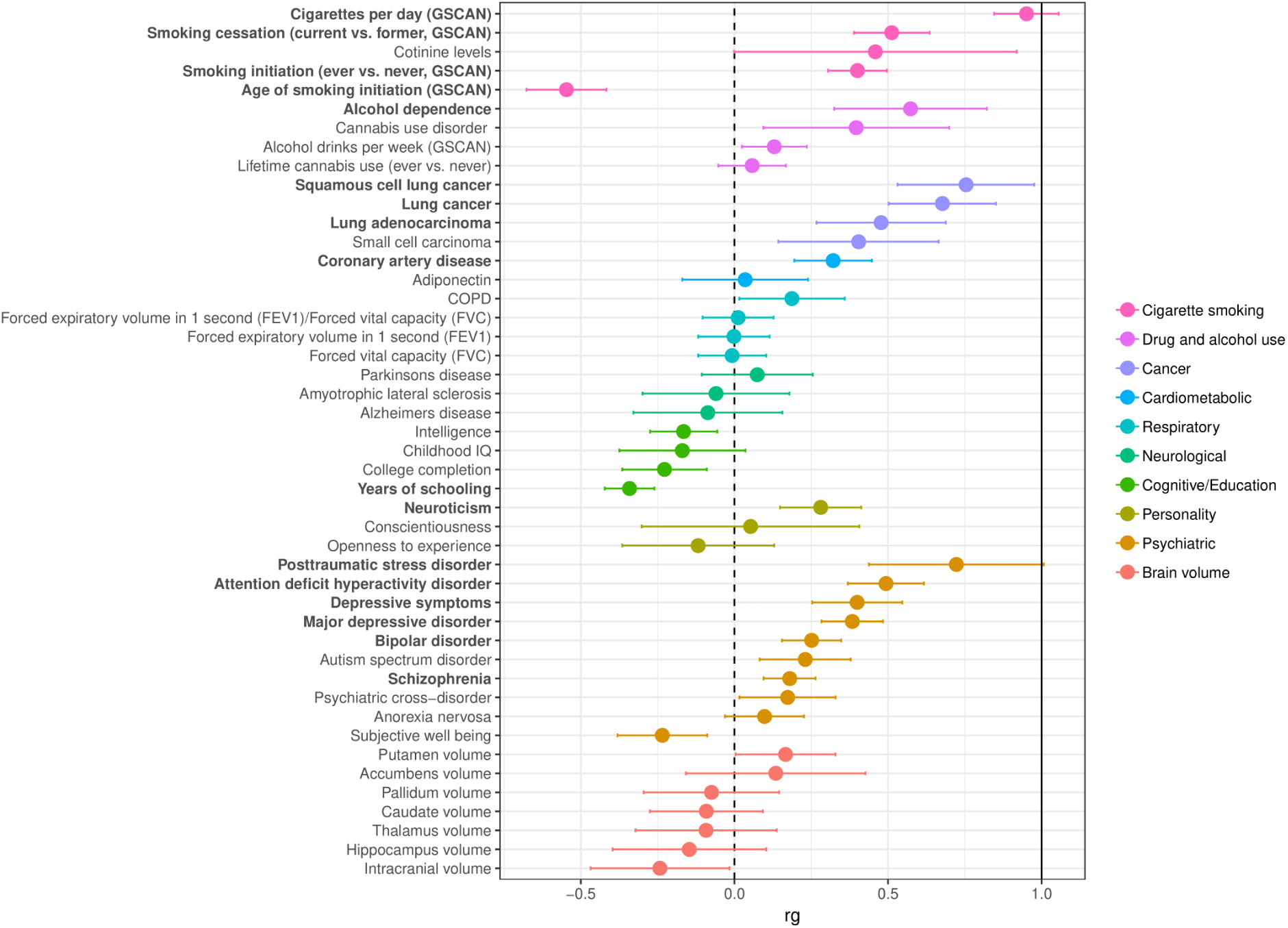
Genetic correlations of nicotine dependence (ND) with 46 other phenotypes. Correlations were calculated using linkage disequilibrium (LD) score regression with the iNDiGO European ancestry-specific GWAS meta-analysis results for ND (N=46,213), compared with results made available via LD Hub or study investigators (see Supplementary Table 3 for original references). Phenotypes were grouped by disease/trait or measurement category, as indicated by different colorings. Point estimates equate to genetic correlation (r_g_) values; error bars show the 95% confidence intervals; and the dotted vertical grey line corresponds to r_g_ =0 (no correlation with ND). Phenotypes with significant correlations (P<0.0011, α=0.05/46 tested) are bolded.

We compared the novel ND-associated SNPs with results reported for other smoking traits by GSCAN.^5^ European ancestry participants from 8 iNDiGO studies were included in GSCAN (**Supplementary Table 3**). Both the *MAGI2/GNAI1* SNP rs2714700 and the *TENM2* SNP rs1862416 were nominally associated at P<0.05 with ever vs. never smoking and rs2714700 with CPD in consistent directions with ND; neither SNP was associated with age at initiation or smoking cessation (**Supplementary Table 4**).

For replication in an independent sample, we analyzed the two novel SNPs (rs2714700 and rs1862416) for association with the heaviness of smoking index (HSI) in the UK Biobank. Results are shown in **Supplementary Table 6**. HSI is based on two items (CPD and TTFC) of the 6 items that constitute the FTND; the HSI and full-scale FTND are highly correlated (e.g., r=0.7 among nondaily smokers and 0.9 among daily smokers).^26^ The *MAGI2/GNAI1* SNP, rs2714700, was associated with HSI at P=0.014, which surpassed Bonferroni correction for two SNP tests, and meta-analysis of iNDiGO studies with UK Biobank (total N=91,791) supported rs2714700-T being associated with milder ND (P=7.7×10^−9^). The *TENM2* SNP, rs1862416, was not associated with HSI in the UK Biobank (P=0.39).

To determine whether the novel genome-wide associations were driven by specific FTND items or shared across items, we returned to the iNDiGO studies, tested SNP associations with each specific FTND item, and combined results via cross-ancestry meta-analyses. For rs2714700, we observed the lowest p-values for the two items that constitute the HSI (**Figure 2A**): TTFC (P=5.3×10^−4^) and CPD (P=1.1×10^−3^). Rs2714700 was also associated at P<0.05 with difficult in refraining from smoking in forbidden places (P=0.025) and the cigarette most hated to give up (P=0.030). Rs1862416 was associated with TTFC (P=0.018) and two items that are not captured by the HSI: the cigarette most hated to give up (P=0.015) and smoking when ill (P=0.023) (**Figure 2B**).

### GWAS findings for other smoking traits extend to ND

We assessed whether genome-wide significant SNPs identified for smoking traits in GSCAN extended to ND using results from the cross-ancestry GWAS meta-analysis. We focused on the 55 genome-wide significant SNPs from 40 loci associated with CPD, given that it displayed the best genetic correlation with ND (**Figure 3**). After applying Bonferroni correction for the 53 SNPs that were available in our meta-analysis (P<9.4×10^−4^), 17 SNPs had a statistically significant and directionally consistent association with ND (**Table 2**). These SNPs span six loci reported at genome-wide or nominal significance in prior GWAS of ND (*CHRNA5-A3-B4* [chr15], *CHRNA4* [chr20], *DBH* [chr9], *CHRNB3* [chr8], *CYP2A6* [chr19], and *NOL4L* [near *DNMT3B*, chr20])^6^ and three loci not reported in prior ND GWAS—*DRD2* (chr11), *C16orf97* (chr16), and *CHRNB2* (chr1).

### Gene-based association analyses highlight known genetic loci

Using Hi-C coupled multi-marker analysis of genomic annotation (H-MAGMA)^27^ on the EUR-specific GWAS meta-analysis results from iNDiGO, 11 genes when using fetal brain tissue and 13 genes when using adult brain tissue were associated with ND at P<2.7×10^−6^, based on correction for testing 18,655 protein coding genes. See **Supplementary Tables 7** and **8** for the genome-wide H-MAGMA results for fetal and adult tissues, respectively. Of the 16 unique genes identified, 10 genes in three known loci were associated with HSI in the UK Biobank at P<0.0031, based on correction for testing 16 genes (**Supplementary Table 9**): the *ACSBG1-WDR61-IREB2-HYKK-PSMA4-CHRNA5-CHRNA3-CHRNAB4-ADAMTS7-MORF4L1* gene cluster on chr. 15q25, *CHRNA4* on chr. 20q13, and the *ADAMTSL2* and *DBH* genes in close proximity on chr. 9q34. Two novel genes on distinct chromosomes were identified in iNDiGO (*AFG1L* on chr. 6q21 and *AK2* on chr. 1p35), but their associations were not corroborated in UK Biobank.

We also applied Summary-MultiXcan (S-MultiXcan)^28^ to the EUR-specific GWAS meta-analysis results and found significant associations for two chromosome 15q25 genes (*PSMA4* and *CHRNA5*), when considering *cis*-eQTL evidence from either the multi-tissue or single best tissue (substantia nigra). See **Supplementary Table 10** for the genome-wide S-MultiXcan results. Both genes were also associated with HSI in UK Biobank from multi-tissue (P=2.4× 10^−8^ for *PSMA4* and 1.3×10^−6^ for *CHRNA5*) or single best tissue (P=9.6×10^−14^ for *PSMA4* and 4.6×10^−8^ for *CHRNA5*, both in substantia nigra).

### ND is genetically correlated with 17 other phenotypes

We estimated the heritability explained by common SNPs of ND at 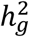 (standard error) = 0.086 (0.012), using LDSC^29^ and the EUR-specific GWAS meta-analysis results. We also found statistically significant genetic correlations of ND with 17 phenotypes (Bonferroni-corrected P<0.0011; **Figure 3** and **Supplementary Table 11**). Positive correlations indicate that the genetic predisposition to higher ND risk was correlated with genetic risks for other smoking traits^5^ (smallest P=3.1×10^−70^ for higher CPD [r_g_=0.95], followed by P=3.2×10^−16^ for current smoking [r_g_=0.54] and P=3.2×10^−16^ for ever smoking [r_g_=0.40]). We repeated LDSC, after removing all chr15q25 variants between 78.5 and 79.5 MB and found only negligible differences in these correlations (r_g_=0.94 for higher CPD, r_g_=0.51 for current smoking, and r_g_=0.42 for ever smoking). Beyond the smoking traits, with all SNPs included, higher ND was genetically correlated with higher risks of alcohol dependence,^30^ neuroticism,^31^ psychiatric diseases (attention deficit hyperactivity disorder,^32^ bipolar disorder,^33^ major depressive disorder^34^ and its symptoms,^31^ posttraumatic stress disorder, and schizophrenia^35^), and smoking-related consequences (lung cancer and its histological subtypes ^36^ and coronary artery disease ^37^). Among these positively correlated traits, r_g_ values ranged from 0.16 (schizophrenia) to 0.77 (squamous cell lung cancer). Higher risk of ND was genetically correlated with lower age of smoking initiation^5^ r_g_=-0.55) and fewer years of schooling^38^ (r_g_=-0.34).

For the traits with statistically significant genetic correlations with ND from the cigarette smoking, drug and alcohol use, personality, and psychiatric categories, we applied pairwise GWAS (GWAS-PW)^39^ to identify shared genetic influences between FTND and each of these traits (**Supplementary Figure 5**). GWAS-PW provides posterior probabilities for several models of genetic influence, including whether a given genomic region contains a variant that influences only ND (model 1), only the other trait (model 2), or both ND and the other trait (model 3). It also considers the scenario of whether the region contains a variant that influences ND and a separate variant influences the other trait (model 4). Both novel FTND-associated GWAS loci showed large probabilities for model 4 when comparing alcohol dependence and ND (posterior probabilities > 0.97). The region surrounding rs2714700 also showed large model 4 probabilities for comparisons with depressive symptoms and schizophrenia. The region surrounding rs1862416 exhibited large model 3 probabilities for major depressive disorder and smoking initiation.

Rs1862416 was located within the boundaries of a genome-wide significant locus for smoking initiation (chr5:164,596,435-168,114,971), and to assess the independence of association signals at the single variant level, we performed conditional modeling using Genome-wide Complex Trait Analysis (GCTA).^40,41^ All 6 lead SNPs in this GSCAN-identified locus were in low LD with rs1862416 (maximum r =0.0047 [**Supplementary Figure 6**], maximum D’=0.46), and three were nominally associated with ND at P<0.05 (**Supplementary Table 5**). Among our iNDiGO studies, rs1862416 remained associated with ND in models conditioned on each GSCAN lead SNP individually (P=7.9× 10^−8^–1.8×10 ^-8^) and with all 6 SNPs taken together (P=2.2×10^−7^). Rs2714700 was located >1 MB away from any GSCAN-identified locus, so conditional modeling was not necessary. Altogether, the GWAS-PW results suggest pleiotropy of smoking-related and comorbid traits in our two novel ND-associated regions, but at the variant level, the rs2714700 and rs1862416 associations with ND are independent of the GSCAN-identified variants.

### Gene expression data implicates target genes for novel ND-associated SNPs and identifies ND heritability enrichment in cerebellum

Credible set analysis of the chr7q21 locus narrowed the list of most likely causal variants to the lead SNP (rs2714700) and three others (rs2714674, rs1464692, and rs2707864) (**Supplementary Table 12**). Rs2714700, an intergenic SNP, is not a significant *cis*-eQTL with any gene-level expression in the Genotype-Tissue Expression (GTEx; v8) project, but it was implicated as a *cis*-eQTL for the *MAGI2-AS3* transcript in hippocampus from BrainSeq ^42^ (N=551; P=8.5×10 ^-4^). The protective allele for ND (rs2714700-T) was associated with higher expression of the *MAGI2-AS3* transcript ENST00000414797.5. Rs1464692 was also implicated as a *cis*-eQTL for the *MAGI2-AS3* transcript in hippocampus from BrainSeq (N=551; P=8.1×10 ^-4^), and rs2707864 is located in a DNaseI hypersensitivity site in adult and fetal fibroblast cells in HaploReg ^39^ (**Supplementary Table 12**).

The lead SNP at the chr5q34 locus, rs1862416, is annotated to enhancer histone marks in brain (specifically, germinal matrix during fetal development and the developed prefrontal cortex, anterior caudate, and cingulate gyrus tissues) and several other tissues in HaploReg. ^43^ It is also located in the promoter of *CTB-77H17*.*1*, which is a novel antisense RNA transcript encoded within a *TENM2* intron. In GTEx, rs1862416 was reported as a significant lung-specific *cis*-eQTL SNP *TENM2*. The ND risk-conferring allele (rs1862416-T) was associated with decreased gene-level *TENM2* expression in lung. *CTB-77H17*.*1* was too lowly expressed across GTEx tissues to test its expression levels by rs1862416. Two additional, potentially causal variants identified in a credible set analysis were similarly annotated to enhancer and promoter markers in brain (prefrontal cortex, astrocyte) and fetal lung in HaploReg (rs36064369) and as lung-specific *cis*-eQTL in GTEx (rs116612101) (**Supplementary Table 12**).

To assess whether heritability of ND is enriched in regions surrounding genes with the highest specific gene expression patterns in given tissue/cell type(s), we applied LDSC-SEG ^44^ using the EUR-specific ND GWAS meta-analysis results with reference to 205 tissues/cell types with publicly available gene expression data assembled from GTEx^45^ (53 human tissues/cell types) and the underlying data that is used to comprise the Data-driven Expression Prioritized Integration for Complex Traits (DEPICT) tool ^46,47^ (152 tissues/cell types from humans and rodent models). We observed statistically significant enrichment in one tissue (cerebellum) at Bonferroni-corrected P<2.4×10^−4^ (**Supplementary Table 13**), indicating that genes spanning ND-associated SNPs are enriched for specific expression in the cerebellum relative to other tissues/cell types.

## Discussion

We expanded current knowledge of ND in this largest GWAS to date, by identifying two novel genome-wide significant loci as well as 3 known loci, extending associations of additional loci implicated for other smoking phenotypes, and detecting significant genetic correlations of ND with 13 other complex phenotypes and with gene expression in cerebellum. The top novel SNPs between *MAGI2* and *GNAI1* (chr7q21) and in *TENM2* (chr5q34) were independent of previously reported GWAS signals for any smoking trait. Three of our genome-wide significant loci were known: (1) *CHRNA5-CHRNA3-CHRNB4* (chr15q25) is irrefutably associated with ND, as driven largely by CPD^6^ (2) Our initial GWAS meta-analysis of 5 studies (now part of the iNDiGO consortium)^21^ identified *CHRNA4* (chr20q13) at genome-wide significance. Subsequent associations were found with heavy vs. never smoking in the UK Biobank^48^ and with initiation, CPD, and cessation in GSCAN. ^5^ (3) *DBH* (chr9q34) was first identified as genome-wide significant for smoking cessation but later associated with ND in our meta-analysis of 15 studies (now part of the iNDiGO consortium)^22^ and with CPD and cessation in GSCAN. ^5^ Known loci were corroborated at the gene level with aggregated single SNP associations that take physical proximity and chromatin interactions or *cis*-eQTL evidence into account.

The novel ND-associated locus with lead SNP rs2714700 is intergenic between *MAGI2* (membrane associated guanylate kinase, WW and PDZ domain containing 2) and *GNAI1* (G protein subunit alpha i1). We identified rs2714700 at genome-wide significance for its association with ND, which was driven by CPD (unlike rs1862416), TTFC, and other FTND items, indicating that this SNP association may reflect both primary and secondary features of ND. Primary (or core) features of ND are necessary and sufficient for habit formation (heaviness of smoking [tolerance], automaticity, loss of control, and craving), while secondary features of ND underlie smoking that is goal based, e.g., relief of negative mood or cognitive enhancement^49-52^. Rs2714700 was also associated with HSI in the independent UK Biobank. The *cis*-eQTL evidence for rs2714700 in the hippocampus suggests that it may influence expression of the long noncoding RNA *MAGI2-AS3* (MAGI2 antisense RNA 3). *MAGI2-AS3* has been mainly studied for its role in the progression of cancer, including glioma in the brain.^53^ No genome-wide significant associations have been reported within 1MB of rs2714700 in the GWAS catalog. Our evidence of genome-wide significance for rs2714700 points to a novel locus that has not been associated with smoking or any related trait, and its functional relevance merits further investigation.

We also observed a genome-wide significant association of ND with rs1862416, a lung-specific *cis*-eQTL for *TENM2. TENM2* encodes teneurin transmembrane protein 2, a cell surface receptor that plays a fundamental role in neuronal connectivity and synaptogenesis.^54^ With rs1862416 residing in the promoter of *CTB-77H17*.*1*, it could influence this antisense RNA, which in turn could dysregulate its sense transcript, *TENM2*. As an illustrative example, the SNP rs4307059, identified at genome-wide significance and independently replicated for autism^55^ is annotated to and acts as a promoter region *cis*-eQTL for the antisense RNA *MSNP1AS* (moesin pseudogene 1, antisense) that influences regulation of its sense transcript, *MSN*.^56^ However, while rs1862416 is generally indicated for its potential regulatory role (i.e., enhancer and promoter annotations and *cis*-eQTL evidence), its specific effect on either *CTB-77H17*.*1* or *TENM2* regulation in brain tissue was not evident in currently available data.

Further, independent association testing using HSI in the UK Biobank did not yield statistical significance for rs1862416. Similarly, the gene-based associations for the novel loci were not corroborated in UK Biobank. These differences in observed SNP- and gene-based associations may reflect components of ND that are not fully captured by the two FTND items that comprise the HSI (TTFC and CPD), as suggested by the specific FTND item association testing among the iNDiGO studies. Rs1862416 was suggestively associated (P<0.05) with TTFC, “Which cigarette would you hate most to give up?” (the first one in the morning vs. all others), and “Do/did you smoke if you are so ill that you are in bed most of the day?” (yes/no). These item responses reflect withdrawal symptoms that are indicative of secondary features of ND, as compared with primary features of ND associated with habit formation. ^49-52^ Having the composite ND phenotype may have enhanced our power for discovering *TENM2*, but its detection in the UK Biobank may have been limited by the reliance on the HSI.

Beyond our discovery of rs1862416 with ND, SNPs across the *TENM2* gene have been identified at genome-wide significance, as presented in the GWAS catalog^57^, for educational attainment,^38^ smoking initiation (ever vs. never smoking),^5-58-60^age of smoking initiation,^5^ smoking cessation (current vs. former smoking),^5^ cigarette pack-years,^61^ alc^60^number of sexual partners,^58^ depression,^63,64^ risk taking tendency,^58^ body mass index,^60^ menarche (age at onset) ^65^, and regular attendance at a religious group^66^. Our pairwise comparisons supported pleiotropic associations in the *TENM2* region. At the variant level, all *TENM2* SNPs in the GWAS catalog have very low r^2^ values with our novel SNP, rs1862416 (**Supplementary Figure 6**), and our conditional modeling results showed that rs1862416 was associated with ND independently from other *TENM2* SNPs implicated in GSCAN. While rs1862416 may have an ND-specific effect, the *TENM2* region has pleiotropic effects on ND, traits that are genetically correlated with ND, and other traits.

The genetics of smoking behaviors, more broadly, has rapidly evolved with the GSCAN consortium having amassed a very large sample size and identified 298 genome-wide significant loci for smoking traits representing single components: ever vs. never smoking, age of smoking initiation, CPD, and current vs. former smoking.^5^ We observed statistically significant genetic correlations of each of these smoking traits with ND (highest r_g_ =0.95, as observed between ND and CPD), yet our two novel ND-associated loci were not identified at genome-wide significance by GSCAN (smallest P=0.033 for rs1862416-T; smallest P=0.016 for rs2714700-T), suggesting that these loci are specific to ND. Similarly, the majority of GSCAN-identified loci were trait-specific (191 of the 298 loci), where the other 107 loci were pleiotropic with associations identified for two or more of the smoking traits^5^ In our evaluation of GSCAN-identified loci, we corroborated associations of several previously implicated loci for ND (e.g., nicotine acetylcholine receptors genes *CHRNA5-A3-B4* and *CHRNA4*) and three additional loci (*DRD2, C16orf97*, and *CHRNB2*) that have not been reported in prior ND GWAS. Of these loci, *DRD2* is notable as a long-studied addiction candidate gene4 and its recent identification as genome-wide significant for alcohol use disorder for rs493627767, which is correlated (r ^2^ =0.94 in 1000G EUR, 0.82 in 1000G AFR) with rs7125588, the top SNP identified for CPD in GSCAN and associated with ND in iNDiGO; these results support a shared genetic effect of *DRD2* underlying addiction. Notably, rs7125588 is not correlated (r2^2^ =0.04 in 1000G EUR, 0.01 in 1000G AFR) with the *DRD2* variant rs1800497 (historically referred to as the ‘Taq1A’ polymorphism), which is not significantly associated with ND in iNDiGO (P=0.24).

Other GSCAN loci were detected for the single component smoking traits but show no evidence for association in our study (**Supplementary Table 14**), suggesting that these loci influence stages of smoking other than ND, or they exert weak effects on ND that we were underpowered to detect. We expect that additional GSCAN-identified loci are associated with ND, but their detection will require a larger sample size. These results demonstrate the utility of studying the genetics of the composite ND phenotype and comparing with GWAS of other smoking traits to tease apart loci that are specific to one stage (i.e., initiation, regular smoking, ND, cessation) vs. loci that influence multiple stages to better understand the full spectrum of smoking behaviors.

Beyond the smoking traits, we observed significant genetic correlations between ND and alcohol dependence^30^, years of schooling^38^, neuroticism^31^, comorbid psychiatric traits (attention deficit hyperactivity disorder^32^, bipolar disorder ^33^, major depression ^34^, schizophrenia ^35^, and posttraumatic stress disorder^68^), and smoking-related health consequences (lung cancer36 and coronary artery disease^37^). Some of these observations corroborate prior findings (for example, alcohol dependence^30^ and schizophrenia ^69,70^ with ND), whereas the other correlations extend to ND prior observations for the single component smoking traits (for example, CPD with years of schooling ^5^, neuroticism^5^, major depression^5^, coronary artery disease^5^, and lung cancer^36^). The genetic correlation between ND and gene expression in cerebellum is a notable observation consistent with cerebellum-specific *cis*-eQTL effects observed for the ND-associated *DNMT3B* SNP rs910083^22^ and the age of smoking initiation-associated *CHRNA2* SNP rs11780471^36^, both of which are also associated with lung cancer. These findings add to the evidence that the cerebellum may be important for ND risk, ^71,72^ in addition to the other addiction-relevant brain tissues. However, since the cerebellum contains a higher neuronal concentration than other brain tissues^44,73^ future studies are needed to decipher whether the cerebellar gene regulatory effects in the etiology of ND are due to neuronal activity. Additionally, although genetic correlation between ND and another trait suggest shared genetics underlying the phenotypes, multiple mechanisms can produce significant correlations (i.e., unmeasured intermediary phenotypes, correlated risk variants, mediation).^74-76^ Identifying the true mechanistic explanation requires further investigations.

The present ND GWAS meta-analysis follows two prior waves of data assembly by the iNDiGO consortium (Ns=17,074^21^, 38,602 ^22^, and now 58,000) and is the largest to date for the field. Despite still having substantially smaller sample sizes than the GSCAN GWAS, at each wave, increasing sample size for diverse ancestry groups (EURs and AAs) has illuminated ND-associated loci, some of which are shared with other stages of smoking while others are specific to ND. Our present findings underscore the complexity even within the ND phenotype, as our novel loci displayed patterns of association with specific FTND items that reflect primary or secondary ND features, e.g., the *TENM2* SNP influenced secondary features that are not captured simply by heaviness of smoking. Future studies are needed to further dissect the genetic architectures underlying each of the specific FTND items. Understanding genetic similarities and differences that underlie these items and their contributions to primary vs. secondary ND may better inform treatment strategies, e.g., changing environmental cues for individuals whose smoking is driven solely by primary ND features vs. treating withdrawal for individuals whose ND is augmented with secondary features.51 Studying the genetics of ND alongside other smoking traits (e.g., initiation and cessation) is key to gaining a better understanding of the neurobiological perturbations that influence the trajectory of smoking behaviors and their treatment implications.

## Methods

We assembled 58,000 participants from 23 iNDiGO consortium studies with genome-wide SNP genotypes and FTND phenotype data available for ever smokers to perform ND GWAS meta-analyses. Fifteen of the studies were included from our prior GWAS using their original or updated sample sizes (total N increased from 38,602^22^ to 46,098 in the current analysis), while 8 studies were added for the current study (total N=11,902). Participant characteristics are provided in **Supplementary Table 3**, and details of the study designs, genotyping, quality control, imputation, and statistical analyses are provided in the **Supplementary Methods**. Institutional review boards at the respective sites approved the study protocols, and all participants provided written informed consent.

### ND GWAS meta-analysis

The FTND is a well-validated, widely used 6-item questionnaire that assesses psychologic dependence on nicotine, with scores ranging from 0 (no dependence) to 10 (highest dependence level).^20,21^ As done before^21,22^we categorized FTND scores as mild (scores 0–3), moderate (scores 4–6), or severe (scores 7–10). FTND data reflected current smoking behaviors at the time of interview (i.e., current FTND) or the period of heaviest smoking among ever smokers (i.e., lifetime FTND). We previously found only small differences in genetic association results due to any measurement variance when using current vs. lifetime FTND. ^77^ See **Supplementary Methods** for further details on the ND phenotype data by study.

For each study, genome-wide SNP/indel associations with the 3-level categorical ND outcome were tested within an ancestry group using linear regression. Covariates included age, sex, principal component eigenvectors, and study-specific covariates where warranted. For studies that included relatives, relatedness was accounted for in the regression modeling. See the **Supplementary Methods** for additional study-specific details.

GWAS results were combined using fixed-effect inverse variance-weighted meta-analyses in METAL.^78^ Prior to performing meta-analyses, we applied genomic control to results from one study, deCODE, to adjust for inflation due to relatedness among participants (λ=1.12); all other studies had low inflation (λ=0.99–1.04) (**Supplementary Table 3**). We removed SNPs/indels with minor allele frequency (MAF) <1% in the 1000G phase 3 reference panel for the analyzed ancestry group (1000G European or African superpopulations), imputation info score<0.3, or availability in only one study. All variant annotations correspond to the National Center for Biotechnology Information (NCBI) build 37. As before ^22^, the threshold of genome-wide significance was set at P = 5×10 ^-8^. Regional association plots of novel genome-wide significant loci were constructed using LocusZoom^79^ with references of either 1000G European or African panels to estimate linkage disequilibrium of the lead SNP (based on smallest meta-analysis P-value) and surrounding SNPs. The lead SNP for each novel FTND locus was tested for association with each of the specific FTND items (**Supplementary Methods**).

For any ND-associated SNPs located within the bounds of loci identified by GSCAN (1 MB surrounding the lead SNP),^5^ conditional models were analyzed using our GWAS summary statistics and the Genome-wide Complex Trait Analysis (GCTA) tool, adjusted for the lead SNPs in GSCAN.^40,41^ To contextualize the magnitude of the observed effect sizes, we calculated odds ratios (ORs) using the β estimate from the single SNP linear regression model (OR=exp[2×βSNP] for severe vs. mild ND, with OR>1 corresponding to an increased risk of severe ND) and compared these values across studies and ancestries using the Forest Plot Viewer.^80^

### Independent testing using heaviness of smoking index in the UK Biobank

Novel, genome-wide significant SNPs from our ND GWAS meta-analysis were tested in the UK Biobank. Although all 6 items of the FTND were not collected in the UK Biobank, two items (CPD and TTFC) were collected among current smokers. These two items together form the HSI, which is highly correlated with the full-scale FTND.^26^ We derived HSI scores, ranging from 0 (no dependence) to 6 (highest dependence level), and categorized them as follows: mild (scores 0–2), moderate (scores 3–4), and severe (scores 5–6). These HSI categories were highly concordant (89.3%) with our routinely used FTND categories using the COGEND study, which was ascertained specifically for ND (**Supplementary Methods** and **Supplementary Table 15**). The final analysis dataset included 33,791 current smokers (18,063 mildly, 13,395 moderately, and 2,333 severely dependent, as defined by HSI). Our linear regression model included covariates for age, sex, and principal component eigenvectors (**Supplementary Methods**).

### Gene-based association testing

To assess evidence for association beyond single variants, we applied two methods that aggregate SNP-based summary statistics at the gene level. For genome-wide testing with both methods, we used the EUR-specific GWAS meta-analysis results from iNDiGO as the input dataset, given the reliance on linkage disequilibrium (LD) reference data by ancestry in calculating the gene-based summary statistics. First, H-MAGMA^27^ computes gene-based association statistics by aggregating SNP associations based on physical proximity to the target gene(s) measured by chromatin interaction maps from human brain tissue. We included SNPs with an rs identification number (9,525,836 SNPs) and coupled them with Hi-C reference datasets from fetal^81^ and adult brain tissues, specifically cortical tissues,^82^ that are available for running H-MAGMA. H-MAGMA converted SNP-level p-values into gene-level p-values. We identified statistically significant genes that were associated with ND at Bonferroni-corrected threshold of P<2.7×10 ^-6^ (α=0.05/18,655 protein coding genes).

Second, we applied Summary-MultiXcan (S-MultiXcan)^28^ to compute gene-level associations by leveraging imputed genetically driven gene expression using RNA-Seq across the 13 adult brain tissues in GTEx as reference data. S-MultiXcan^28^, an extension of the S-PrediXcan method for integrating eQTLs with GWAS summary statistics^83^, aggregates eQTL information across multiple tissue types to enhance statistical power, while still presenting the single tissue with the best evidence for association. We applied Bonferroni correction to declare statistically significant gene-based associations as P<3.5×10^−6^ (α=0.05/14,494 genes).

For both gene-based methods, we carried forward significant gene-level associations and tested them in the UK Biobank, using HSI as a proxy for ND, as done with the single SNP associations.

### Genetic correlations of ND with other complex phenotypes and with gene expression

Summary statistics from the EUR-specific meta-analyses were used as input into LD score regression (LDSC)^29^ with reference to the 1000G EUR panel to estimate the SNP heritability 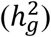 of ND and its genetic correlations with 46 other complex phenotypes, including other smoking, drug, and alcohol use and dependence traits, smoking-related health consequences (e.g., cancer, COPD, and coronary heart disease), psychiatric and neurologic disorders, cognitive and educational traits, and brain volume metrics. The full list of phenotypes and GWAS datasets, as obtained from LD Hub^84^ or shared by the original study investigators, are provided in **Supplementary Table 11**.

Similarly, EUR-specific GWAS meta-analysis summary statistics were input into stratified LDSC, as applied to specifically expressed genes (LDSC-SEG),^44^ with reference to 205 tissues and cell types from two sources—RNA-sequencing data on 53 human tissues/cell types in GTEx^85^ and array-based data on 152 tissues/cell types from humans and rodent models that underlie the DEPICT tool and made available in Gene Expression Omnibus.^46,47^ See full list of the 205 tissues/cell types in **Supplementary Table 13**. Similarly to the initial application of LDSC-SEG,^44^ these two sources were selected because their expression data included a wide range of ND-relevant and other tissues and cell types in humans, as opposed to focused information on a particular tissue. LDSC-SEG involved comparing expression of each gene in each tissue/cell type with that in other tissues/cell types, selecting the top 10% of differentially expressed genes, annotating SNPs from the GWAS summary statistics that lie within 100kb windows of the selected genes, and using the stratified LDSC method to estimate the enrichment in SNP heritability for ND for the given gene set compared to the baseline LDSC model with all genes. For each analysis, a Bonferroni correction was applied to assess statistical significance: P<0.0011 (α=0.05/46 phenotypes) for LDSC and P<2.4×10-4 (α=0.05/205 tissues/cell types) for LDSC-SEG.

### Associations of ND loci with other complex traits

We applied pairwise GWAS (GWAS-PW v0.21 [github.com/joepickrell/gwas-pw/])^39^ to characterize the cross-phenotype associations for ND and its genetically correlated phenotypes, as revealed in the LDSC analyses. Specifically, we applied GWAS-PW to the “Cigarette smoking”, “Drug and alcohol use”, “Personality”, and “Psychiatric” phenotypes with significant genetic correlation with ND. Using EUR-specific GWAS summary statistics for ND and its correlated phenotypes, for each pairwise comparison of ND to a given phenotype, we calculated a correlation statistic that is used by GWAS-PW to account for potential sample overlaps between studies. We followed the approach as detailed in Pickrell et al.^39^ We then reduced the SNP set to only SNPs with summary statistics available from both studies and that also were located within an LD block for any of the 5 FTND GWAS significant loci. We defined the LD blocks by using LDproxy^86^ with the top (i.e., most significant) SNP from each FTND-associated locus and extracting r^2^ values (based on 1000 Genomes Phase 3 EUR populations) for all SNPs within 0.5 Mb of the top SNP. The minimum and maximum genomic coordinate for all extracted SNPs with r^2^ > 0.2 were used as the LD block boundaries.

### cis-eQTL assessment of novel ND-associated SNPs

For each novel locus, we identified a credible set, or the set of SNPs most likely to contain the causal variant, using a Bayesian method^87^ implemented via LocusZoom.^79^ To assess evidence for SNP-gene associations, SNPs in the credible set were queried against GTEx (version 8) *cis*-expression quantitative trait loci (*cis*-eQTL) results derived from SNP genotype and RNA-sequencing data across 44 tissues (N=126–209 for the 13 brain tissues).^85^ The GTEx portal (https://gtexportal.org/home/) presents significant single-tissue *cis*-eQTLs, based on a false discovery rate (FDR) <5%.

We also assessed single-tissue *cis*-eQTL evidence from the BrainSeq consortium that includes larger sample sizes with SNP genotype and RNA-sequencing data available in two brain tissues, dorsolateral prefrontal cortex (N=453) and hippocampus (N=447).^42^ Of the 551 individuals with data available in at least one brain tissue, 286 were schizophrenia cases; case/control status was included as a covariate for adjustment in the *cis*-eQTL analysis, as described elsewhere.88 Significant *cis*-eQTLs at FDR <10% are available at http://eqtl.brainseq.org/phase2/eqtl/.

## Data Availability

The prior meta-analysis summary statistics^22^ are available via dbGaP accession number phs001532.v1.p1. The summary statistics generated from the current study are included under version 2 of this dbGaP study (accession number phs001532.v2.p1), or are available upon request to the corresponding author (D.B.H.). Individual-level genotype and phenotype data for many of the contributing cohorts are also available via dbGaP, as outlined in the study descriptions in the Supplementary Information.

## Acknowledgements

We are grateful to the many study participants, who made this work possible. The meta-analysis was supported by the National Institute on Drug Abuse grant numbers R01 DA042090 (PI: DBH) and R01 DA036583 (PI: LJB). The authors thank deCODE Genetics / Amgen and its investigators (Gunnar W. Reginsson, Thorgeir E. Thorgeirsson, and Kari Stefansson) for contributing and analyzing their data for inclusion in the meta-analysis. Their work was supported in part by NIDA R01 DA017932 (PI: Kari Stefansson). Acknowledgments for all other ND studies, contributed by the authors and/or made publicly available, are included in the Supplementary Information. This research also leveraged the UK Biobank Resource under Application Number 24603.

## Author Contributions

Contributions of each author are categorized using terms from the Contributor Roles Taxonomy (https://casrai.org/credit/). Conceptualization: D.B.H., D.W.M., E.O.J., L.J.B., M.J.B., M.R., N.E.C., T.B.B.; Data curation: B.C.Q., C.C.M., F.A., J.H., J.K., M.N., N.C.G., N.E.C., N.L.S., R.G., S.Z., Y.G.; Formal analysis: A.W., B.C.Q., C.A.M., C.C.M., D.D., F.A., H.Y., J.A.M., J.H., M.J.B., M.N., M.U.C., N.C.G., R.S., T.P., Y.G.; Funding acquisition: D.B.H., D.M.D., D.W.M., E.O.J., J.K., L.J.B., M.R., W.G.I.; Investigation: D.I.B., E.O.J., F.A., G.W., H.R.K., J.G., J.H., J.K., J.M.V., L.A.F., L.J.B., M.C.N., M.J.B., M.M., M.R., M.T.L., N.L.S., P.M., S.V., W.G.I.; Methodology: D.W.M.; Project administration: D.B.H., D.M.D., D.W.M., J.K.; Resources: D.B.H., D.I.B., D.M.D., D.W.M., G.W., J.G., J.K., J.M.V., K.A.Y., L.A.F., L.J.B., M.C.N., M.J.B., M.L.M., M.M., M.R., M.T.L., N.C.G., N.E.C., P.M., R.G., S.V., W.G.I.; Software: A.W., B.C.Q., H.W., J.A.M., M.U.C., N.C.G., N.Y.A.S.; Supervision: D.B.H., D.M.D., D.W.M., E.O.J., J.K., M.J.B., M.N.; Validation: D.B.H., M.L.; Visualization: B.C.Q., D.B.H., J.A.M., M.J.B.; Writing – original draft: B.C.Q., D.B.H., J.A.M., S.Z., T.B.B.; Writing – review & editing: A.W., B.C.Q., C.A.M., C.C.M., D.B.H., D.I.B., D.M.D., D.W.M., E.O.J., F.A., G.W., H.R.K., J.E.H., J.G., J.K., J.M.V., K.A.Y., L.A.F., L.J.B., M.C.N., M.J.B., M.L.M., M.M., M.R., M.T.L., N.C.G., N.E.C., N.L.S., P.M., S.V., T.B.B.

## References

1. World Health Organization. WHO report on the global tobacco epidemic, 2017: monitoring tobacco use and prevention policies. (Geneva, 2017).

2. U.S. Department of Health and Human Services. The Health Consequences of Smoking-50 Years of Progress: A Report of the Surgeon General, (Atlanta (GA), 2014).

3. Sullivan, P.F. & Kendler, K.S. The genetic epidemiology of smoking. Nicotine Tob Res 1 Suppl 2, S51–7; discussion S69-70 (1999).

4. Agrawal, A. et al. The genetics of addiction-a translational perspective. Transl Psychiatry 2, e140 (2012).

5. Liu, M. et al. Association studies of up to 1.2 million individuals yield new insights into the genetic etiology of tobacco and alcohol use. Nat Genet 51, 237–244 (2019).

6. Hancock, D.B., Markunas, C.A., Bierut, L.J. & Johnson, E.O. Human Genetics of Addiction: New Insights and Future Directions. Curr Psychiatry Rep 20, 8 (2018).

7. Baker, T.B. et al. Are tobacco dependence and withdrawal related amongst heavy smokers? Relevance to conceptualizations of dependence. J Abnorm Psychol 121, 909–21 (2012).

8. Zelman, D.C., Brandon, T.H., Jorenby, D.E. & Baker, T.B. Measures of affect and nicotine dependence predict differential response to smoking cessation treatments. J Consult Clin Psychol 60, 943–52 (1992).

9. Gu, F. et al. Time to smoke first morning cigarette and lung cancer in a case-control study. J Natl Cancer Inst 106, dju118 (2014).

10. Guertin, K.A. et al. Time to First Morning Cigarette and Risk of Chronic Obstructive Pulmonary Disease: Smokers in the PLCO Cancer Screening Trial. PLoS One 10, e0125973 (2015).

11. Fagerstrom, K. Determinants of tobacco use and renaming the FTND to the Fagerstrom Test for Cigarette Dependence. Nicotine Tob Res 14, 75–8 (2012).

12. Heatherton, T.F., Kozlowski, L.T., Frecker, R.C. & Fagerstrom, K.O. The Fagerstrom Test for Nicotine Dependence: a revision of the Fagerstrom Tolerance Questionnaire. Br J Addict 86, 1119–27 (1991).

13. Breslau, N. & Johnson, E.O. Predicting smoking cessation and major depression in nicotine-dependent smokers. Am J Public Health 90, 1122–7 (2000).

14. Agrawal, A. et al. A latent class analysis of DSM-IV and Fagerstrom (FTND) criteria for nicotine dependence. Nicotine Tob Res 13, 972–81 (2011).

15. Paik, S.H. et al. Prevalence and analysis of tobacco use disorder in patients diagnosed with lung cancer. PLoS One 14, e0220127 (2019).

16. Baker, T.B. et al. Time to first cigarette in the morning as an index of ability to quit smoking: implications for nicotine dependence. Nicotine Tob Res 9 Suppl 4, S555–70 (2007).

17. Sweitzer, M.M., Denlinger, R.L. & Donny, E.C. Dependence and withdrawal-induced craving predict abstinence in an incentive-based model of smoking relapse. Nicotine Tob Res 15, 36–43 (2013).

18. Bolt, D.M. et al. The Wisconsin Predicting Patients’ Relapse questionnaire. Nicotine Tob Res 11, 481–92 (2009).

19. Haberstick, B.C. et al. Genes, time to first cigarette and nicotine dependence in a general population sample of young adults. Addiction 102, 655–65 (2007).

20. Conway, K.P. et al. Data compatibility in the addiction sciences: an examination of measure commonality. Drug Alcohol Depend 141, 153–8 (2014).

21. Hancock, D.B. et al. Genome-wide meta-analysis reveals common splice site acceptor variant in CHRNA4 associated with nicotine dependence. Transl Psychiatry 5, e651 (2015).

22. Hancock, D.B. et al. Genome-wide association study across European and African American ancestries identifies a SNP in DNMT3B contributing to nicotine dependence. Mol Psychiatry 23, 1911–1919 (2018).

23. Thorgeirsson, T.E. et al. A variant associated with nicotine dependence, lung cancer and peripheral arterial disease. Nature 452, 638–42 (2008).

24. Bierut, L.J. et al. Variants in nicotinic receptors and risk for nicotine dependence. Am J Psychiatry 165, 1163–71 (2008).

25. Huedo-Medina, T.B., Sanchez-Meca, J., Marin-Martinez, F. & Botella, J. Assessing heterogeneity in meta-analysis: Q statistic or I2 index? Psychol Methods 11, 193–206 (2006).

26. DiFranza, J.R. et al. What aspect of dependence does the fagerstrom test for nicotine dependence measure? ISRN Addict 2013, 906276 (2013).

27. Sey, N.Y.A. et al. A computational tool (H-MAGMA) for improved prediction of brain-disorder risk genes by incorporating brain chromatin interaction profiles. Nat Neurosci (2020).

28. Barbeira, A.N. et al. Integrating predicted transcriptome from multiple tissues improves association detection. PLoS Genet 15, e1007889 (2019).

29. Bulik-Sullivan, B. et al. An atlas of genetic correlations across human diseases and traits. Nat Genet 47, 1236–41 (2015).

30. Walters, R.K. et al. Transancestral GWAS of alcohol dependence reveals common genetic underpinnings with psychiatric disorders. Nat Neurosci 21, 1656–1669 (2018).

31. Okbay, A. et al. Genetic variants associated with subjective well-being, depressive symptoms, and neuroticism identified through genome-wide analyses. Nat Genet 48, 624–33 (2016).

32. Demontis, D. et al. Discovery of the first genome-wide significant risk loci for attention deficit/hyperactivity disorder. Nat Genet 51, 63–75 (2019).

33. Stahl, E.A. et al. Genome-wide association study identifies 30 loci associated with bipolar disorder. Nat Genet 51, 793–803 (2019).

34. Howard, D.M. et al. Genome-wide meta-analysis of depression identifies 102 independent variants and highlights the importance of the prefrontal brain regions. Nat Neurosci 22, 343–352 (2019).

35. Schizophrenia Working Group of the Psychiatric Genomics, C. Biological insights from 108 schizophrenia-associated genetic loci. Nature 511, 421–7 (2014).

36. McKay, J.D. et al. Large-scale association analysis identifies new lung cancer susceptibility loci and heterogeneity in genetic susceptibility across histological subtypes. Nat Genet 49, 1126–1132 (2017).

37. Nikpay, M. et al. A comprehensive 1,000 Genomes-based genome-wide association meta-analysis of coronary artery disease. Nat Genet 47, 1121–1130 (2015).

38. Lee, J.J. et al. Gene discovery and polygenic prediction from a genome-wide association study of educational attainment in 1.1 million individuals. Nat Genet 50, 1112–1121 (2018).

39. Pickrell, J.K. et al. Detection and interpretation of shared genetic influences on 42 human traits. Nat Genet 48, 709–17 (2016).

40. Yang, J., Lee, S.H., Goddard, M.E. & Visscher, P.M. GCTA: a tool for genome-wide complex trait analysis. Am J Hum Genet 88, 76–82 (2011).

41. Yang, J. et al. Conditional and joint multiple-SNP analysis of GWAS summary statistics identifies additional variants influencing complex traits. Nat Genet 44, 369–75, S1-3 (2012).

42. BrainSeq: A Human Brain Genomics Consortium. BrainSeq: Neurogenomics to Drive Novel Target Discovery for Neuropsychiatric Disorders. Neuron 88, 1078–1083 (2015).

43. Ward, L.D. & Kellis, M. HaploReg: a resource for exploring chromatin states, conservation, and regulatory motif alterations within sets of genetically linked variants. Nucleic Acids Res 40, D930–4 (2012).

44. Finucane, H.K. et al. Heritability enrichment of specifically expressed genes identifies disease-relevant tissues and cell types. Nat Genet 50, 621–629 (2018).

45. GTEx Consortium. Human genomics. The Genotype-Tissue Expression (GTEx) pilot analysis: multitissue gene regulation in humans. Science 348, 648–60 (2015).

46. Pers, T.H. et al. Biological interpretation of genome-wide association studies using predicted gene functions. Nat Commun 6, 5890 (2015).

47. Fehrmann, R.S. et al. Gene expression analysis identifies global gene dosage sensitivity in cancer. Nat Genet 47, 115–25 (2015).

48. Wain, L.V. et al. Novel insights into the genetics of smoking behaviour, lung function, and chronic obstructive pulmonary disease (UK BiLEVE): a genetic association study in UK Biobank. Lancet Respir Med 3, 769–781 (2015).

49. Piper, M.E. et al. Refining the tobacco dependence phenotype using the Wisconsin Inventory of Smoking Dependence Motives. J Abnorm Psychol 117, 747–61 (2008).

50. Piasecki, T.M., Piper, M.E. & Baker, T.B. Refining the tobacco dependence phenotype using the Wisconsin Inventory of Smoking Dependence Motives: II. Evidence from a laboratory self-administration assay. J Abnorm Psychol 119, 513–23 (2010).

51. Piasecki, T.M., Piper, M.E. & Baker, T.B. Tobacco Dependence: Insights from Investigations of Self-Reported Smoking Motives. Curr Dir Psychol Sci 19, 395–401 (2010).

52. Piasecki, T.M., Piper, M.E., Baker, T.B. & Hunt-Carter, E.E. WISDM primary and secondary dependence motives: associations with self-monitored motives for smoking in two college samples. Drug Alcohol Depend 114, 207–16 (2011).

53. Chen, X.D., Zhu, M.X. & Wang, S.J. Expression of long non-coding RNA MAGI2AS3 in human gliomas and its prognostic significance. Eur Rev Med Pharmacol Sci 23, 3455–3460 (2019).

54. Silva, J.P. et al. Latrophilin 1 and its endogenous ligand Lasso/teneurin-2 form a high-affinity transsynaptic receptor pair with signaling capabilities. Proc Natl Acad Sci U S A 108, 12113–8 (2011).

55. Wang, K. et al. Common genetic variants on 5p14.1 associate with autism spectrum disorders. Nature 459, 528–33 (2009).

56. Kerin, T. et al. A noncoding RNA antisense to moesin at 5p14.1 in autism. Sci Transl Med 4, 128ra40 (2012).

57. Welter, D. et al. The NHGRI GWAS Catalog, a curated resource of SNP-trait associations. Nucleic Acids Res 42, D1001–6 (2014).

58. Karlsson Linner, R. et al. Genome-wide association analyses of risk tolerance and risky behaviors in over 1 million individuals identify hundreds of loci and shared genetic influences. Nat Genet 51, 245–257 (2019).

59. Erzurumluoglu, A.M. et al. Meta-analysis of up to 622,409 individuals identifies 40 novel smoking behaviour associated genetic loci. Mol Psychiatry (2019).

60. Kichaev, G. et al. Leveraging Polygenic Functional Enrichment to Improve GWAS Power. Am J Hum Genet 104, 65–75 (2019).

61. Buchwald, J. et al. Genome-wide association meta-analysis of nicotine metabolism and cigarette consumption measures in smokers of European descent. Mol Psychiatry (2020).

62. Lutz, S.M. et al. A genome-wide association study identifies risk loci for spirometric measures among smokers of European and African ancestry. BMC Genet 16, 138 (2015).

63. Nagel, M. et al. Meta-analysis of genome-wide association studies for neuroticism in 449,484 individuals identifies novel genetic loci and pathways. Nat Genet 50, 920–927 (2018).

64. Wray, N.R. et al. Genome-wide association analyses identify 44 risk variants and refine the genetic architecture of major depression. Nat Genet 50, 668–681 (2018).

65. Perry, J.R. et al. Parent-of-origin-specific allelic associations among 106 genomic loci for age at menarche. Nature 514, 92–97 (2014).

66. Day, F.R., Ong, K.K. & Perry, J.R.B. Elucidating the genetic basis of social interaction and isolation. Nat Commun 9, 2457 (2018).

67. Kranzler, H.R. et al. Genome-wide association study of alcohol consumption and use disorder in 274,424 individuals from multiple populations. Nat Commun 10, 1499 (2019).

68. Nievergelt, C.M. et al. International meta-analysis of PTSD genome-wide association studies identifies sex- and ancestry-specific genetic risk loci. Nat Commun 10, 4558 (2019).

69. Reginsson, G.W. et al. Polygenic risk scores for schizophrenia and bipolar disorder associate with addiction. Addict Biol (2017).

70. Hartz, S.M. et al. Genetic correlation between smoking behaviors and schizophrenia. Schizophr Res (2017).

71. Moulton, E.A., Elman, I., Becerra, L.R., Goldstein, R.Z. & Borsook, D. The cerebellum and addiction: insights gained from neuroimaging research. Addict Biol 19, 317–31 (2014).

72. Miquel, M. et al. Have we been ignoring the elephant in the room? Seven arguments for considering the cerebellum as part of addiction circuitry. Neurosci Biobehav Rev 60, 1–11 (2016).

73. Herculano-Houzel, S. & Lent, R. Isotropic fractionator: a simple, rapid method for the quantification of total cell and neuron numbers in the brain. J Neurosci 25, 2518–21 (2005).

74. Timofeeva, M.N. et al. Genetic polymorphisms in 15q25 and 19q13 loci, cotinine levels, and risk of lung cancer in EPIC. Cancer Epidemiol Biomarkers Prev 20, 2250–61 (2011).

75. Martin, J., Taylor, M.J. & Lichtenstein, P. Assessing the evidence for shared genetic risks across psychiatric disorders and traits. Psychol Med 48, 1759–1774 (2018).

76. Hjelmborg, J. et al. Lung cancer, genetic predisposition and smoking: the Nordic Twin Study of Cancer. Thorax 72, 1021–1027 (2017).

77. Glasheen, C. et al. Is the Fagerstrom test for nicotine dependence invariant across secular trends in smoking? A question for cross-birth cohort analysis of nicotine dependence. Drug Alcohol Depend 185, 127–132 (2018).

78. Willer, C.J., Li, Y. & Abecasis, G.R. METAL: fast and efficient meta-analysis of genomewide association scans. Bioinformatics 26, 2190–1 (2010).

79. Pruim, R.J. et al. LocusZoom: regional visualization of genome-wide association scan results. Bioinformatics 26, 2336–7 (2010).

80. Boyles, A.L., Harris, S.F., Rooney, A.A. & Thayer, K.A. Forest Plot Viewer: a new graphing tool. Epidemiology 22, 746–7 (2011).

81. Nowakowski, T.J. et al. Spatiotemporal gene expression trajectories reveal developmental hierarchies of the human cortex. Science 358, 1318–1323 (2017).

82. Wang, D. et al. Comprehensive functional genomic resource and integrative model for the human brain. Science 362(2018).

83. Barbeira, A.N. et al. Exploring the phenotypic consequences of tissue specific gene expression variation inferred from GWAS summary statistics. Nat Commun 9, 1825 (2018).

84. Zheng, J. et al. LD Hub: a centralized database and web interface to perform LD score regression that maximizes the potential of summary level GWAS data for SNP heritability and genetic correlation analysis. Bioinformatics 33, 272–279 (2017).

85. GTEx Consortium. et al. Genetic effects on gene expression across human tissues. Nature 550, 204–213 (2017).

86. Machiela, M.J. & Chanock, S.J. LDlink: a web-based application for exploring population-specific haplotype structure and linking correlated alleles of possible functional variants. Bioinformatics 31, 3555–7 (2015).

87. Wellcome Trust Case Control Consortium. et al. Bayesian refinement of association signals for 14 loci in 3 common diseases. Nat Genet 44, 1294–301 (2012).

88. Collado-Torres, L. et al. Regional Heterogeneity in Gene Expression, Regulation, and Coherence in the Frontal Cortex and Hippocampus across Development and Schizophrenia. Neuron 103, 203–216 e8 (2019).

