## Supplementary File for "Expanding the Genetic Architecture of Nicotine Dependence and its Shared Genetics with Multiple Traits: Findings from the Nicotine Dependence GenOmics (iNDiGO) Consortium"

**Supplementary Figure 6**. Linkage disequilibrium (LD) matrix of rs1862416 (marked by blue boxes) and other *TENM2* single nucleotide polymorphisms (SNPs) included in the genome-wide association study (GWAS) catalog[87] (https://www.ebi.ac.uk) for their genome-wide significant associations ($P<5\times10^{-8}$). r2 values, as obtained from LDlink51, correspond to the 1000 Genomes European panel. Numerical values correspond to the originally reported GWAS: 1, educational attainment[63]; 2a, smoking initiation (ever vs. never smoking)[48,88-90]; 2b, age of smoking initiation[48]; 2c, smoking cessation (current vs. former smoking)[48]; 2d, alcohol consumption (drinks per week)[48]; 3, lung function[90,91]; 4, height[90]; 5, number of sexual partners[88]; 6, depression[92,93]; 7, risk taking tendency[88]; 8, body mass index[90]; 9, menarche (age at onset)[94]; 10, cigarette pack-years[95]; and 11, regular attendance at a religious group[96]. rs11739827, associated with alcohol consumption[48], was not available for comparison with rs1862416 in LDlink...... 59

References ........ 61

### Supplementary Methods

#### Nicotine Dependence (ND) Study Descriptions, Quality Control (QC), and 1000 Genomes (1000G) Imputation

Our study included European (EUR) and African American (AA) ancestry ever smokers from 23 independent studies. Data were contributed by original study investigators or obtained via the database of Genotypes and Phenotypes (dbGaP). Fifteen of the studies (total N=38,062) were included in our prior genome-wide association study (GWAS) meta-analysis of ND.[1] The present study included the same sample size for 10 prior studies, an updated size for 5 of the prior studies, and 8 newly added studies. Additional details of each study are provided below and in **Supplementary Table 3**. As before,[1,2] ever smokers were defined by having reported smoking 100 or more cigarettes in their lifetime, unless otherwise stated, and ND was defined by the Fagerström Test for Nicotine Dependence (FTND).[3]

Our standard QC pipeline was applied to each study, unless otherwise stated. All studies were imputed to 1000G phase 3, except for deCODE as detailed below. Participants were removed due to missing rate >3%, sample duplication (identity-by-state >90%), first-degree relatedness (identity-by-descent >40%), gender discordance ($F_{st}$ <0.2 for chromosome X single nucleotide polymorphisms (SNPs) to confirm females and $F_{st}$ >0.8 to confirm males), excessive homozygosity ($F_{st}$ >0.5 or $F_{st}$ <-0.2), or chromosomal anomalies. SNPs were removed due to missing rate >3% or Hardy-Weinberg equilibrium (HWE) $P<1\times10^{-4}$. Genotyped SNPs passing QC were used as input for imputation with reference to 1000G phase 3 across all studies,[4] unless otherwise stated.

African American Nicotine Dependence (AAND). The community-based AAND study was designed to compare nicotine dependent smokers with smokers who never developed ND symptoms. Recruitment focused on AAs from the Chicago area between 2010 and 2013.

Participants reported smoking >100 cigarettes during their lifetime, and their ND was assessed using the FTND. Genotyping was performed on the Illumina Omni Express array. Following standard QC, the final analysis data set included 1,687 AAs with complete data on lifetime FTND (i.e., FTND based on when they reported smoking the most) and covariates—age, sex, and principal component (PC) eigenvectors. PC eigenvectors were computed to remove any residual bias due to population stratification.

Alcohol Dependence in African Americans: A Case-Control Genetic Study (ADAA). Data for the ADAA study were collected between 2009 and 2013. Alcohol dependent cases, who met Diagnostic and Statistical Manual of Mental Disorders, 4th. Edition (DSM-IV) criteria as assessed using a modified version of the Semi-Structured Assessment for the Genetics of Alcoholism, were recruited from treatment centers in St. Louis, Missouri. Alcohol dependent controls, who had consumed at least 12 alcohol beverages in their lifetime but did not meet DSM-IV criteria for alcohol abuse or dependence, were recruited from households in neighborhoods located in proximity to neighborhoods where the alcohol dependent cases resided. Participants were genotyped on a custom array that is based on an Illumina HumanOmniExpressExome background, and QC steps were applied with procedures that largely mimic our standard QC, excluding participants with call rate >1%, gender discrepancy, ancestry discrepancy, chromosomal anomalies, duplicate samples, or first-degree relatives and excluding SNPs with call rate >2%, no mapping, >2 discordant calls in duplicated samples, >5 discordant calls at the same position, or HWE $P<1\times10^{-4}$. The ND GWAS analysis included 1,145 current and former smokers, and covariate adjustments were made for age, sex, alcohol dependence (DSM-IV), cocaine dependence (DSM-IV), and PC eigenvectors.

Collaborative Genetic Study of Nicotine Dependence (COGEND and COGEND2). AAND was modeled after its predecessor, COGEND. Beginning in 2001, COGEND compared nicotine dependent smokers to smokers who never developed dependences symptoms.[5] Participants included EURs and AAs, who were aged 25 to 44 years old and recruited from St. Louis and Detroit. The FTND was administered to determine study eligibility as either nicotine dependent cases (current smokers who reported an FTND score of $\geq$4) or controls (smokers who reported >100 cigarettes during their lifetime but reported an FTND score $\leq$1). Participants were genotyped on either the Illumina Human1M-Duo array as part of the Study of Addiction: Genetics and Environment (SAGE)[6] or the Illumina HumanOmni2.5 array as part of the Gene Environment Association Studies Initiative (GENEVA).[7] The genotyping data are available via dbGaP accession numbers phs000092.v1.p1 and phs000404.v1.p1, respectively. We retained genotyped SNPs surpassing call rate >98% and HWE $P \geq 1\times10^{-4}$ thresholds in each subset, combined the subsets, removed duplicated participants and first-degree relatives, and retained only the SNPs genotyped at the intersection of the different arrays to circumvent potential bias.[8] After applying standard QC on the combined COGEND subsets, the final dataset included 1,935 EAs and 704 AAs with lifetime FTND scores and covariates (age, sex, and PC eigenvectors) for analysis.

Our analyses also included COGEND2 participants who were recruited more recently (2011–2014) following the COGEND study design. COGEND2 participants were genotyped on the Illumina Omni Express array, alongside the AAND participants, but analyzed separately. Following standard QC, there were 292 EURs and 313 AAs from COGEND2 for analysis with lifetime FTND scores and covariates (age, sex, and PC eigenvectors).

Center for Oral Health Research in Appalachia 1 (COHRA1). COHRA1 was primarily designed to conduct GWAS analyses of dental caries,[9] as one contributing site of a four-site study. COHRA1 was the only site that collected FTND data. COHRA1 recruited families beginning in 2003 from Appalachian regions: four rural counties (West Virginia and Pennsylvania) and an urban area. Eligible families included at least one adult and one biological child residing in the same household. We obtained COHRA1 genotyping data, as assayed on the Illumina Human610 array, via dbGaP accession number phs000095.v2.p1. We obtained FTND phenotype data from the original study investigators. When FTND data were available on parent(s) and children, we selected a single person from each relative pair/cluster based on the following criteria: (1) FTND data availability and (2) highest call rate if more than one relative had FTND data available. Following standard QC, we retained 243 EAs with current FTND scores and covariates (age, sex, and PC eigenvectors) for our final analysis dataset. Twelve of the participants were <18 years old.

Chronic Obstructive Pulmonary Disease Gene (COPDGene and COPDGene2). COPDGene is a longitudinal observational study of COPD with participants ascertained at multiple centers across the United States.[10] Participants, aged 45 to 80 years old, reported a current or former history of smoking and 10 or more cigarette pack-years. The Global Initiative for Chronic Obstructive Lung Disease (GOLD) criteria were used to stage disease severity among COPD cases based on their post-bronchodilator pulmonary function measures: GOLD = 1–2 for mild cases and 3–4 for moderate/severe cases. COPD controls had pulmonary function measures in the normal range for their sex, age and height. Acute and chronic respiratory disease, cancer and other conditions were used as exclusion criteria.

Our prior GWAS included 2,211 Non-Hispanic white (henceforth referred to as EUR) and 2,115 AA current smokers with current FTND data available from the baseline examination and with a determinant COPD case/control status.[1] For the present study, we added participants from the same examination with current FTND data available and an indeterminant COPD status (GOLD = -1). Together, we included 2,549 EUR and 2,534 AA current smokers from the baseline examination (denoted COPDGene1).

In a further expansion of COPDGene for the present study, we included participants who were not captured in COPDGene1 but had lifetime FTND data collected as part of the phase 2 follow-up examination (denoted COPDGene2; total N=2,630 EURs and 267 AAs). COPDGene2 comprised mostly former smokers but some current smokers, who had missing FTND at the baseline examination. Both COPDGene1 and COPDGene2 participants were genotyped on the Illumina HumanOmni1-Quad array and made available via dbGaP accession number phs000765.v1.p2. We conducted QC, imputation, and GWAS analysis for each ancestry in each of the two study phases, separately. Covariates for the GWAS analysis included age, sex, GOLD stage (-1 for indeterminant status, 1 or 2 for mild cases, and 3 or 4 for moderate/severe cases, with 0 for controls as the reference category), and PC eigenvectors.

deCODE**.** deCODE Genetics is a large population-based study from Iceland with data collection spanning 1996 to 2014. It was approved by the Data Protection Commission of Iceland and the National Bioethics Committee of Iceland. Participants were originally recruited to conduct genetic studies of smoking-related and a range of other phenotypes along with population controls. Personally identifiable information that was associated with phenotypic information and blood samples were encrypted by a third-party system.[11] Collection of smoking data has been described elsewhere.[12] Briefly, questionnaires were used to gather data on cigarettes per day (CPD) and the other FTND items.

Our prior ND GWAS meta-analyses included 9,090 smokers from deCODE.[1,2] The present analysis used an expanded sample size of 15,312 smokers using new smoking data collected in deCODE. Participants were genotyped on Illumina SNP arrays, and QC was applied as described before.[13] The ND definitions mimicked our prior analyses,[1,2] whereby mild dependence included smokers with lifetime FTND data (here, N=6333) as well as low-intensity smokers with CPD, but not the full-scale FTND, data available (N=8979). Smokers defined as moderately and severely dependent all had the full-scale FTND data available. See "FTND and categorical ND definitions for discovery GWAS" for further details. Association tests were carried out using a linear mixed model implemented in BOLT-LMM[14]. The FTND score was corrected for age and sex. LD score regression[15] was applied to account for inflation in test statistics due to cryptic relatedness and stratification. The $\chi 2$ statistics from the GWAS was regressed against LD score with a set of 1.1 M variants, and the intercept was used as the correction factor. The LD scores were downloaded from a LD score database (ftp://atguftp.mgh.harvard.edu/brendan/1k_eur_r2_hm3snps_se_weights.RDS; accessed 23 June 2015).

Environment and Genetics in Lung Cancer Etiology Study (EAGLE). EAGLE is a population-based study of newly diagnosed lung cancer cases and matched controls, who were aged 35 to 79 years old and recruited from the Italian region of Lombardy.[16,17] Genotyping was done on the Illumina HumanHap550v3 array, as part of GENEVA.[7] We obtained the genome-wide genotype, phenotype, and covariate data via dbGaP accession number phs000093.v2.p2 as well as the original study investigators. Lifetime FTND scores were collected among current and former smokers. Following our standard QC steps, the final analysis data set for EAGLE included 3,006 participants with complete data available on FTND scores and covariates (age, sex, and PC eigenvectors). As before,[1,2] lung cancer case/control status was not included as a covariate, because FTND scores were collected among current and former smokers based on lifetime and not current smoking habits.

Electronic Medical Records and Genomics (eMERGE) network**.** We obtained data from eMERGE participants in "A Genome-Wide Association Study on Cataract and HDL in the Personalized Medicine Research Project Cohort" via dbGaP accession number phs000170.v2.p1. These eMERGE participants were recontacted to collect data on a broad range of phenotypes and exposures to facilitate harmonization with other studies as part of the PhenX Rising project (https://www.genome.gov/27549243/phenx-rising/). FTND data were collected in this study based on current habits among current smokers and on period of maximum usage (i.e., lifetime) among former smokers. We combined the data from current and former smokers, given our prior findings that any measurement variance in the FTND has negligible effects on genetic association results, with very similar patterns observed between current and lifetime FTND.[18] The final analysis data set included 730 EURs. Covariates for GWAS analysis included age, sex, and PCs.

FINRISK**.** The population-based FINRISK study was initiated in 1972 with follow-up taking place every 5 years until 2012. Recruitment occurred in several geographic areas across Finland, making FINRISK a nationally representative study as previously described.[19] Genotyping was performed on the Illumina Human610-Quad or HumanCoreExome array, followed by QC and imputation with reference to the all-Finnish panel from the Sequencing Initiative Suomi project,[20] as described before.[21] The final analysis data set included 2,211 unrelated participants, including current and former smokers, with complete data on lifetime FTND scores and covariates (age, sex, and PC eigenvectors).

Finnish Twin Cohort (FTC). As before,[1] FTC participants originated from these sub-cohorts: the Nicotine Addiction Genetics study of adult twins, born 1938–1957 and concordant for being ever smokers, and their relatives (mainly siblings); and population-based longitudinal studies of five consecutive Finnish twin birth cohorts from 1983–1987 (FinnTwin12) and 1975–1979 (FinnTwin16).[22,23] The FTC sample size has increased from our prior GWAS analyses[1] due to new genotyping data. Genotyping was done using Illumina's Human610-Quad, Human670-QuadCustom, or HumanCoreExome array. QC was performed in two batches—(1) Human610-Quad and Human670-QuadCustom together and (2) HumanCoreExome—with variants removed for low call rate (<97.5% in batch 1 or <95% in batch 2), MAF<1%, or HWE $P<1\times10^{-6}$ and participants removed for low call rate (<98% for batch 1 or <95% for batch 2), excessive heterozygosity, discordant sex, or ancestry outlier. Imputation was conducted separately by genotyping array with Minimac3 v2.0.1 using the Michigan Imputation Server.[24] Imputed variants were merged across batches to construct the final analysis dataset of 2,507 participants with complete data on lifetime FTND scores and covariates (age, sex, birth cohort, and PC eigenvectors). Their kinship matrix was taken into account as a random effect in a linear mixed model. Imputation quality scores were re-calculated across the merged batches using the impute-info plugin for BCFtools.

Molecular Genetics of Schizophrenia—Genetic Association Information Network (GAIN) and nonGAIN studies. The overarching Molecular Genetics of Schizophrenia study was designed as a United States-based case-control study of schizophrenia/schizoaffective disorder. Cases were diagnosed with schizophrenia or schizoaffective disorder according to

DSM-IV criteria, whereas controls were assessed and determined to have no history of these illnesses. One subset of the study participants were genotyped as part of GAIN[25] with data obtained via dbGaP accession number phs000021.v3.p2, whereas the other subset was genotyped separately and denoted nonGAIN with data obtained via dbGaP accession number phs000167.v1.p1. Both subsets were genotyped using the Affymetrix 6.0 array. For the prior[1] and present ND GWAS analyses, we applied our standard QC steps and used only the schizophrenia controls. The final analysis datasets included 774 EAs from GAIN, 477 AAs from GAIN, and 471 EAs from nonGAIN with complete data available on lifetime FTND scores and covariates (age, sex, and PC eigenvectors for each dataset analyzed separately).

**German Nicotine Cohort study (NCS) (German study).** The German study is a population-based case-control study specifically conducted to assess the genetics of ND[26]. Data collection occurred from 2007-2009 at 7 recruitment centers across Germany (Departments of Psychiatry at the Universities of Aachen, Berlin, Bonn, Düsseldorf, Erlangen, Mainz, Mannheim). Probands were randomly selected from the local population via residents' registers at each site, and subjects were required to meet the following inclusion criteria: age 18-65 years; current smoker or occasional smoker (>=7 cigarettes per week or 1 cigarette per day) or never smoker (<=20 cigarettes over lifetime); grandparents born in Germany or adjacent country; native-level German language proficiency; letter invitation via official local residents' register. Furthermore, the following exclusion criteria were applied: former smoker; alcohol or substance abuse within previous six months (DSM-IV); a history of alcohol or substance dependence (DSM-IV); DSM-IV axis-1 psychiatric diagnosis within previous six months; non-German origin; not native-level proficient in German language; pregnant; any medical condition that may interfere with the study; CNS-relevant medication within previous 6 months; CNS-relevant (neurological) illnesses (lifetime). Out of 55,000 subjects contacted, 2,396 were enrolled in the study.

DNA extracted from whole-blood samples acquired from study subjects were genotyped using the Illumina InfiniumOmniExpressExome-8v1-3_A array. Genotype QC steps included missing rate (missing rate>=0.05 and MAF>=0.05 or missing rate>=0.03 and MAF<0.05) and HWE $P<5.38\times10^{-8}$. Subject QC steps included missing rate >=5%, excess heterozygosity (plink --het, F more than 2*sigma deviations from the mean), high degree of relatedness (plink –genome full, pi_hat>=0.26), and PCA-based ancestral outlier removal (1000 Genomes Phase 3 reference). Following QC, imputation was performed using IMPUTE2 with the 1000 Genomes Phase 3 reference panel. The final analysis dataset with complete phenotype and genotype information included 991 current smokers of EUR ancestry.

Jackson Heart Study (JHS) / Atherosclerosis Risk in Communities (ARIC)**.** The longitudinal JHS was designed to evaluate cardiovascular disease risk among AAs from the general population in Jackson, Mississippi and its surrounding area.[27] JHS was an extension of the ARIC study of EURs and AAs from 4 communities across the United States (Jackson, Mississippi; Forsyth County, North Carolina; suburbs of Minneapolis, Minnesota; and Washington County, Maryland).[28] JHS recruited AAs were aged 35 to 84 years old, alongside their relatives who were aged 21 to 34 years old. Across JHS and ARIC, smokers were defined based on reports of having smoked 400 or more cigarettes in their lifetime. In parallel with the approach taken in deCODE, we included smokers with lifetime FTND data (N=682, all from JHS) and augmented the sample size by including 461 low-intensity AA smokers from ARIC with only CPD data available in the mild ND category (see "FTND and categorical ND definitions for discovery GWAS" for additional details).

Genotyping for both JHS and ARIC were performed on the Affymetrix 6.0 array, and we obtained these data via dbGaP (accession numbers phs000286.v3.p1 for JHS and phs000090.v1.p1 for ARIC). After applying our standard QC, there were 1,143 AA smokers with FTND scores or CPD reported and covariate (age, sex, and PC eigenvectors) data available: N=628 from JHS and 515 from ARIC. We confirmed that no participants were duplicated across the JHS and ARIC subsets in our final analysis data set, with identity-by-state estimates <0.9 for all pairwise comparisons.

Minnesota Center for Twin and Family Research (MCTFR)**.** MCTFR is composed of two longitudinal studies, the Minnesota Twin Family Study and the Sibling Interaction and Behavior Study. The Minnesota Twin Family Study recruited three studies of twin pairs and their parents and the Sibling Interaction and Behavior Study recruited adoptive and biological siblings and their parents. Families were initially recruited as a community study to study a broad range of psychological domains. Altogether, we included data for 1,073 current and former smokers, with lifetime FTND data available, who were genotyped on the Illumina 660W-Quad. Their genotyping protocols and QC were described previously.[29,30] Sex and age were included as covariates and to account for family relatedness, we used a kinship matrix and included PC eigenvectors as covariates. Additional covariates were included based on sample ascertainment and structure; we used four dummy coded variables to account for each of the three Minnesota Twin Family Study intake studies and the Sibling Interaction and Behavior Study, and a variable indicating if an individual was a parent.

Netherlands Twin Registry (NTR)**.** The NTR began in 1987 as a longitudinal study of twins and other multiple birth siblings. The NTR is comprised of two collections: (1) adult twins and their family members, and (2) younger twins recruited at birth or in early life, their parents, and their siblings.[31] Genome-wide genotyping was performed on a subset of NTR participants using various Affymetrix and Illumina arrays,[32,33] followed by QC as described elsewhere.[33] Genotyped SNPs passing QC were merged across different arrays and used for imputation. Imputed SNPs were filtered out for the following reasons: MAF<0.5%, HWE $P<1\times10^{-5}$, estimated $r^2<0.3$, Mendelian error rate<2%, or absolute reference frequency allele difference>0.15 between NTR and 1000G. With an increased sample size from before due to a continued increase in the number of NTR participants who were genotyped, the present analysis included 4,489 NTR participants who had lifetime FTND[34] and covariate (age, sex, dummy variables to correct for genotyping array, and PC eigenvectors) data available.

GWAS of Alcohol Use and Alcohol Use Disorder in Australian Twin-Families (OZ-ALC) Study. Data for the present study were obtained from dbGaP study "International Consortium on the Genetics of Heroin Dependence" (accession number phs000277.v1.p1), for which OZ-ALC participants served as a source of DSM-IV-assessed non-opioid dependent controls. No other components of the heroin dependence study had FTND data. The OZ-ALC study data were derived from telephone diagnostic interviews of Australian twins from the general population and their spouses. Alcohol dependent cases from OZ-ALC were minimized for inclusion in the heroin dependence study made available in dbGaP. We began with the 1,172 participants, who were all of Australian European ancestry, genotyped on the CIDR370v1 or CIDR370v3 array, and had lifetime FTND data available.[35] Genotype imputation was based on the overlap of the two arrays. The final analysis dataset included 1,138 unrelated participants. Our statistical analyses included adjustment for age, sex, and PCs.

Study of Addiction: Genetics and Environment (SAGE)**.** SAGE was assembled from three case-control studies collected in the United States for addictive disorders: COGEND, the

Collaborative Study on the Genetics of Alcoholism (COGA),[36] and the Family Study of Cocaine Dependence (FSCD).[37] Genotyping was conducted using the Illumina Human1M-Duo array, from which we obtained via dbGaP accession number phs000092.v1.p1. COGEND participants were removed to avoid participant overlap; all other participants from COGA and FSCD (henceforth referred to as SAGE*) were analyzed together as previously done.[1,2,6] Following our standard QC, there remained 832 EAs and 633 AAs with lifetime FTND scores and covariates (age, sex, DSM-IV-defined cocaine dependence, DSM-IV-defined alcohol dependence, and PC eigenvectors) for GWAS analysis.

Spit for Science**.** Spit for Science, is an ongoing longitudinal study of Virginia Commonwealth University students. Briefly, incoming students age 18 or older were eligible to complete phenotypic assessments, which covered a wide range of topics but focused on alcohol use.[38] Study data were collected and managed using REDCap electronic data capture tools[39] hosted at Virginia Commonwealth University. Follow-up assessments were completed in subsequent spring semesters. Individuals who did not participate in the first wave of data collection (including those who turned 18 after the end of the first wave of data collection) had the opportunity to join the study the following spring; those who participated during their first year were eligible to complete follow-up assessments each spring. Participants who completed the phenotypic assessments were eligible to provide a DNA sample. There was a total of 7,603 participants across three studies, which matriculated in Fall 2011 (N=2,714), 2012 (N=2,486), and 2013 (N=2,403). Of these, 98% provided a DNA sample. The current analyses are based on FTND data captured after the Spring 2014 survey, with data available for up to 4 waves per participant. Lifetime FTND data were collected among current and former smokers, using the FTND with the heaviest smoking reported when data were available from more than one wave.

Genotyping was performed on the Affymetrix BioBank array, and QC steps were applied as detailed elsewhere.[40] For this study, we used only genotyped EURs, which was the largest ancestry group and had sufficient representation in each of the three ND categories (mild/moderate/severe). Following QC, there were 1,717 individuals with FTND scores and covariate data (age, sex, and PCs) available.

University of Wisconsin-Transdisciplinary Tobacco Use Research Center (UW-TTURC). UW-TTURC represents a collection of smokers recruited from Madison and Milwaukee, Wisconsin, beginning in 2001, for smoking cessation treatment clinical trials.[9] Participants were deemed eligible, based on having smoked at least 10 CPD and reported being motived to quit smoking. Genotyping was performed using the Illumina HumanOmni2.5 array. We obtained their genotypes, FTND scores, and covariate data via dbGaP accession number phs000404.v1.p1. After applying standard QC, there remained 1,534 EAs and 247 AAs with current FTND scores and covariate data (age, sex, and PC eigenvectors) for analysis.

Yale-Penn. The Yale-Penn study was designed to conduct genetic studies for addiction using mostly unrelated individuals but also small nuclear families, all of whom were recruited from the eastern United States.[41-43] ND was not considered in the inclusion or exclusion criteria, but lifetime FTND data were collected among smokers.[44] Genotyping was conducted on the Illumina HumanOmni1-Quad array. QC mimicked prior analysis,[1] except that ancestry assignments were refined using K-means clustering to assign individuals based on the nearest centroid across the first 10 PC eigenvectors with reference to 1000G EUR or AFR population. There were 1,579 EAs and 2,637 AAs in the final analyses, which included adjustment for age, sex, and PC eigenvectors.

### FTND and categorical ND definitions for discovery GWAS

Our discovery GWAS meta-analyses included studies with ND defined by the full 6-item FTND. The FTND[3] queries the following six items:

(1) **How soon after you wake up do/did you smoke your first cigarette?** Categorical responses: within 5 minutes, 6–30 minutes, 31–60 minutes, or after 60 minutes.

(2) **Do/Did you find it difficult to refrain from smoking in places where it is forbidden, e.g., in church, at the library, in a cinema, etc.?** Binary response: yes or no.

(3) **Which cigarette would you hate most to give up?** Binary response: the first one in the morning or all others.

(4) **How many cigarettes per day do/did you smoke?** Categorical response: 10 or less, 11–20, 21–30, or 31 or more.

(5) **Do/did you smoke more frequently during the first hours after waking than during the rest of the day?** Binary response: yes or no.

(6) **Do/did you smoke if you are so ill that you are in bed most of the day?** Binary response: yes or no.

Additional details on the protocol and scoring algorithm are provided in the PhenX Toolkit[45], a catalog of commonly ascertained phenotype and exposure measures: https://www.phenxtoolkit.org/protocols/view/31001. Briefly, FTND scores range from 0 (no dependence) to 10 (highest dependence level). FTND can be administered on current or former smokers based on the time period when they reported smoking the most (i.e., lifetime FTND) or among current smokers based around the time of interview (i.e., current FTND). We used lifetime FTND collected among current and former smokers in AAND, ADAA, EAGLE, COGEND, COGEND2, COPDGene2, deCODE, eMERGE, FINRISK, FTC, GAIN, JHS/ARIC,

MCTFR, nonGAIN, NTR, OZ-ALC, SAGE*, Spit for Science, and Yale-Penn. We used current FTND that was available in COHRA1, COPDGene, eMERGE, German, and UW-TTURC.

We used the FTND to derive a categorical variable for ND: scores of 0–3 for mild, 4–6 for moderate, and 7–10 for severe. We relied solely on the FTND to define ND, except in two of the 23 studies (deCODE and JHS/ARIC) where we included smokers with FTND data available as well as low-intensity smokers who had only CPD data that we used as a proxy measure to define mild dependence (CPD ≤10) as done before.[1,2] Our prior assessment showed high concordance of CPD ≤10 and FTND scores 0–3 (86.4%),[1] suggesting that CPD can be used to define mild dependence with little phenotype misclassification. However, any phenotype misclassification would be expected to conservatively bias results, leading to reduced statistical power, attenuated effect size estimates, and thus underestimate SNP associations with ND.[46,47] Moderate and severe dependence was defined solely by FTND scores across all studies, as our prior assessment showed lower concordance between FTND and CPD for defining these categories.[1] In follow-up analyses of lead SNPs from novel loci, we tested associations of each specific FTND item, using linear regression models for items with categorical responses (items #1 and 4) and logistic regression models for items with binary responses (items #2, 3, 5, and 6), followed by meta-analysis of results across studies. Due to varying genotype and phenotype data availability for the novel loci and specific FTND items, some studies could not utilize the full sample set for specific FTND item testing. These studies include AAND, COGEND, COGEND2, deCODE, Dental Caries, GAIN, German, JHS/ARIC, and nonGAIN.[48,49]

### Heaviness of smoking index (HSI) in the UK Biobank for independent testing

Since there are no other datasets with comparably large sample sizes, we relied on the HSI that is available in the UK Biobank for independent testing of our genome-wide significant

FTND-based GWAS meta-analysis findings. The UK Biobank collected data on two of the 6 FTND items (CPD and time to first cigarette in the morning [TTFC]) among current smokers, who reported smoking on most or all days. These two items comprise the HSI, which has historically been considered a suitable proxy for the full-scale FTND.[3] To evaluate the agreement in our FTND categories (score range = 0–10; mild [scores 0–3], moderate [scores 4–6], and severe [scores 7–10] as we have routinely used before[1,2]) with HSI categories (score range = 0–6; mild [scores 0–2], moderate [scores 3–4], and severe [scores 5–6], in accordance with the scoring algorithm by the American Society of Clinical Oncology[50]), we used EURs in COGEND, which was ascertained specifically for ND. We compared FTND and HSI categories based on lifetime FTND (i.e., FTND based on time smoked most). Results are presented in **Supplementary Table 15**. Concordance was highest for mild dependence at 94.9%; i.e., among those defined as having mild dependence by the HSI, 94.87% also had mild dependence as defined by the full-scale FTND. We observed concordance of 81.2% for moderate dependence and 84.9% for severe dependence. Overall concordance was high (89.3%), corroborating the utility of the HSI categories as a proxy for defining ND in the UK Biobank.

Genome-wide significant SNP associations from our GWAS meta-analysis were tested for association using 33,791 current smokers with HSI data available in UK Biobank: 18,063 mild (HSI scores 0–2), 13,395 moderate (HSI scores 3–4), and 2,333 severe (scores 5–6) dependence. This final analysis data set included only unrelated individuals, as we removed 844 third-degree or closer relatives prior to analysis; for each related pair/cluster, individuals who had more relatives and who were light smokers were prioritized for removal. For our SNP-HSI association testing, we followed the model employed by systemic GWAS analyses for a multitude of phenotypes (see http://www.nealelab.is/uk-biobank/), adjusting for the following covariates: sex, age, age$^2$, age $\times$ sex, age$^2$ $\times$ sex, and PC eigenvectors, with the age$^2$ and interaction terms among the age and sex variables intended to account for non-linear associations.

### Acknowledgements

AAND was funded by the National Institute on Drug Abuse (NIDA) grant R01 DA025888 (Multiple Principal Investigators [MPIs] Eric O. Johnson and Laura J. Bierut).

ADAA data were funded by the National Institute on Alcohol Abuse and Alcoholism (NIAAA) grant R01 AA017444 (PI: Richard Grucza).

COGEND and COGEND2 were supported by grants from the National Cancer Institute (NCI; grant number P01 CA089392, PI: Laura Bierut) and NIDA (R01 DA036583 and R01 DA025888, PI: Laura Bierut), both of the National Institutes of Health (NIH). Genotype data are available via dbGaP as part of the "Genetic Architecture of Smoking and Smoking Cessation" (accession number phs000404.v1.p1) and "Study of Addiction: Genetics and Environment (SAGE)" (accession number phs000092.v1.p1). Funding support for genotyping, which was performed at CIDR, was provided by 1 X01 HG005274-01 and by the NIH Genes, Environment and Health Initiative [GEI] (U01 HG004422). CIDR is fully funded through a federal contract from the NIH to The Johns Hopkins University, contract number HHSN268200782096C. Assistance with genotype cleaning, as well as with general study coordination, was provided by the GENEVA Coordinating Center (U01 HG004446).

Funding support for the Center for Oral Health Research in Appalachia (COHRA) Dental Caries GWAS study (study designation COHRA1) was provided by the NIH, National Institute of Dental and Craniofacial Research (NIDCR) grant number U01 DE018903 (PI: MLM). Data and samples were provided by COHRA, a collaboration of the University of Pittsburgh and West

Virginia University funded by NIDCR R01 DE014899 (PIs: MLM and DWM). Genotype and phenotype data are available through dbGaP (accession number phs000095.v2.p1).

The COPDGene® project was supported by award numbers R01 HL089897 and R01 HL089856 from the National Heart, Lung, and Blood Institute (NHLBI). Research reported in this publication was also supported by the NHLBI award number K01 HL125858. The content is solely the responsibility of the authors and does not necessarily represent the official views of NHLBI or NIH. The COPDGene® project is also supported by the COPD Foundation through contributions made to an Industry Advisory Board comprised of AstraZeneca, Boehringer Ingelheim, Novartis, Pfizer, Siemens, Sunovion, and GlaxoSmithKline. The authors acknowledge investigators of the COPDGene® project core units: Administrative (MPIs James Crapo and Edwin Silverman), Barry Make, and Elizabeth Regan); Genetic Analysis (Terri Beaty, Nan Laird, Christoph Lange, Michael Cho, Stephanie Santorico, Dawn DeMeo, Nadia Hansel, Craig Hersh, Peter Castaldi, Merry-Lynn McDonald, Emily Wan, Megan Hardin, Jacqueline Hetmanski, Margaret Parker, Marilyn Foreman, Brian Hobbs, Robert Busch, Adel El-Bouiez, Megan Hardin, Dandi Qiao, Elizabeth Regan, Eitan Halper-Stromberg, Ferdouse Begum, Sungho Won); Imaging (David Lynch, Harvey Coxson, MeiLan Han, Eric Hoffman, Stephen Humphries Francine Jacobson, Philip Judy, Ella Kazerooni, John Newell, Jr., Elizabeth Regan, James Ross, Raul San Jose Estepar, Berend Stoel, Juerg Tschirren, Eva van Rikxoort, Bram van Ginneken, George Washko, Carla Wilson, Mustafa Al Qaisi, Teresa Gray, Alex Kluiber, Tanya Mann, Jered Sieren, Douglas Stinson, Joyce Schroeder, Edwin Van Beek); Pulmonary Function Testing Quality Assurance (Robert Jensen); Data Coordinating Center and Biostatistics (Douglas Everett, Anna Faino, Matt Strand, Carla Wilson); and Epidemiology (Jennifer Black-Shinn, Gregory Kinney, Katherine Pratte). The authors also acknowledge the clinical center investigators: Jeffrey Curtis, Carlos Martinez, Perry G. Pernicano, Nicola Hanania, Philip Alapat, Venkata Bandi, Mustafa Atik, Aladin Boriek, Kalpatha Guntupalli, Elizabeth Guy, Amit Parulekar, Arun Nachiappan, Dawn DeMeo, Craig Hersh, George Washko, Francine Jacobson, R. Graham Barr, Byron Thomashow, John Austin, Belinda D'Souza, Gregory D.N. Pearson, Anna Rozenshtein, Neil MacIntyre, Jr., Lacey Washington, H. Page McAdams, Charlene McEvoy, Joseph Tashjian, Robert Wise, Nadia Hansel, Robert Brown, Karen Horton, Nirupama Putcha, Richard Casaburi, Alessandra Adami, Janos Porszasz, Hans Fischer, Matthew Budoff, Dan Cannon, Harry Rossiter, Amir Sharafkhaneh, Charlie Lan, Christine Wendt, Brian Bell, Marilyn Foreman, Gloria Westney, Eugene Berkowitz, Russell Bowler, David Lynch, Richard Rosiello, David Pace, Gerard Criner, David Ciccolella, Francis Cordova, Chandra Dass, Robert D'Alonzo, Parag Desai, Michael Jacobs, Steven Kelsen, Victor Kim, A. James Mamary, Nathaniel Marchetti, Aditti Satti, Kartik Shenoy, Robert M. Steiner, Alex Swift, Irene Swift, Gloria Vega-Sanchez, Mark Dransfield, William Bailey, J. Michael Wells, Surya Bhatt, Hrudaya Nath, Joe Ramsdell, Paul Friedman, Xavier Soler, Andrew Yen, Alejandro Cornellas, John Newell, Jr., Brad Thompson, MeiLan Han, Ella Kazerooni, Fernando Martinez, Joanne Billings, Tadashi Allen, Frank Sciurba, Divay Chandra, Joel Weissfeld, Carl Fuhrman, Jessica Bon, Antonio Anzueto, Sandra Adams, Diego Maselli-Caceres, and Mario Ruiz.

The deCODE work was supported in part by NIDA R01 DA017932 (PI: Kari Stefansson) and R01s DA035825 and DA042090 (PI: Dana Hancock). deCODE thanks the participants in the genetic studies whose contributions made this work possible.

Funding support for the EAGLE study was provided through the NIH GEI (Z01 CP 010200). The human participants participating in this study and its companion "Prostate, Lung Colon and Ovary Screening Trial" are supported by intramural resources of the NCI. Assistance with phenotype harmonization and genotype cleaning, as well as with general study coordination, was provided by the GENEVA Coordinating Center (U01 HG004446). Assistance with data cleaning was provided by the National Center for Biotechnology Information (NCBI). Funding support for genotyping, which was performed at The Johns Hopkins University's Center for Inherited Disease Research (CIDR), was provided by the NIH GEI (U01 HG004438). The datasets used for the analyses described in this manuscript were obtained from the database of Genotypes and Phenotypes (dbGaP, http://www.ncbi.nlm.nih.gov/gap) through accession number phs000093.v2.p2.

For eMERGE, funding support was derived from the Personalized Medicine Research Project (PMRP) through a cooperative agreement (U01 HG004608) with the National Human Genome Research Institute (NHGRI), with additional funding from the National Institute for General Medical Sciences. The samples used for PMRP analyses were obtained with funding from Marshfield Clinic, Health Resources Service Administration Office of Rural Health Policy grant number D1A RH00025, and Wisconsin Department of Commerce Technology Development Fund contract number TDF FYO10718. Funding support for genotyping, which was performed at Johns Hopkins University, was provided by the NIH (U01HG004438). Assistance with phenotype harmonization and genotype data cleaning was provided by the eMERGE Administrative Coordinating Center (U01 HG004603) and NCBI. The datasets used for the analyses described in this manuscript were obtained from dbGaP via dbGaP accession number phs000170.v1.p1.

The FINRISK authors would like to acknowledge the National Institute for Health and Welfare (Helsinki, Finland) for data collection. The surveys and clinical examination were approved by local ethical committees, and participants provided written informed consent. The authors also thank the Biobank of the National Institute for Health and Welfare for making data available to external researchers at https://thl.fi/en/web/thl-biobank.

The FTC authors warmly thank the participating twin pairs and their family members of the Finnish Twin Cohort Study for their contribution. We would like to express our appreciation to the skilled study interviewers A-M Iivonen, K Karhu, H-M Kuha, U Kulmala-Gråhn, M Mantere, K Saanakorpi, M Saarinen, R Sipilä, L Viljanen, and E Voipio. Anja Häppölä and Kauko Heikkilä are acknowledged for their valuable contribution in recruitment, data collection, and data management. Phenotyping and genotyping of the Finnish twin cohorts was supported by the Academy of Finland Center of Excellence in Complex Disease Genetics (grants 213506, 129680), the Academy of Finland (grants 100499, 205585, 118555, 141054, 265240, 263278 and 264146 to Jaakko Kaprio), NIAAA (grant numbers R37 AA012502, K05 AA000145, and R01 AA009203 to R. J. Rose and R01 AA015416 and K02 AA018755 to D. M. Dick), NIDA (grant number R01 DA012854, PI: Pamela A. F. Madden), Sigrid Juselius Foundation (to Jaakko Kaprio), Global Research Award for Nicotine Dependence, Pfizer Inc. (to Jaakko Kaprio), and the Welcome Trust Sanger Institute, UK. Antti-Pekka Sarin and Samuli Ripatti are acknowledged for genotype data quality controls and imputation. Association analyses were run at the ELIXIR Finland node hosted at CSC – IT Center for Science for ICT resources.

The GAIN and nonGAIN studies originated from the same overarching Genome-Wide Association of Schizophrenia Study as provided by the Molecular Genetics of Schizophrenia Collaboration (PI: Pablo V. Gejman, Evanston Northwestern Healthcare [ENH] and Northwestern University, Evanston, IL, USA). For the GAIN subset, funding support was provided by the National Institute of Mental Health (NIMH) (R01 MH67257, R01 MH59588, R01 MH59571, R01 MH59565, R01 MH59587, R01 MH60870, R01 MH59566, R01 MH59586,

R01 MH61675, R01 MH60879, R01 MH81800, U01 MH46276, U01 MH46289 U01 MH46318, U01 MH79469, and U01 MH79470) and the genotyping of samples was provided through GAIN. The datasets used for the analyses described in this manuscript were obtained from dbGaP through accession number phs000021.v3.p2. Funding support for the nonGAIN subset was provided by Genomics Research Branch at NIMH, and the genotyping and analysis of samples was also provided through GAIN and under the MGS U01s: MH79469 and MH79470. Assistance with data cleaning was provided by NCBI. The dataset used for the analyses described in this manuscript was obtained via dbGaP accession number phs000167.v1.p1. Samples and associated phenotype data for the nonGAIN study were collected under the following grants: NIMH Schizophrenia Genetics Initiative U01s: MH46276 (CR Cloninger), MH46289 (C Kaufmann), and MH46318 (MT Tsuang); and MGS Part 1 (MGS1) and Part 2 (MGS2) R01s: MH67257 (NG Buccola), MH59588 (BJ Mowry), MH59571 (PV Gejman), MH59565 (Robert Freedman), MH59587 (F Amin), MH60870 (WF Byerley), MH59566 (DW Black), MH59586 (JM Silverman), MH61675 (DF Levinson), and MH60879 (CR Cloninger). Further details of collection sites, individuals, and institutions may be found in data supplement Table 1 of Sanders et al. (2008; PMID: 18198266) and at the study dbGaP pages.

The study 'Genetics of Nicotine Dependence and Neurobiological Phenotypes' (Wi1316/9-1—Principle Investigator: Georg Winterer) was funded by the German Research Foundation (Deutsche Forschungsgemeinschaft, DFG) as part of the Priority Program (Schwerpunktprogramm) SPP1226: Nicotine—Molecular and Physiological Effects in CNS (Coordinator: Georg Winterer; http://www.nicotine-research.com). Additional funding was obtained by Georg Winterer for follow-up investigations from the DFG as part of the Collaborative Research Centre Transregio 265 (Losing and Regaining control over drug in take).

Funding was also provided to Marcella Rietschel for conducting genome-wide analyses by the European Community's Seventh Framework Programme (CRESTAR: Pharmacogenomic biomarkers as clinical decision making tools for clozapine treatment of schizophrenia, EU-FP7: 279227), from the German Federal Ministry of Education and Research (BMBF) through ERA-NET NEURON, 01EW1810, "SynSchiz - Linking synaptic dysfunction to disease mechanisms in schizophrenia - a multilevel investigation", from the German Federal Ministry of Education and Research (BMBF) through e:Med SysmedSUD grant 01ZX01909A "A systems-medicine approach towards distinct and shared resilience and pathological mechanisms of substance use disorders", and from the German Research Foundation (DFG) through SFB/Transregio 265 "Losing and Regaining Control over Drug Intake - S01: Central recruitment, imaging and biobanking". The German Nicotine Cohort Study acknowledges the following major contributors: Arian Mobascher (Department of Psychiatry, Mainz University Hospital, Mainz, Germany), Oliver Tüscher (Department of Psychiatry, Mainz University Hospital, Mainz, Germany), Amalia Diaz-Lacava (Cologne Center for Genomics, University of Cologne, Cologne, Germany), Peter Nürnberg (Cologne Center for Genomics, University of Cologne, Cologne, Germany), Thomas F. Wienker (Max-Planck-Institute for Molecular Genetics, Berlin, Germany), Michael Wagner (Department of Psychiatry, Bonn University Hospital, Bonn, Germany), Jürgen Gallinat (Department of Psychiatry and Psychotherapy, University Medical Center Hamburg Eppendorf (UKE), Hamburg, Germany), Gerhard Gründer (Department of Molecular Neuroimaging, Central Institute of Mental Health, Medical Faculty Mannheim, Heidelberg University Mannheim, Germany), Falk Kiefer (Department of Addictive Behaviour and Addiction Medicine, Central Institute of Mental Health, Medical Faculty Mannheim, Heidelberg University, Mannheim, Germany), Josef Frank (Department of Genetic

Epidemiology in Psychiatry, Central Institute of Mental Health, Medical Faculty Mannheim, Heidelberg University, Mannheim, Germany), Johannes Kornhuber (Department of Psychiatry, Friedrich-Alexander University, University Hospital, Erlangen- Nürnberg, Erlangen, Germany), and Markus Nöthen (Institute of Human Genetics, University Hospital Bonn, Bonn, Germany, Department of Genomics, Life & Brain Center, University of Bonn, Bonn).

JHS (dbGaP accession no. phs000286.v3.p1) is supported and conducted in collaboration with Jackson State University (HHSN268201300049C and HHSN268201300050C), Tougaloo College (HHSN268201300048C), and the University of Mississippi Medical Center (HHSN268201300046C and HHSN268201300047C) contracts from the NHLBI and the National Institute for Minority Health and Health Disparities. Funding for genotyping was provided by NHLBI contract N01-HC-65226. ARIC (dbGaP accession number phs000090.v1.p1) is carried out as a collaborative study supported by NHLBI contracts N01-HC-55015, N01-HC-55016, N01-HC-55018, N01-HC-55019, N01-HC-55020, N01-HC-55021, N01-HC-55022, R01 HL087641, R01 HL59367 and R01 HL086694; NHGRI contract U01 HG004402; and NIH contract HHSN268200625226C. The authors thank the staff and participants of the ARIC study for their important contributions. Infrastructure was partly supported by grant UL1RR025005, a component of the NIH and NIH Roadmap for Medical Research. Genotyping was conducted by GENEVA. Funding for GENEVA was provided by NHGRI grant U01 HG004402 (E. Boerwinkle).

Research reported in this publication for the MCTFR was supported by NIH awards R01 DA037904, R01 DA044283, R01 HG008983, R21 DA046188, R21 DA040177, R01 DA042755, R37 DA005147, R37 AA009367, R01 DA036216.

We thank all NTR participants whose data we analyzed in this study. Camelia C. Minica and Michael M. Neale are supported by the NIDA grant R01 DA018673. We acknowledge the Netherlands Organisation for Scientific Research (NWO) and the Netherlands Organisation for Health Research and Development (ZonMW Addiction 31160008; ZonMW 940-37-024; NWO/SPI 56-464-14192; NWO-400-5-717; NWO-MW 904-61-19; NWO-MagW 480-4-004; NWO-Veni 016-115-035; NWO-Groot 480-15-001/674), Amsterdam Public Health (APH), BBMRI–NL (BBMRI –NL, 184.021.007 and 184.033.111: Biobanking and Biomolecular Resources Research Infrastructure), the Avera Institute, Sioux Falls, South Dakota (USA) and the European Research Council (230374, 284167) for support. Genotyping was funded in part by grants from NIH (4R37DA018673-6, RC2 MH089951). The statistical analyses were carried out on the Genetic Cluster Computer (http://www.geneticcluster.org) which is supported by the Netherlands Scientific Organization (NWO 480-5-003), the Dutch Brain Foundation and the Department of Psychology and Education of the VU University Amsterdam.

We acknowledge the contribution of data from the OZ-ALC subset of the GWAS of Heroin Dependence study that was supported by R01 DA17305 (PI: Elliot Nelson) and accessed through dbGaP accession number phs000277.v1.p1. We also thank Dr. Nelson for feedback on the overarching study design, as related to the current study.

Funding support for SAGE was further provided through the NIH GEI (U01 HG004422). Assistance with phenotype harmonization and genotype cleaning, as well as with general study coordination, was provided by the GENEVA Coordinating Center (U01 HG004446). Assistance with data cleaning was provided by NCBI. Support for collection of datasets and samples was provided by the Collaborative Study on the Genetics of Alcoholism (COGA; U10 AA008401) and the Family Study of Cocaine Dependence (FSCD; R01 DA013423, PI: Laura Bierut).

Funding support for genotyping, which was performed at CIDR, was provided by the NIH GEI (U01HG004438), NIAAA, NIDA, and the NIH contract "High throughput genotyping for studying the genetic contributions to human disease" (HHSN268200782096C). The datasets used for the analyses described in this manuscript were obtained via dbGaP accession number phs000092.v1.p1.

Spit for Science has been supported by Virginia Commonwealth University, P20 AA017828, R37AA011408, K02AA018755, and P50 AA022537 from NIAAA, and UL1RR031990 from the National Center for Research Resources and NIH Roadmap for Medical Research.

The UW-TTURC study was accessed via dbGaP as part of “Genetic Architecture of Smoking and Smoking Cession” (accession number phs000404.v1.p1). Funding support for collection of the UW-TTURC dataset and samples was provided by P50 DA019706 and P50 CA084724. Funding support for genotyping, which was performed at CIDR, was provided by 1 X01 HG005274-01. CIDR is fully funded through a federal contract from NIH to The Johns Hopkins University, contract number HHSN268200782096C. Assistance with genotype cleaning, as well as with general study coordination, was provided by the GENEVA Coordinating Center (U01 HG004446).

The Yale-Penn study was supported by NIH grants RC2 DA028909 (PI: Joel Gelernter), R01 DA012690 (PI: Joel Gelernter), R01 DA012849 (PI: Joel Gelernter), R01 DA018432 (PI: Henry Kranzler), R01 AA011330 (PI: Joel Gelernter), R01 AA017535 (PI: Joel Gelernter), and the VA Connecticut and Philadelphia VA MIRECCs. Genotyping services for a part of the Yale-Penn GWAS were provided by CIDR and Yale University (Center for Genome Analysis). CIDR is fully funded through a federal contract from NIH to The Johns Hopkins University (contract number N01-HG-65403). We are grateful to Ann Marie Lacobelle, Catherine Aldi, and Christa Robinson for their excellent technical assistance, to the SSADDA interviewers, led by Yari Nuñez and Michelle Slivinsky, who devoted substantial time and effort to phenotype the study, and to John Farrell and Alexan Mardigan for database management assistance.

**Supplementary Table 1.** Genome-wide significant single nucleotide polymorphism (SNP) and insertion/deletion (indel) associations from the cross-ancestry genome-wide association study (GWAS) meta-analysis (total N=58,000). The SNPs and indels span five loci: chromosomes 5q34, 7q21, 9q34, 15q25, and 20q13. "Direction" indicates the association of the "Allele1" allele, corresponding to the "Effect" (β coefficient, with + corresponding to increased risk and – corresponding to decreased risk of nicotine dependence), across the 23 studies.

See attached spreadsheet for results.

**Supplementary Table 2.** Leave-one-study-out analyses of lead single nucleotide polymorphisms (SNPs) discovered in the cross-ancestry genome-wide association study (GWAS) meta-analysis of nicotine dependence (ND).

| Study (ancestry) removed | Remaining N | rs1862416-T association with ND | | | rs2714700-T association with ND | | |
|---|---|---|---|---|---|---|---|
| | | β | SE | P | β | SE | P |
| AAND (AA) | 56,313 | 0.039 | 0.0069 | $1.2\times10^{-8}$ | -0.023 | 0.0041 | $2.2\times10^{-8}$ |
| ADAA (AA) | 56,855 | 0.038 | 0.0069 | $2.6\times10^{-8}$ | -0.022 | 0.0041 | $6.6\times10^{-8}$ |
| COGEND (EUR) | 56,065 | 0.039 | 0.0069 | $2.5\times10^{-8}$ | -0.023 | 0.0041 | $3.4\times10^{-8}$ |
| COGEND (AA) | 57,296 | 0.039 | 0.0068 | $1.3\times10^{-8}$ | -0.022 | 0.0041 | $4.3\times10^{-8}$ |
| COGEND2 (EUR) | 57,708 | 0.038 | 0.0068 | $2.2\times10^{-8}$ | -0.022 | 0.0041 | $3.6\times10^{-8}$ |
| COGEND2 (AA) | 57,687 | 0.038 | 0.0068 | $2.2\times10^{-8}$ | -0.023 | 0.0041 | $1.4\times10^{-8}$ |
| COHRA1 (EUR) | 57,757 | 0.039 | 0.0068 | $1.2\times10^{-8}$ | -0.023 | 0.0041 | $2.3\times10^{-8}$ |
| COPDGene1 (EUR) | 55,451 | 0.038 | 0.0070 | $4.0\times10^{-8}$ | -0.024 | 0.0041 | $9.8\times10^{-9}$ |
| COPDGene1 (AA) | 55,466 | 0.038 | 0.0069 | $3.6\times10^{-8}$ | -0.024 | 0.0041 | $7.4\times10^{-9}$ |
| COPDGene2 (EUR) | 55,370 | 0.038 | 0.0070 | $5.3\times10^{-8}$ | -0.022 | 0.0042 | $1.2\times10^{-7}$ |
| COPDGene2 (AA) | 57,733 | 0.038 | 0.0068 | $2.8\times10^{-8}$ | -0.022 | 0.0041 | $3.2\times10^{-8}$ |
| deCODE (EUR) | 42,688 | 0.036 | 0.0078 | $3.9\times10^{-6}$ | -0.024 | 0.0047 | $3.0\times10^{-7}$ |
| EAGLE (EUR) | 54,994 | 0.039 | 0.0070 | $2.9\times10^{-8}$ | -0.021 | 0.0042 | $3.1\times10^{-7}$ |
| eMERGE (EUR) | 57,270 | 0.038 | 0.0070 | $6.2\times10^{-8}$ | -0.023 | 0.0042 | $7.4\times10^{-8}$ |
| FINRISK (EUR) | 55,789 | 0.039 | 0.0071 | $4.7\times10^{-8}$ | -0.022 | 0.0042 | $1.3\times10^{-7}$ |
| FTC (EUR) | 55,493 | 0.040 | 0.0070 | $7.2\times10^{-9}$ | -0.023 | 0.0042 | $5.6\times10^{-8}$ |
| GAIN (EUR) | 57,226 | 0.038 | 0.0068 | $2.0\times10^{-8}$ | -0.023 | 0.0041 | $1.6\times10^{-8}$ |

| | | | | | | | |
|---|---|---|---|---|---|---|---|
| GAIN (AA) | 57,523 | 0.039 | 0.0068 | $1.6\times10^{-8}$ | -0.023 | 0.0041 | $3.1\times10^{-8}$ |
| German (EUR) | 57,009 | 0.036 | 0.0069 | $1.3\times10^{-7}$ | -0.023 | 0.0041 | $1.7\times10^{-8}$ |
| JHS/ARIC (AA) | 56,857 | 0.039 | 0.0069 | $1.6\times10^{-8}$ | -0.022 | 0.0041 | $8.4\times10^{-8}$ |
| MCTFR (EUR) | 56,927 | 0.040 | 0.0069 | $7.2\times10^{-9}$ | -0.022 | 0.0041 | $4.9\times10^{-8}$ |
| nonGAIN (EUR) | 57,329 | 0.039 | 0.0068 | $1.3\times10^{-8}$ | -0.022 | 0.0041 | $7.3\times10^{-8}$ |
| NTR (EUR) | 53,511 | 0.040 | 0.0070 | $1.0\times10^{-8}$ | -0.023 | 0.0042 | $4.0\times10^{-8}$ |
| OZ-ALC (EUR) | 56,862 | 0.038 | 0.0070 | $6.6\times10^{-8}$ | -0.023 | 0.0041 | $2.7\times10^{-8}$ |
| SAGE (EUR) | 57,168 | 0.038 | 0.0069 | $2.5\times10^{-8}$ | -0.023 | 0.0041 | $3.0\times10^{-8}$ |
| SAGE (AA) | 57,367 | 0.039 | 0.0068 | $1.1\times10^{-8}$ | -0.022 | 0.0041 | $3.7\times10^{-8}$ |
| Spit for Science (EUR) | 56,283 | 0.039 | 0.0072 | $6.0\times10^{-8}$ | -0.022 | 0.0043 | $2.1\times10^{-7}$ |
| UW-TTURC (EUR) | 56,466 | 0.040 | 0.0069 | $1.1\times10^{-8}$ | -0.023 | 0.0041 | $4.1\times10^{-8}$ |
| UW-TTURC (AA) | 57,753 | 0.038 | 0.0068 | $2.2\times10^{-8}$ | -0.023 | 0.0041 | $2.1\times10^{-8}$ |
| Yale-Penn (EUR) | 56,421 | 0.040 | 0.0069 | $5.6\times10^{-9}$ | -0.023 | 0.0041 | $1.9\times10^{-8}$ |
| Yale-Penn (AA) | 55,363 | 0.038 | 0.0069 | $4.4\times10^{-8}$ | -0.022 | 0.0042 | $1.2\times10^{-7}$ |

Abbreviations: AA, African American ancestry; EUR, European ancestry; SE, standard error.

**Supplementary Table 3**. Characteristics of participants included in the genome-wide association study (GWAS) meta-analyses of nicotine dependence (ND), separated into each of the 23 studies and the two ancestry groups.

| Study | Total N | N (%), females | Mean age (SD) | European ancestry (total N=46,213) | | | | African American ancestry (total N=11,787) | | | |
|---|---|---|---|---|---|---|---|---|---|---|---|
| | | | | N (%), mild ND | N (%), moderate ND | N (%), severe ND | GWAS λ | N (%), mild ND | N (%), moderate ND | N (%), severe ND | GWAS λ |
| AAND | 1,687 | 969 (57.4) | 41.2 (10.3) | NA | NA | NA | NA | 526 (31.2) | 830 (49.2) | 331 (19.6) | 1.00 |
| ADAA | 1,145 | 472 (41.2) | 41.2 (10.3) | NA | NA | NA | NA | 526 (31.2) | 830 (49.2) | 331 (19.6) | 1.01 |
| COGEND [a] | 2,639 | 1,628 (61.7) | 36.6 (5.57) | 941 (48.6) | 521 (26.9) | 473 (24.4) | 1.01 | 248 (35.2) | 283 (40.2) | 173 (24.6) | 1.01 |
| COGEND2 | 605 | 324 (53.6) | 34.4 (5.87) | 60 (20.5) | 91 (31.2) | 141 (48.3) | 1.01 | 13 (4.2) | 137 (43.8) | 163 (52.1) | 1.03 |
| COHRA1 | 243 | 129 (53.1) | 32.1 (9.1) | 79 (32.5) | 127 (52.3) | 37 (15.2) | 1.01 | NA | NA | NA | NA |
| COPDGene1 [a] | 5,083 | 2,817 (55.4) | 55.4 (7.3) | 743 (29.1) | 1,118 (43.9) | 688 (27.0) | 1.03 | 711 (28.1) | 1,149 (45.3) | 674 (26.6) | 1.00 |

| | | | | | | | | | | | |
|---|---|---|---|---|---|---|---|---|---|---|---|
| COPDGene2 [a] | 2,897 | 1,395 (48.2) | 63.7 (8.0) | 955 (36.3) | 1,172 (44.6) | 503 (19.1) | 1.03 | 146 (54.7) | 103 (38.6) | 18 (6.7) | 1.00 |
| deCODE [a,b] | 15,312 | 9,127 (59.6) | 66.5 (15.4) | 11,494 (75.1) | 2,250 (16.6) | 1,268 (8.3) | 1.12 | NA | NA | NA | NA |
| EAGLE | 3,006 | 478 (15.9) | NA [c] | 1,416 (47.1) | 1,027 (34.2) | 563 (18.7) | 1.00 | NA | NA | NA | NA |
| eMERGE [a] | 730 | 319 (43.7) | 72.1 (9.2) | 487 (66.7) | 193 (26.4) | 50 (6.8) | 1.02 | NA | NA | NA | NA |
| FINRISK | 2,211 | 1,025 (46.4) | 50.5 (13.3) | 1,401 (63.4) | 614 (27.8) | 196 (8.9) | 1.02 | NA | NA | NA | NA |
| FTC [a] | 2,507 | 1,111 (44.3) | 45.5 (16.2) | 1,436 (57.3) | 828 (33.0) | 243 (9.7) | 1.00 | NA | NA | NA | NA |
| GAIN | 1,251 | 655 (52.4) | 52.0 (15.2) | 327 (42.2) | 280 (36.2) | 167 (21.6) | 1.01 | 221 (46.3) | 176 (36.9) | 80 (16.8) | 0.99 |
| German | 991 | 543 (54.8) | 36.3 (12.6) | 565 (57.0) | 313 (31.6) | 113 (11.4) | 1.00 | NA | NA | NA | NA |
| JHS/ARIC [b] | 1,143 | 641 (56.1) | 52.9 (9.2) | NA | NA | NA | NA | 867 (75.9) | 218 (19.1) | 58 (5.1) | 1.01 |
| MCTFR [a] | 1,073 | 492 (45.9) | 20.6 (5.4) | 687 (64.0) | 293 (27.3) | 93 (8.7) | 1.01 | NA | NA | NA | NA |
| nonGAIN | 671 | 322 (48.0) | 52.9 (15.5) | 298 (44.4) | 234 (34.9) | 139 (20.7) | 1.02 | NA | NA | NA | NA |

| | | | | | | | | | | | |
|---|---|---|---|---|---|---|---|---|---|---|---|
| NTR [a] | 4,489 | 2,750 (61.3) | 45.5 (15.0) | 2,842 (63.3) | 1,276 (28.4) | 371 (8.3) | 1.01 | NA | NA | NA | NA |
| OZ-ALC | 1,138 | 379 (33.3) | 45.6 (9.3) | 976 (85.8) | 125 (11.0) | 37 (3.3) | 1.01 | NA | NA | NA | NA |
| SAGE | 1,465 | 649 (44.3) | 40.9 (9.9) | 243 (29.2) | 295 (35.5) | 294 (35.3) | 1.01 | 211 (33.3) | 272 (43.0) | 150 (23.7) | 1.00 |
| Spit for Science | 1,717 | 994 (57.9) | 20.4 (1.5) | 1,532 (89.2) | 158 (9.2) | 33 (1.9) | 0.99 | NA | NA | NA | NA |
| UW-TTURC | 1,781 | 1,040 (58.4) | 43.4 (11.2) | 311 (20.3) | 723 (47.1) | 500 (32.6) | 1.01 | 40 (16.2) | 119 (48.2) | 88 (35.6) | 1.01 |
| Yale-Penn | 4,216 | 1,833 (43.5) | 40.1 (9.42) | 284 (18.0) | 751 (47.6) | 544 (34.4) | 1.01 | 837 (31.7) | 1,346 (51.0) | 454 (17.2) | 1.04 |

Abbreviations: NA, not available; SD, standard deviation

[a] European ancestry participants were included in the GWAS and Sequencing Consortium of Alcohol and Nicotine use (GSCAN) consortium.[48] [b] ND was categorized according to Fagerström Test for Nicotine Dependence (FTND) scores: mild (FTND score 0–3), moderate (FTND score 4–6) or severe (FTND score 7–10). For deCODE and JHS/ARIC only, the mild category included participants with FTND score 0–3 as well as low-intensity smokers with no FTND data available but with ≤10 cigarettes per day (CPD). [c] Age was only available as a categorical variable: 23.2% aged 59 or less, 18.2% aged 60–64, 22.4% aged 65–69, 21.4% aged 70–74 and 14.8% aged 75–79.

**Supplementary Table 4.** Associations of the novel nicotine dependence (ND)-implicated single nucleotide polymorphisms (SNPs) with other smoking traits in the GWAS and Sequencing Consortium of Alcohol and Nicotine use (GSCAN) consortium[48]—initiation (ever vs. never smoking), age at initiation, cigarettes per day, and cessation (current vs. former smoking).

SNP associations at $P<0.05$ are bolded.

| SNP (effect allele) | Initiation (N=1,232,091) | | Age at initiation (N=341,427) | | Cigarettes per day (N=337,334) | | Cessation (N=547,219) | |
|---|---|---|---|---|---|---|---|---|
| | **β (SE)** | **P** | **β (SE)** | **P** | **β (SE)** | **P** | **β (SE)** | **P** |
| rs1862416 (T) | **0.005 (0.003)** | **0.033** | -0.01 (0.005) | 0.080 | 0.001 (0.003) | 0.61 | $-6.6\times10^{-5}$ (0.004) | 0.50 |
| rs2714700 (T) | **-0.003 (0.002)** | **0.016** | $-4\times10^{-4}$ (0.003) | 0.80 | **-0.004 (0.002)** | **0.045** | -0.001 (0.002) | 0.31 |

Abbreviation: SE, standard error.

**Supplementary Table 5.** Linkage disequilibrium (LD) structure and conditional association testing of the nicotine dependence (ND)-associated *TENM2* single nucleotide polymorphism (SNP) rs1862416 (chr. 5: 167,394,595) and the nearby lead SNPs implicated at genome-wide statistical significance for smoking initiation (ever vs. never smoking) by the GWAS and Sequencing Consortium of Alcohol and Nicotine use (GSCAN) consortium.[48]

LD was determined using the LDlink tool,[51] and conditional modeling was conducted using ND GWAS meta-analysis summary statistics as input into the Genome-wide Complex Trait Analysis (GCTA) tool.[52,53]

| GSCAN lead SNP | chr. 5 position (NCBI build 37) | P, cross-ancestry meta-analysis for ND | LD in 1000G European panel | | LD in 1000G African panel | | rs1862416 associations with ND conditioned on GSCAN lead SNP(s) | | |
|---|---|---|---|---|---|---|---|---|---|
| | | | $r^2$ | D’ | $r^2$ | D’ | P, European ancestry-specific meta-analysis | P, African American-specific meta-analysis | P, cross ancestry meta-analysis |
| rs3909281 | 165,096,435 | 0.12 | 0.0020 | 0.11 | 0.00090 | 0.041 | $7.5\times10^{-7}$ | $7.0\times10^{-3}$ | $2.2\times10^{-8}$ |
| rs3843905 | 165,427,280 | 0.55 | 0.00010 | 0.025 | 0.00080 | 0.065 | $6.2\times10^{-7}$ | $7.3\times10^{-3}$ | $1.8\times10^{-8}$ |
| rs79476395 | 166,063,680 | 0.018 | 0.00030 | 0.20 | 0.0014 | 0.46 | $1.6\times10^{-6}$ | $6.7\times10^{-3}$ | $4.6\times10^{-8}$ |
| rs6890961 | 166,778,503 | $9.9\times10^{-4}$ | 0.0047 | 0.24 | 0.00030 | 0.048 | $2.9\times10^{-6}$ | $6.1\times10^{-3}$ | $7.9\times10^{-8}$ |
| rs4044321 | 166,989,513 | 0.78 | 0.00040 | 0.040 | 0.00050 | 0.072 | $6.7\times10^{-7}$ | $6.9\times10^{-3}$ | $1.9\times10^{-8}$ |
| rs2173019 | 167,614,971 | 0.036 | 0.0017 | 0.024 | 0.0017 | 0.16 | $1.0\times10^{-6}$ | $6.0\times10^{-3}$ | $2.7\times10^{-8}$ |
| | | | | | | All SNPs | $7.2\times10^{-6}$ | $6.6\times10^{-3}$ | $2.2\times10^{-7}$ |

Abbreviations: 1000G, 1000 Genomes; NCBI, National Center for Biotechnology Information.

**Supplementary Table 6**. Independent testing of novel nicotine dependence (ND)-associated single nucleotide polymorphisms (SNPs) with heaviness of smoking (HSI) in the UK Biobank.

| SNP (effect allele) | Chr:position (NCBI build 37) | Gene / closest genes | HSI in UK Biobank (N=33,791) | | | | Meta-analysis of FTND GWAS and UK Biobank HSI results (total N=91,791) | |
|---|---|---|---|---|---|---|---|---|
| | | | Effect allele freq. | estimated $r^2$ | β (SE) | P | β (SE) | P |
| rs1862416 (T) | 5:167,394,595 | *TENM2* | 0.89 | 1 | -0.0064 (0.0075) | 0.39 | 0.018 (0.0050) | $3.0 \times 10^{-4}$ |
| rs2714700 (T) | 7:79,367,667 | *MAGI2 / GNAI1* | 0.47 | 1 | -0.012 (0.0047) | 0.014 | -0.018 (0.0031) | $7.7 \times 10^{-9}$ |

Abbreviations: FTND, Fagerström Test for Nicotine Dependence (FTND); GWAS, genome-wide association study; NCBI, National Center for Biotechnology Information; SE, standard error.

**Supplementary Table 7.** Genome-wide H-MAGMA results using the nicotine dependence GWAS meta-analysis of European ancestry participants in the iNDiGO consortium with reference to chromatin interaction maps from fetal brain tissue.

Results are sorted by the H-MAGMA P value.

See the attached spreadsheet for results.

**Supplementary Table 8.** Genome-wide H-MAGMA results using the nicotine dependence GWAS meta-analysis of European ancestry participants in the iNDiGO consortium with reference to chromatin interaction maps from adult brain tissue.

Results are sorted by the H-MAGMA P value.

See the attached spreadsheet for results.

**Supplementary Table 9**. Statistically significant H-MAGMA results from the nicotine dependence GWAS meta-analysis of European ancestry participants in the iNDiGO consortium, based on $P<2.7\times10^{-6}$ (Bonferroni correction for testing 18,655 genes), with look-up in the UK Biobank using heaviness of smoking index as a proxy for nicotine dependence.

H-MAGMA was applied in both the iNDiGO consortium and the UK Biobank using chromatin interaction maps in fetal and adult brain tissues, separately, as the reference datasets. Results are sorted, within each tissue, by H-MAGMA p-values in the iNDiGO consortium. H-MAGMA p-values in the UK Biobank that surpass Bonferroni correction for testing 16 unique genes (P<0.0031) are shown in bold.

| Gene | Chr. band | iNDiGO (N=46,213) | | UK Biobank (N=33,791) | |
|---|---|---|---|---|---|
| | | No. SNPs annotated to gene | P | No. SNPs annotated to gene | P |
| ***Fetal brain tissue as chromatin interaction mapping reference in H-MAGMA*** | | | | | |
| *CHRNA5* | 15q25 | 17 | $2.6\times10^{-28}$ | 19 | $\mathbf{8.9\times10^{-26}}$ |
| *IREB2* | 15q25 | 40 | $1.7\times10^{-27}$ | 42 | $\mathbf{3.3\times10^{-22}}$ |
| *HYKK* | 15q25 | 16 | $2.4\times10^{-27}$ | 20 | $\mathbf{4.8\times10^{-25}}$ |
| *CHRNA3* | 15q25 | 82 | $6.4\times10^{-24}$ | 84 | $\mathbf{5.7\times10^{-25}}$ |
| *CHRNB4* | 15q25 | 93 | $1.8\times10^{-14}$ | 101 | $\mathbf{2.3\times10^{-15}}$ |
| *ADAMTS7* | 15q25 | 30 | $1.6\times10^{-12}$ | 32 | $\mathbf{2.9\times10^{-14}}$ |
| *CHRNA4* | 20q13 | 264 | $7.7\times10^{-12}$ | 282 | $1.0\times10^{-2}$ |
| *PSMA4* | 15q25 | 52 | $2.3\times10^{-11}$ | 58 | $\mathbf{1.3\times10^{-10}}$ |
| *MORF4L1* | 15q25 | 60 | $2.9\times10^{-11}$ | 63 | $\mathbf{1.7\times10^{-11}}$ |
| *ADAMTSL2* | 9q34 | 277 | $3.4\times10^{-8}$ | 296 | $\mathbf{3.2\times10^{-8}}$ |
| *DBH* | 9q34 | 114 | $1.7\times10^{-6}$ | 127 | $\mathbf{1.1\times10^{-4}}$ |
| ***Adult brain tissue as chromatin interaction mapping reference in H-MAGMA*** | | | | | |
| *CHRNA5* | 15q25 | 17 | $2.6\times10^{-28}$ | 19 | $\mathbf{8.9\times10^{-26}}$ |
| *WDR61* | 15q25 | 31 | $3.5\times10^{-22}$ | 32 | $\mathbf{1.8\times10^{-20}}$ |
| *IREB2* | 15q25 | 130 | $4.2\times10^{-18}$ | 139 | $\mathbf{2.1\times10^{-13}}$ |
| *CHRNA3* | 15q25 | 101 | $5.4\times10^{-15}$ | 115 | $\mathbf{6.6\times10^{-16}}$ |
| *HYKK* | 15q25 | 143 | $2.2\times10^{-14}$ | 158 | $\mathbf{2.1\times10^{-14}}$ |
| *ACSBG1* | 15q25 | 117 | $8.0\times10^{-14}$ | 126 | $\mathbf{9.4\times10^{-13}}$ |
| *ADAMTS7* | 15q25 | 71 | $2.3\times10^{-11}$ | 76 | $\mathbf{2.0\times10^{-13}}$ |
| *PSMA4* | 15q25 | 52 | $2.3\times10^{-11}$ | 58 | $\mathbf{1.3\times10^{-10}}$ |
| *CHRNA4* | 20q13 | 96 | $1.0\times10^{-10}$ | 99 | $\mathbf{7.0\times10^{-5}}$ |
| *CHRNB4* | 15q25 | 53 | $1.7\times10^{-9}$ | 59 | $\mathbf{1.7\times10^{-10}}$ |
| *AFG1L* | 6q21 | 281 | $1.1\times10^{-6}$ | 319 | 0.54 |
| *AK2* | 1p35 | 60 | $1.3\times10^{-6}$ | 68 | 0.38 |
| *RBBP8NL* | 20q13 | 172 | $2.1\times10^{-6}$ | 178 | $8.6\times10^{-3}$ |

**Supplementary Table 10.** Genome-wide Summary-MultiXcan (S-MultiXcan)[54] results from the European ancestry-specific nicotine dependence GWAS meta-analysis summary statistics with reference to imputed genetically driven gene expression across the 13 adult brain tissues in GTEx.

S-MultiXcan provides gene-level association results based on aggregating *cis*-eQTL evidence across multiple tissues, while also presenting gene-based results from the best and worst single-tissue models. Results are sorted by the multi-tissue P value.

See attached spreadsheet for results.

### **Supplementary Table 11.** Genetic correlations of nicotine dependence (ND) with other phenotypes using linkage disequilibrium (LD) score regression (LDSC).

Phenotypes are sorted by disease or measurement category. Phenotypes that have statistically significant correlations with ND, as determined by Bonferroni correction ($\alpha$=0.05/46 phenotypes, $P_1$<0.0011), are bolded.

| **Category** | **Phenotype** | **Reference** | **$h^2$ (single trait SNP heritability)** | **Cross-trait comparison with ND** | | | | |
|---|---|---|---|---|---|---|---|---|
| | | | | **gcov_int[a]** | **$r_g$** | **SE** | **$P_1$ ($H_0$: $r_g$ = 0)** | **$P_2$ ($H_0$: $r_g$ = 1)** |
| Brain volume | Accumbens volume | 55 | 0.092 | 0.0020 | 0.13 | 0.15 | 0.37 | $6.5\times10^{-9}$ |
| | Caudate volume | 55 | 0.25 | 0.0067 | -0.091 | 0.094 | 0.33 | $3.4\times10^{-22}$ |
| | Hippocampus volume | 55 | 0.14 | 0.0062 | -0.15 | 0.13 | 0.25 | $2.2\times10^{-11}$ |
| | Intracranial volume | 55 | 0.18 | 0.00040 | -0.24 | 0.12 | 0.036 | $5.2\times10^{-11}$ |
| | Pallidum volume | 55 | 0.16 | 0.0050 | -0.075 | 0.11 | 0.50 | $2.0\times10^{-16}$ |
| | Putamen volume | 55 | 0.30 | -0.0010 | 0.17 | 0.083 | 0.045 | $8.8\times10^{-24}$ |
| | Thalamus volume | 55 | 0.14 | -0.0017 | -0.092 | 0.12 | 0.43 | $1.1\times10^{-14}$ |
| Cancer | **Lung adenocarcinoma** | **56** | **0.069** | **0.0037** | **0.48** | **0.11** | **$8.6\times10^{-6}$** | **$1.1\times10^{-6}$** |
| | **Lung cancer (overall)** | **56** | **0.087** | **0.0065** | **0.68** | **0.089** | **$3.4\times10^{-14}$** | **$2.9\times10^{-4}$** |
| | Small cell lung cancer | 56 | 0.11 | 0.0085 | 0.40 | 0.13 | 0.0024 | $7.5\times10^{-6}$ |
| | **Squamous cell lung cancer** | **56** | **0.053** | **0.0065** | **0.75** | **0.11** | **$3.0\times10^{-11}$** | **0.03** |
| Cardiometabolic | Adiponectin | 57 | 0.12 | -0.0051 | 0.035 | 0.11 | 0.74 | $2.6\times10^{-20}$ |
| | **Coronary artery disease** | **58 b** | **0.080** | **-0.0032** | **0.32** | **0.064** | **$6.0\times10^{-7}$** | **$4.6\times10^{-26}$** |
| Cigarette smoking | **Age of smoking initiation** | **48** | **0.047** | **-0.042** | **-0.55** | **0.066** | **$1.7\times10^{-16}$** | **$8.9\times10^{-12}$** |

| | | | | | | | | |
|---|---|---|---|---|---|---|---|---|
| | **Cigarettes per day** | 48 | **0.075** | **0.14** | **0.95** | **0.054** | **$3.1\times10^{-70}$** | **0.35** |
| | Cotinine levels | 59 | 0.22 | 0.020 | 0.46 | 0.23 | 0.051 | 0.021 |
| | **Smoking cessation (current vs. former)** | 48 | **0.032** | **0.032** | **0.51** | **0.063** | **$3.4\times10^{-16}$** | **$8.2\times10^{-15}$** |
| | **Smoking initiation (ever vs. never)** | 48 | **0.069** | **0.012** | **0.40** | **0.049** | **$3.2\times10^{-16}$** | **$2.8\times10^{-34}$** |
| Cognitive / education | Childhood IQ | 60 | 0.28 | 0.00080 | -0.17 | 0.10 | 0.11 | $2.1\times10^{-15}$ |
| | College completion | 61 | 0.079 | -0.024 | -0.23 | 0.070 | 0.0012 | $4.3\times10^{-28}$ |
| | Intelligence | 62 | 0.19 | 0.0024 | -0.17 | 0.056 | 0.0031 | $1.3\times10^{-50}$ |
| | **Years of schooling** | 63 c | **0.11** | **-0.012** | **-0.34** | **0.041** | **$9.2\times10^{-17}$** | **$8.6\times10^{-58}$** |
| Drug and alcohol | **Alcohol dependence** | 64 | **0.096** | **0.025** | **0.57** | **0.13** | **$6.3\times10^{-6}$** | **$1.4\times10^{-4}$** |
| | Alcohol drinks per week | 48 | 0.049 | 0.017 | 0.13 | 0.054 | 0.016 | $1.4\times10^{-58}$ |
| | Cannabis use disorder | 65 | 0.027 | 0.0029 | 0.40 | 0.15 | 0.010 | $9.4\times10^{-5}$ |
| | Lifetime cannabis use (ever vs. never) | 66 | 0.067 | -0.0057 | 0.057 | 0.056 | 0.31 | $3.9\times10^{-63}$ |
| Neurologic | Alzheimer's disease | 67 | 0.045 | -0.0043 | -0.087 | 0.12 | 0.48 | $1.5\times10^{-13}$ |
| | Amyotrophic lateral sclerosis | 68 | 0.049 | 0.0010 | -0.060 | 0.12 | 0.62 | $1.3\times10^{-14}$ |
| | Parkinson's disease | 69 | 0.41 | $6.3\times10^{-7}$ | 0.074 | 0.092 | 0.42 | $9.2\times10^{-24}$ |
| Personality | Conscientiousness | 70 | 0.073 | -0.014 | 0.052 | 0.18 | 0.77 | $1.6\times10^{-7}$ |
| | **Neuroticism** | 71 | **0.089** | **0.0054** | **0.28** | **0.067** | **$3.2\times10^{-5}$** | **$1.4\times10^{-26}$** |
| | Openness to experience | 70 | 0.11 | -0.0077 | -0.12 | 0.13 | 0.35 | $2.8\times10^{-12}$ |
| Psychiatric | Anorexia nervosa | 72 | 0.18 | -0.014 | 0.098 | 0.066 | 0.14 | $5.0\times10^{-43}$ |

| | | | | | | | | |
|---|---|---|---|---|---|---|---|---|
| | **Attention deficit hyperactivity disorder** | [73] | **0.24** | **-0.0033** | **0.49** | **0.063** | **$5.7\times10^{-15}$** | **$9.1\times10^{-16}$** |
| | Autism spectrum disorder | [74] | 0.20 | -0.0068 | 0.23 | 0.078 | 0.0024 | $2.7\times10^{-24}$ |
| | **Bipolar disorder** | [75] | **0.35** | **-0.0024** | **0.25** | **0.050** | **$3.3\times10^{-7}$** | **$2.6\times10^{-52}$** |
| | **Depressive symptoms** | [71] | **0.047** | **-0.00080** | **0.40** | **0.075** | **$9.6\times10^{-8}$** | **$1.1\times10^{-15}$** |
| | **Major depressive disorder** | [76] | **0.038** | **-0.0046** | **0.38** | **0.051** | **$6.1\times10^{-14}$** | **$1.7\times10^{-33}$** |
| | **Posttraumatic stress disorder** | [77] | **0.017** | **-0.0077** | **0.72** | **0.15** | **$6.5\times10^{-7}$** | **0.056** |
| | Psychiatric cross-disorder | [78] | 0.17 | 0.0021 | 0.17 | 0.080 | 0.031 | $4.6\times10^{-25}$ |
| | **Schizophrenia** | [79] | **0.46** | **0.0077** | **0.18** | **0.043** | **$3.2\times10^{-5}$** | **$1.3\times10^{-65}$** |
| | Subjective well being | [71] | 0.025 | -0.0092 | -0.24 | 0.075 | 0.0016 | $9.4\times10^{-25}$ |
| Respiratory | COPD | [80] | 0.10 | -0.0065 | 0.18 | 0.088 | 0.033 | $1.8\times10^{-20}$ |
| | Forced expiratory volume in 1 second ($FEV_1$) | [81] | 0.27 | -0.0048 | -0.0017 | 0.060 | 0.98 | $1.4\times10^{-63}$ |
| | Forced vital capacity (FVC) | [81] | 0.26 | -0.0054 | -0.0073 | 0.057 | 0.90 | $4.2\times10^{-69}$ |
| | $FEV_1$/FVC | [81] | 0.26 | -0.0020 | 0.012 | 0.059 | 0.84 | $6.1\times10^{-63}$ |

[a] Deviation of the cross-trait intercept term from 0 is indicative of study overlap in the GWAS results being compared.

[b] Results are based on cross-ancestry meta-analysis results that are available in LDHub; results for all other results correspond to European-specific meta-analyses.

[c] The GWAS results for educational attainment (years of schooling) include all discovery cohorts, except for 23andMe, resulting in a total sample size of 766,345.

### **Supplementary Table 12**. Credible set analysis and annotation of the novel nicotine dependence (ND)-associated loci.

For the novel ND-associated loci (chromosomes 5q34 and 7q21), we applied a Bayesian method[82] implemented via LocusZoom[83] to identify a credible set likely to contain the causal variant at each loci. The calculated posterior probability for each variant is provided as well as the cross-ancestry, European ancestry-specific, and African American ancestry-specific meta-analysis results for comparison. HetPval is the heterogeneity p-value from the meta-analysis. The credible set was annotated using GTEx,[84] BrainSeq,[85] and HaploReg.[86]

See attached spreadsheet for results.

### **Supplementary Table 13.** Tissues and cell types evaluated for shared genetics with nicotine dependence (ND) using stratified linkage disequilibrium (LD) score regression, as applied to specifically expressed genes (LDSC-SEG).

Tissues and cell types are sorted by data origin (RNA-sequencing in the Genotype-Tissue Expression [GTEx] project or array-based in Gene Expression Omnibus [GEO]) and then by p-value. Tissues/cell types that have statistically significant correlations with ND, as determined by Bonferroni correction ($\alpha$=0.05/205 phenotypes, $P<2.4\times10^{-4}$), are bolded.

See attached spreadsheet for results.

### **Supplementary Table 14.** Look-up of genome-wide significant single nucleotide polymorphisms (SNPs) in the GWAS and Sequencing Consortium of Alcohol and Nicotine use (GSCAN) consortium[48] for association with nicotine dependence (ND) in our cross-ancestry GWAS meta-analysis.

Results are sorted by GSCAN smoking phenotype in descending $|r_g|$ with ND (cigarettes per day [$r_g$=0.95], age at initiation [$r_g$=-0.55], cessation [current vs. former smoking, $r_g$=0.51], and initiation [ever vs. never smoking, $r_g$=0.40], shown in alternating grey shading) and then by FTND GWAS meta-analysis p-value. For SNPs implicated at genome-wide significance for more than one phenotype in GSCAN, the results from the phenotype with the smallest p-value are presented. $\beta$ estimates correspond to the effect alleles. No SNPs from novel loci for ND surpassed Bonferroni correction for multiple testing ($P<1.1\times10^{-4}$, $\alpha$=0.05/452 SNPs available in the iNDiGO cross-ancestry FTND GWAS meta-analysis). Abbreviations: AI, age at initiation; CPD, cigarettes per day; NA, not available; SC, smoking cessation; SE, standard error; and SI, smoking initiation.

See attached spreadsheet for results.

**Supplementary Table 15.** Agreement between heaviness of smoking index (HSI) and Fagerström Test for Nicotine Dependence (FTND) categories for mild, moderate, and severe nicotine dependence (ND) in COGEND participants of European ancestry.

| | | FTND category[a], N (% of total N in HSI category) | | | Total |
|---|---|---|---|---|---|
| | | ***Mild*** | ***Moderate*** | ***Severe*** | |
| **HSI category**[b] | ***Mild*** | 998 (94.9) | 54 (5.1) | 0 (0) | 1,052 |
| | ***Moderate*** | 3 (0.6) | 417 (81.9) | 89 (17.5) | 509 |
| | ***Severe*** | 0 (0) | 72 (15.1) | 404 (84.9) | 476 |

[a] Categories for the full-scale, 6-item FTND (score range = 0–10) were defined as follows: mild (scores 0–3), moderate (scores 4–6), or severe (scores 7–10).

[b] Categories for the 2-item HSI (score range = 0–6) were defined as follows: mild (scores 0–2), moderate (scores 3–4), or severe (scores 5–6).

**Supplementary Figure 1**. Quantile-quantile plots for nicotine dependence genome-wide association study (GWAS) meta-analyses.

Results are shown for (A) the cross-ancestry meta-analysis (European ancestry and African American participants from all studies), (B) the European ancestry-specific meta-analysis, and (C) African American-specific meta-analysis. The observed vs expected meta-analysis $-\log_{10}$ p-values (black dots) are plotted along the identity line (red) with the corresponding genomic inflation factor ($\lambda$) indicated.

(A)

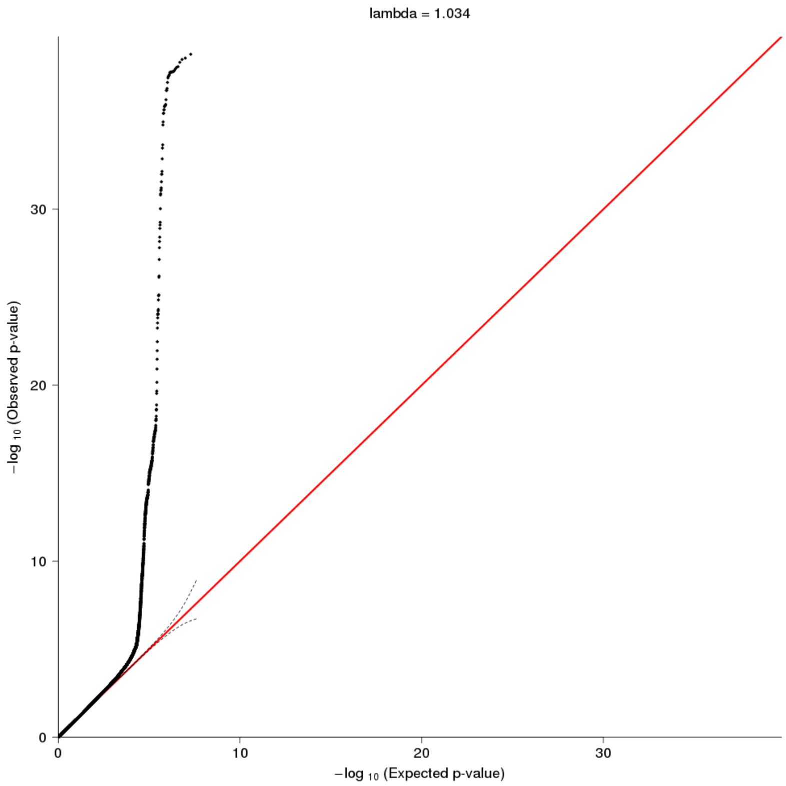

(B)

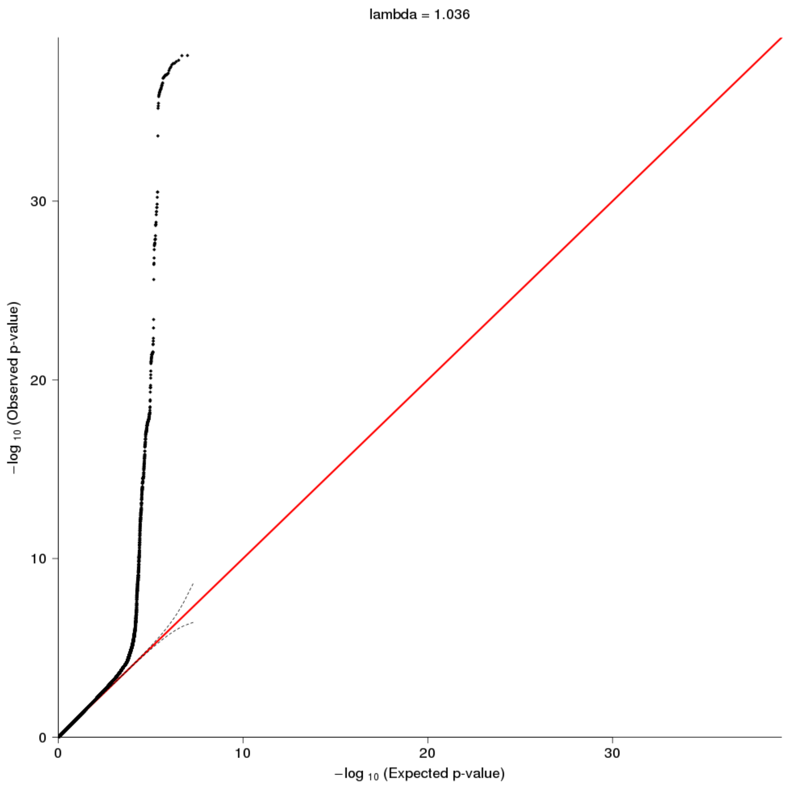
lambda = 1.036
−log 10 (Observed p-value)
−log 10 (Expected p-value)
0
10
20
30
0
10
20
30

(C)

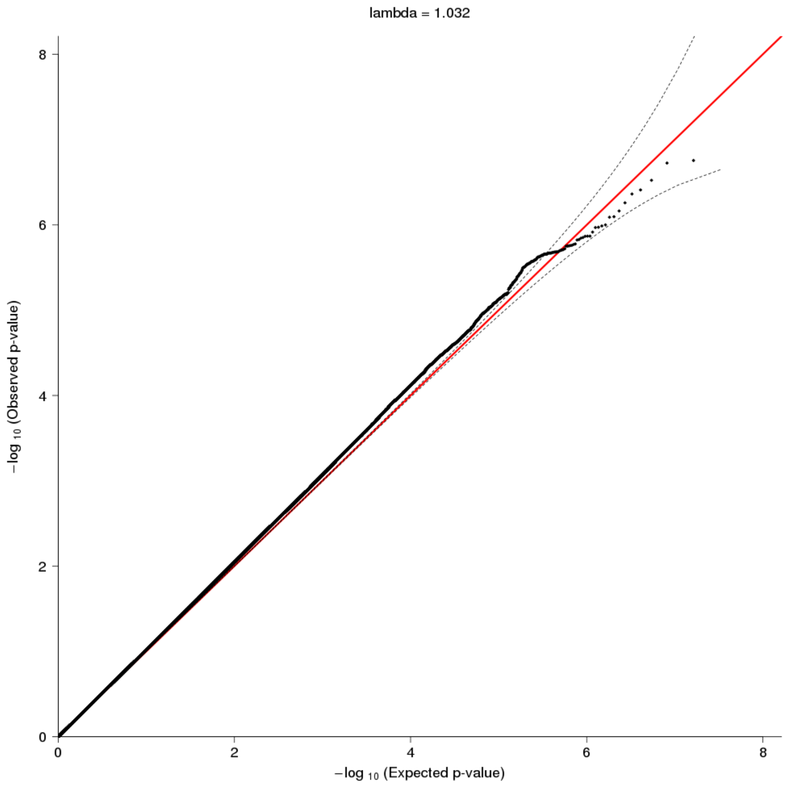
lambda = 1.032
−log 10 (Observed p-value)
−log 10 (Expected p-value)
0
2
4
6
8
0
2
4
6
8

**Supplementary Figure 2**. Manhattan plots for ancestry-specific nicotine dependence genome-wide association study (GWAS) meta-analyses.

Results are shown for (A) the European ancestry-specific (total N=46,213) and (B) African American-specific (total N=11,787) meta-analyses. The $-\log_{10}$ meta-analysis p-values are plotted by chromosomal position of single nucleotide polymorphisms (SNPs; depicted as circles) and insertions/deletions (indels; depicted as triangles). The genome-wide statistical significance threshold ($P<5\times10^{-8}$) is shown as a solid black line.

(A)

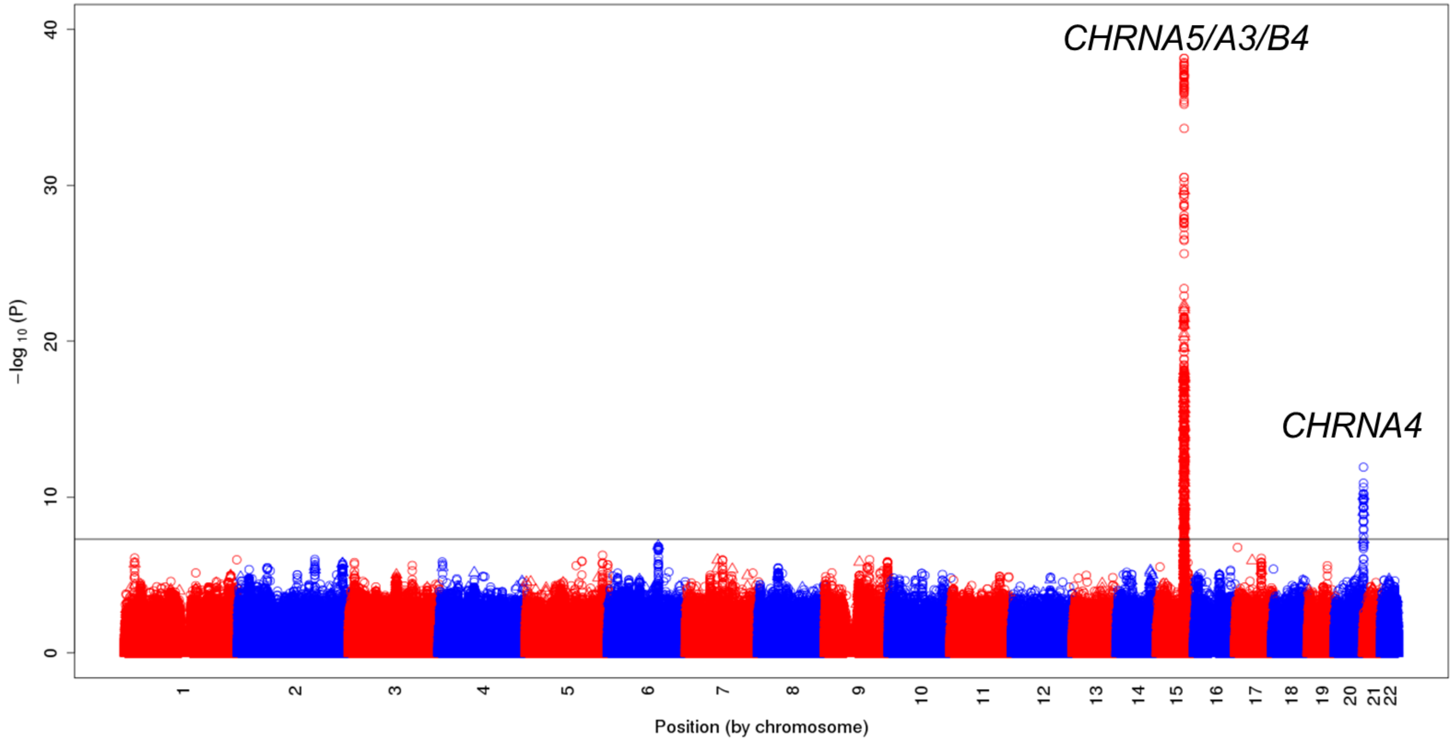

(B)

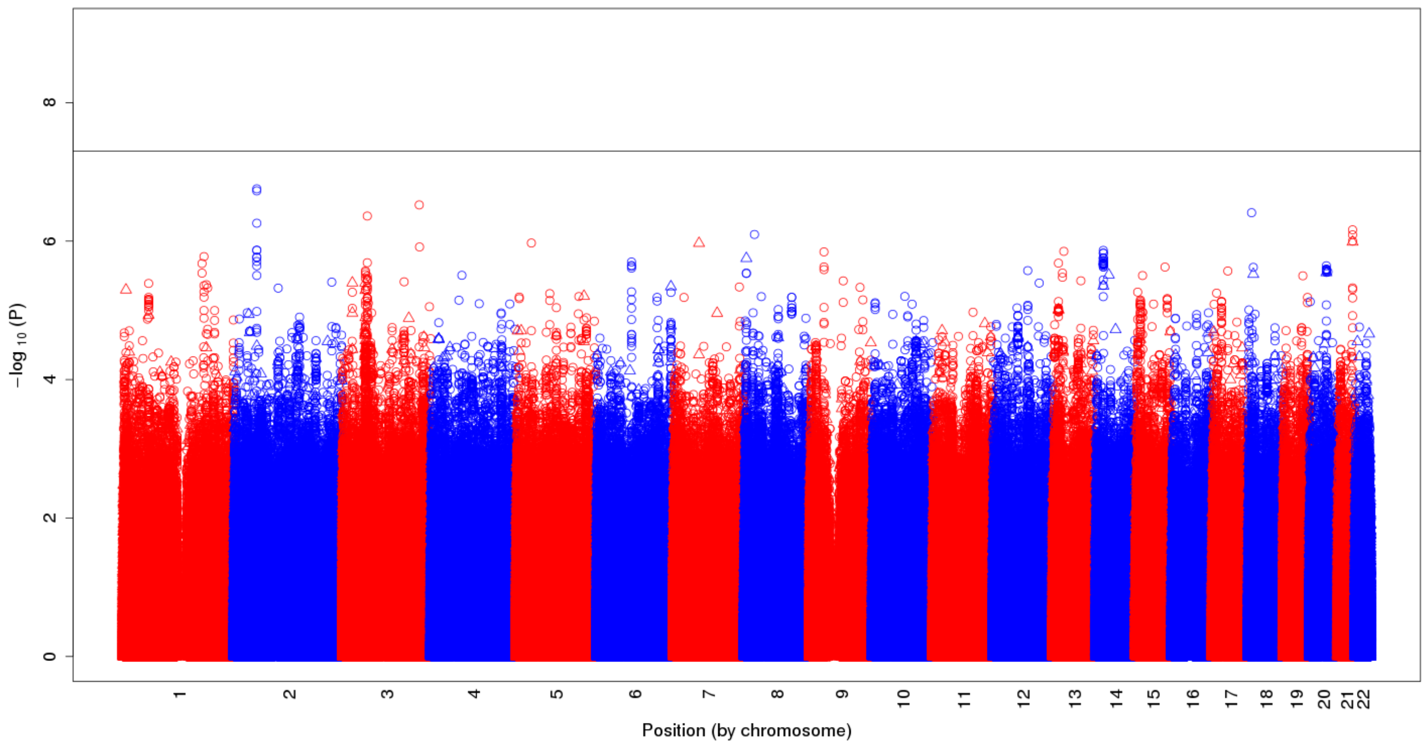

−log 10 (P)
0
2
4
6
8
Position (by chromosome)

**Supplementary Figure 3**. Regional association plots for two novel loci identified at genome-wide significance in the cross-ancestry nicotine dependence genome-wide association study (GWAS) meta-analyses.

Results are shown for rs2714700 on chromosome 7 in reference to the 1000 Genomes (A) European and (B) African superpopulation panels and rs1862416 on chromosome 5in reference to the same (C) European and (D) African panels. The $-\log_{10}$ meta-analysis p-values are plotted by chromosomal position with $r^2$ values between the lead single nucleotide polymorphism (SNP; in purple) and nearby SNPs indicated in 0.2 increments (e.g., 0.8–1 in red).

(A)

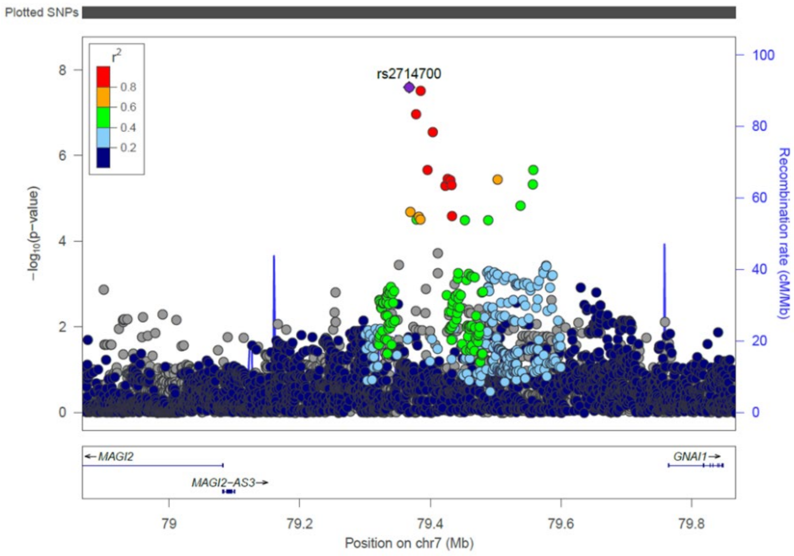

(B)

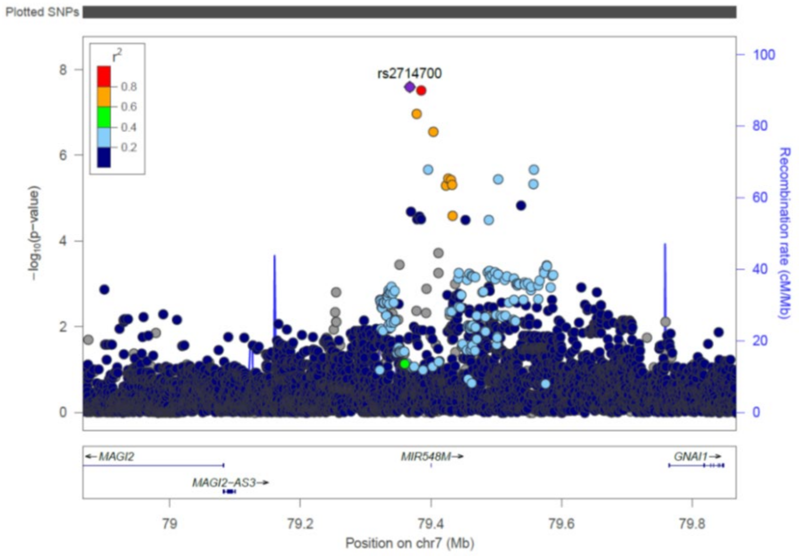

(C)

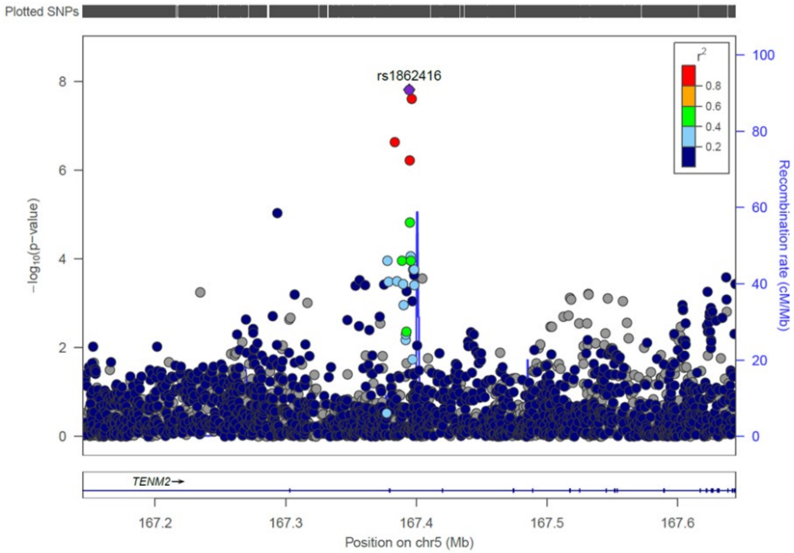
Plotted SNPs
rs1862416
r2
0.8
0.6
0.4
0.2
−log10(p−value)
Recombination rate (cM/Mb)
TENM2
167.2
167.3
167.4
167.5
167.6
Position on chr5 (Mb)

(D)

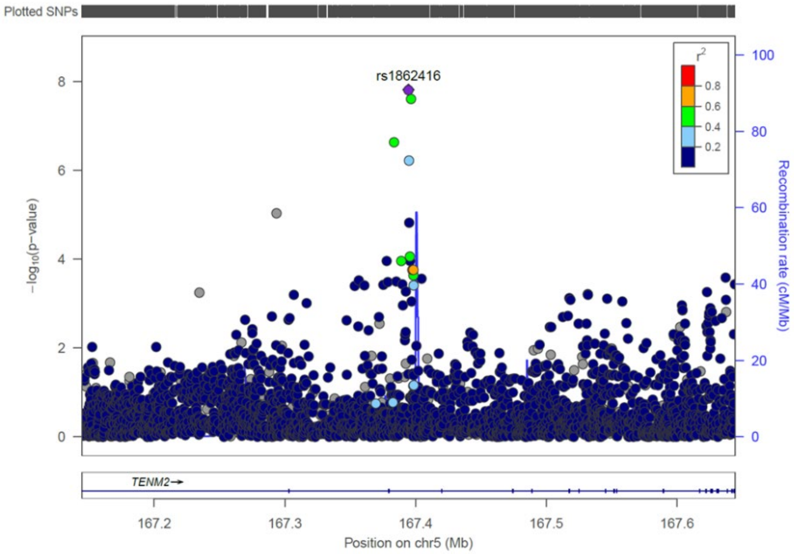
Plotted SNPs
rs1862416
r2
0.8
0.6
0.4
0.2
−log10(p−value)
Recombination rate (cM/Mb)
TENM2
167.2
167.3
167.4
167.5
167.6
Position on chr5 (Mb)

**Supplementary Figure 4**. Novel single nucleotide polymorphism (SNP) associations with nicotine dependence (ND) by study and ancestry group.

Associations are presented for the (A) *MAGI2/GNAI1* SNP allele rs2714700-T and (B) *TENM2* SNP allele rs1862416-T with severe vs. mild ND, by calculating odds ratio (OR) and 95% confidence interval (CI) estimates using the regression coefficients from the discovery genome-wide association study analyses of categorical FTND (i.e., OR=exp[$2\times\beta_{SNP}$] for severe vs. mild ND).

(A)

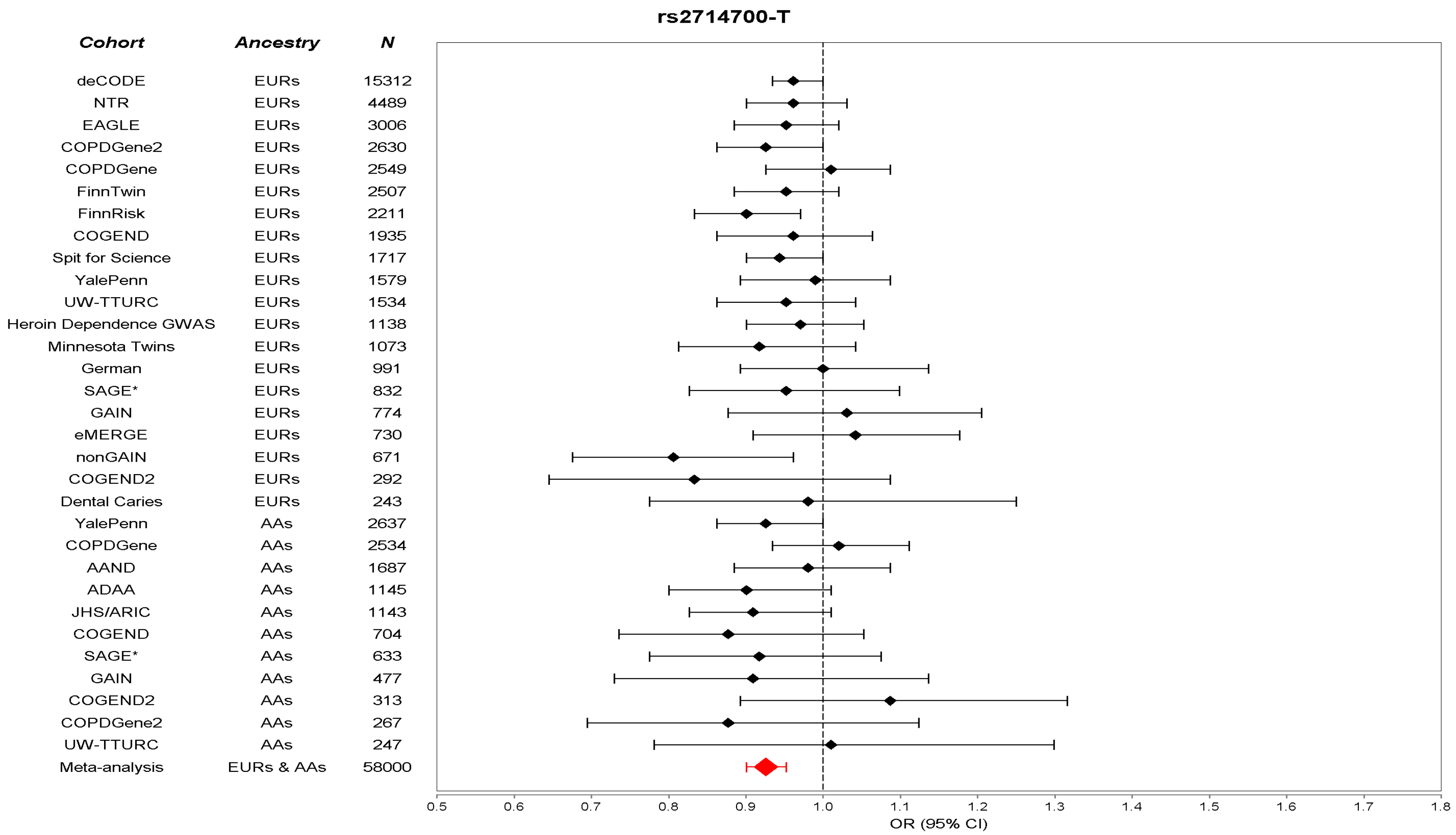

(B)

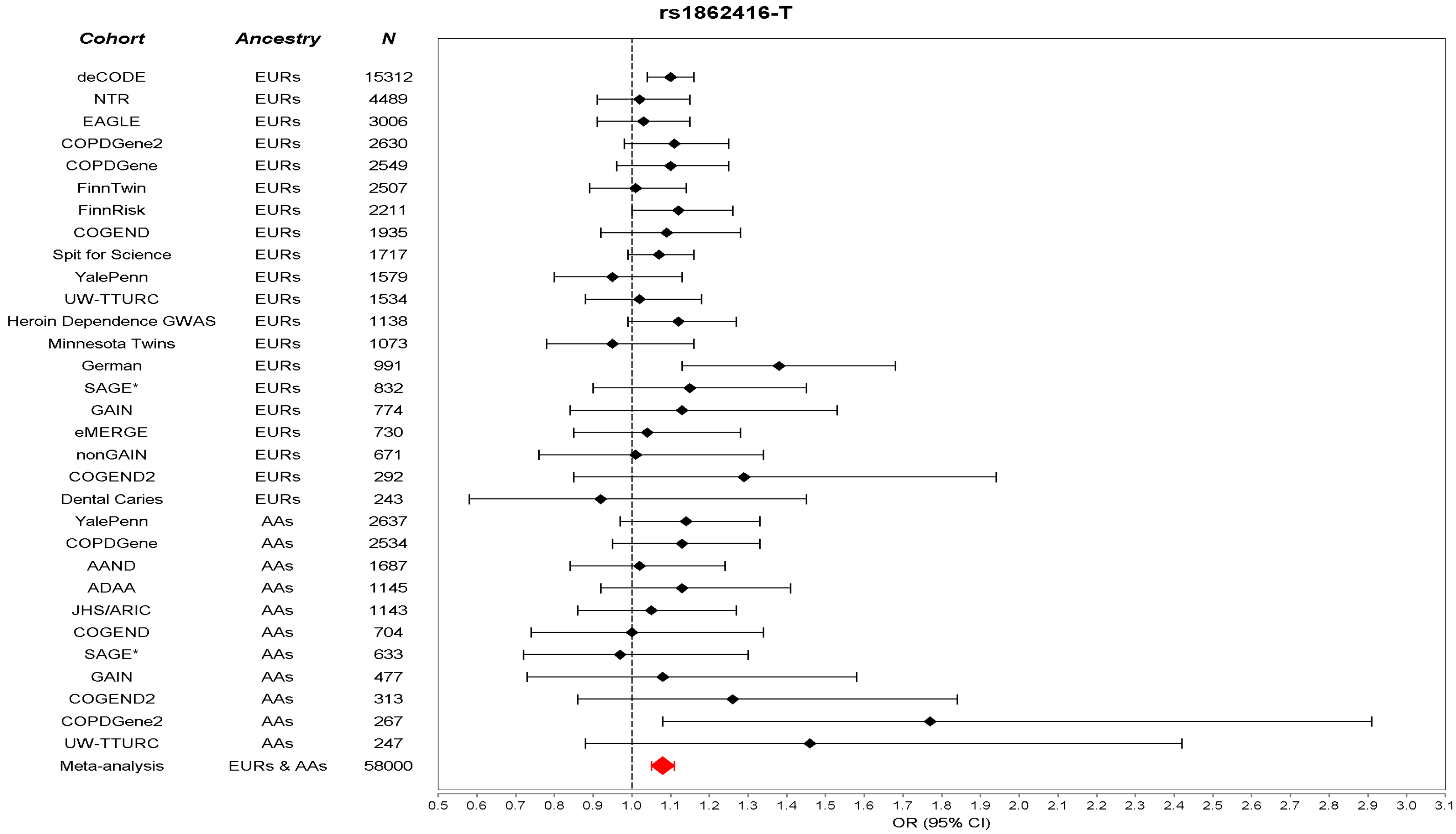
rs1862416-T
Cohort
Ancestry
N
deCODE EURs 15312
NTR EURs 4489
EAGLE EURs 3006
COPDGene2 EURs 2630
COPDGene EURs 2549
FinnTwin EURs 2507
FinnRisk EURs 2211
COGEND EURs 1935
Spit for Science EURs 1717
YalePenn EURs 1579
UW-TTURC EURs 1534
Heroin Dependence GWAS EURs 1138
Minnesota Twins EURs 1073
German EURs 991
SAGE* EURs 832
GAIN EURs 774
eMERGE EURs 730
nonGAIN EURs 671
COGEND2 EURs 292
Dental Caries EURs 243
YalePenn AAs 2637
COPDGene AAs 2534
AAND AAs 1687
ADAA AAs 1145
JHS/ARIC AAs 1143
COGEND AAs 704
SAGE* AAs 633
GAIN AAs 477
COGEND2 AAs 313
COPDGene2 AAs 267
UW-TTURC AAs 247
Meta-analysis EURs & AAs 58000
0.5 0.6 0.7 0.8 0.9 1.0 1.1 1.2 1.3 1.4 1.5 1.6 1.7 1.8 1.9 2.0 2.1 2.2 2.3 2.4 2.5 2.6 2.7 2.8 2.9 3.0 3.1
OR (95% CI)

**Supplementary Figure 5**. Posterior probability matrices for traits evaluated for shared genetics with ND using GWAS-PW at the 5 FTND-associated genome-wide significant loci.

For the 12 traits, variants in LD ($r^2$>0.2 in 1000 Genomes EUR populations) with the lead SNP from each genome-wide significant FTND locus was analyzed using GWAS-PW to find shared genetic influences between FTND and each trait. Shown are GWAS-PW posterior probabilities that the genomic region surrounding a lead GWAS SNP contains a variant that influences ND (Model 1); contains a variant that influences the other trait (Model 2); contains a variant that influences both traits (Model 3); or contains a variant that influences ND and a separate variant that influences the other trait (Model 4). The genomic position for each lead GWAS SNP is in reference to GRCh37.

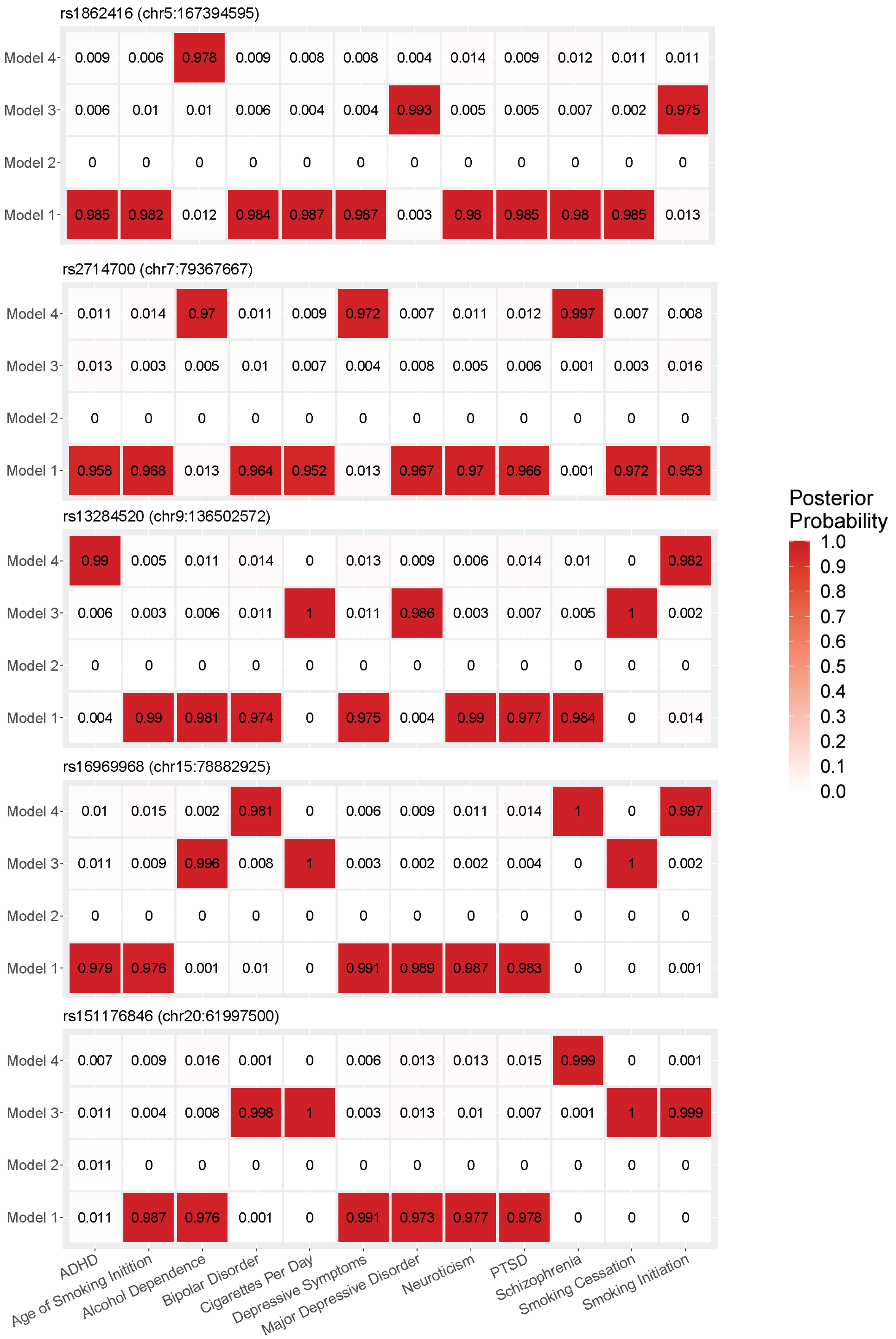
rs1862416 (chr5:167394595)
Model 4 0.009 0.006 0.978 0.009 0.008 0.008 0.004 0.014 0.009 0.012 0.011 0.011
Model 3 0.006 0.01 0.01 0.006 0.004 0.004 0.993 0.005 0.005 0.007 0.002 0.975
Model 2 0 0 0 0 0 0 0 0 0 0 0 0
Model 1 0.985 0.982 0.012 0.984 0.987 0.987 0.003 0.98 0.985 0.98 0.985 0.013
rs2714700 (chr7:79367667)
Model 4 0.011 0.014 0.97 0.011 0.009 0.972 0.007 0.011 0.012 0.997 0.007 0.008
Model 3 0.013 0.003 0.005 0.01 0.007 0.004 0.008 0.005 0.006 0.001 0.003 0.016
Model 2 0 0 0 0 0 0 0 0 0 0 0 0
Model 1 0.958 0.968 0.013 0.964 0.952 0.013 0.967 0.97 0.966 0.001 0.972 0.953
rs13284520 (chr9:136502572)
Model 4 0.99 0.005 0.011 0.014 0 0.013 0.009 0.006 0.014 0.01 0 0.982
Model 3 0.006 0.003 0.006 0.011 1 0.011 0.986 0.003 0.007 0.005 1 0.002
Model 2 0 0 0 0 0 0 0 0 0 0 0 0
Model 1 0.004 0.99 0.981 0.974 0 0.975 0.004 0.99 0.977 0.984 0 0.014
rs16969968 (chr15:78882925)
Model 4 0.01 0.015 0.002 0.981 0 0.006 0.009 0.011 0.014 1 0 0.997
Model 3 0.011 0.009 0.996 0.008 1 0.003 0.002 0.002 0.004 0 1 0.002
Model 2 0 0 0 0 0 0 0 0 0 0 0 0
Model 1 0.979 0.976 0.001 0.01 0 0.991 0.989 0.987 0.983 0 0 0.001
rs151176846 (chr20:61997500)
Model 4 0.007 0.009 0.016 0.001 0 0.006 0.013 0.013 0.015 0.999 0 0.001
Model 3 0.011 0.004 0.008 0.998 1 0.003 0.013 0.01 0.007 0.001 1 0.999
Model 2 0.011 0 0 0 0 0 0 0 0 0 0 0
Model 1 0.011 0.987 0.976 0.001 0 0.991 0.973 0.977 0.978 0 0 0
ADHD
Age of Smoking Initition
Alcohol Dependence
Bipolar Disorder
Cigarettes Per Day
Depressive Symptoms
Major Depressive Disorder
Neuroticism
PTSD
Schizophrenia
Smoking Cessation
Smoking Initiation
Posterior Probability
1.0
0.9
0.8
0.7
0.6
0.5
0.4
0.3
0.2
0.1
0.0

**Supplementary Figure 6**. Linkage disequilibrium (LD) matrix of rs1862416 (marked by blue boxes) and other *TENM2* single nucleotide polymorphisms (SNPs) included in the genome-wide association study (GWAS) catalog[87] (https://www.ebi.ac.uk) for their genome-wide significant associations ($P<5\times10^{-8}$). $r^2$ values, as obtained from LDlink[51], correspond to the 1000 Genomes European panel. Numerical values correspond to the originally reported GWAS: 1, educational attainment[63]; 2a, smoking initiation (ever vs. never smoking)[48,88-90]; 2b, age of smoking initiation[48]; 2c, smoking cessation (current vs. former smoking)[48]; 2d, alcohol consumption (drinks per week)[48]; 3, lung function[90,91]; 4, height[90]; 5, number of sexual partners[88]; 6, depression[92,93]; 7, risk taking tendency[88]; 8, body mass index[90]; 9, menarche (age at onset)[94]; 10, cigarette pack-years[95]; and 11, regular attendance at a religious group[96]. rs11739827, associated with alcohol consumption[48], was not available for comparison with rs1862416 in LDlink.

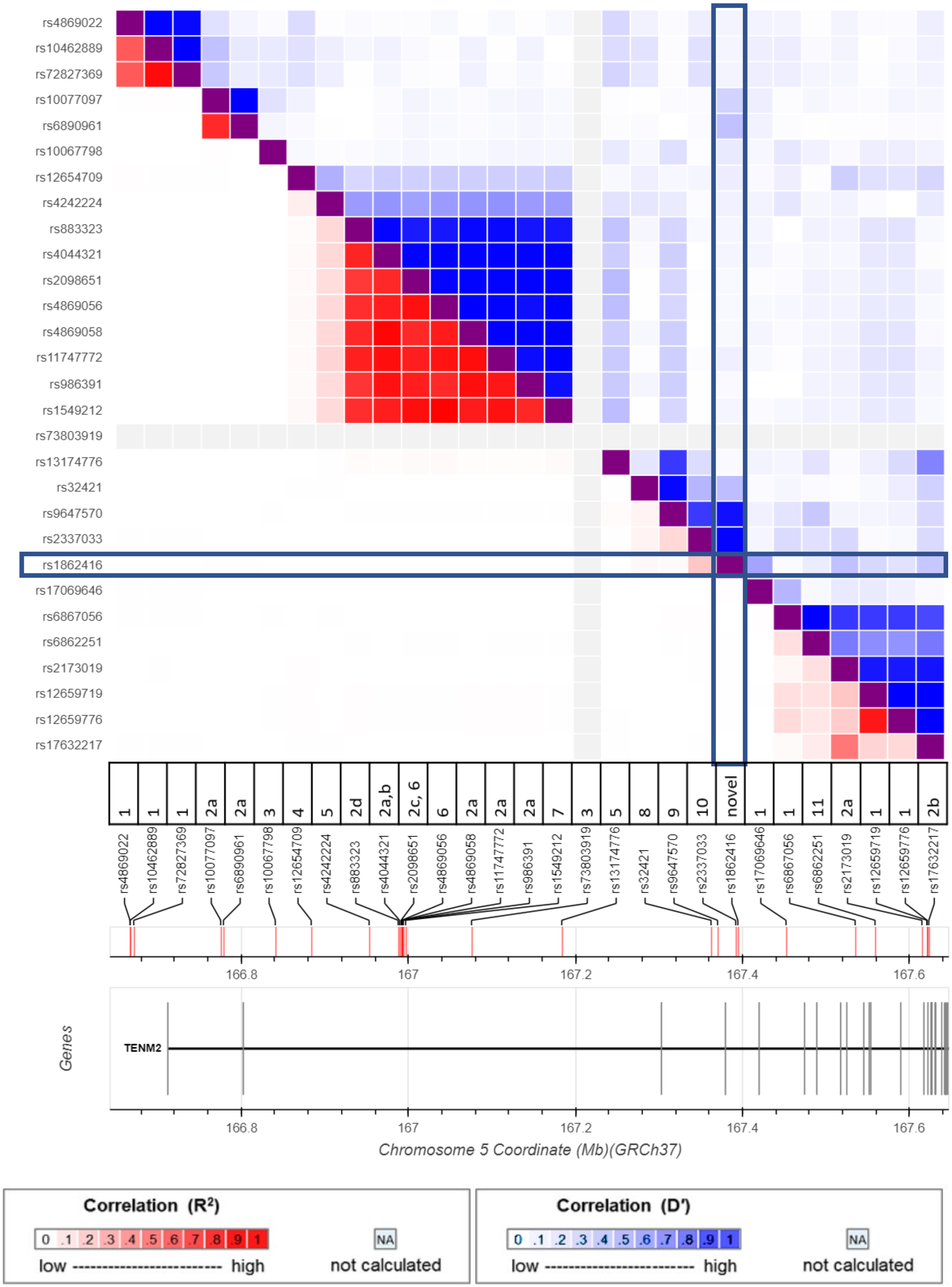
rs4869022
rs10462889
rs72827369
rs10077097
rs6890961
rs10067798
rs12654709
rs4242224
rs883323
rs4044321
rs2098651
rs4869056
rs4869058
rs11747772
rs986391
rs1549212
rs73803919
rs13174776
rs32421
rs9647570
rs2337033
rs1862416
rs17069646
rs6867056
rs6862251
rs2173019
rs12659719
rs12659776
rs17632217
1 1 1 2a 2a 3 4 5 2d 2a,b 2c, 6 6 2a 2a 2a 7 3 5 8 9 10 novel 1 1 11 2a 1 1 2b
166.8
167
167.2
167.4
167.6
Genes
TENM2
Chromosome 5 Coordinate (Mb)(GRCh37)
Correlation (R²)
0 .1 .2 .3 .4 .5 .6 .7 .8 .9 1
low high
NA
not calculated
Correlation (D')
0 .1 .2 .3 .4 .5 .6 .7 .8 .9 1
low high
NA
not calculated

### References

1. Hancock, D.B. *et al.* Genome-wide association study across European and African American ancestries identifies a SNP in DNMT3B contributing to nicotine dependence. *Mol Psychiatry* **23**, 1911-1919 (2018).
2. Hancock, D.B. *et al.* Genome-wide meta-analysis reveals common splice site acceptor variant in CHRNA4 associated with nicotine dependence. *Transl Psychiatry* **5**, e651 (2015).
3. Heatherton, T.F., Kozlowski, L.T., Frecker, R.C. & Fagerstrom, K.O. The Fagerstrom Test for Nicotine Dependence: a revision of the Fagerstrom Tolerance Questionnaire. *Br J Addict* **86**, 1119-27 (1991).
4. 1000 Genomes Project Consortium. *et al.* A global reference for human genetic variation. *Nature* **526**, 68-74 (2015).
5. Bierut, L.J. *et al.* Novel genes identified in a high-density genome wide association study for nicotine dependence. *Hum Mol Genet* **16**, 24-35 (2007).
6. Rice, J.P. *et al.* CHRNB3 is more strongly associated with Fagerstrom Test for Cigarette Dependence-based nicotine dependence than cigarettes per day: phenotype definition changes genome-wide association studies results. *Addiction* **107**, 2019-2028 (2012).
7. Laurie, C.C. *et al.* Quality control and quality assurance in genotypic data for genome-wide association studies. *Genet Epidemiol* **34**, 591-602 (2010).
8. Johnson, E.O. *et al.* Imputation across genotyping arrays for genome-wide association studies: assessment of bias and a correction strategy. *Hum Genet* **132**, 509-522 (2013).
9. Shaffer, J.R. *et al.* Genome-wide association scan for childhood caries implicates novel genes. *J Dent Res* **90**, 1457-62 (2011).
10. Regan, E.A. *et al.* Genetic epidemiology of COPD (COPDGene) study design. *COPD* **7**, 32-43 (2010).
11. Gulcher, J.R., Kristjansson, K., Gudbjartsson, H. & Stefansson, K. Protection of privacy by third-party encryption in genetic research in Iceland. *Eur J Hum Genet* **8**, 739-42 (2000).
12. Thorgeirsson, T.E. *et al.* A variant associated with nicotine dependence, lung cancer and peripheral arterial disease. *Nature* **452**, 638-42 (2008).
13. Sulem, P. *et al.* Identification of low-frequency variants associated with gout and serum uric acid levels. *Nat Genet* **43**, 1127-30 (2011).
14. Loh, P.R. *et al.* Efficient Bayesian mixed-model analysis increases association power in large cohorts. *Nat Genet* **47**, 284-90 (2015).
15. Bulik-Sullivan, B.K. *et al.* LD Score regression distinguishes confounding from polygenicity in genome-wide association studies. *Nat Genet* **47**, 291-5 (2015).
16. Landi, M.T. *et al.* A genome-wide association study of lung cancer identifies a region of chromosome 5p15 associated with risk for adenocarcinoma. *Am J Hum Genet* **85**, 679-91 (2009).
17. Landi, M.T. *et al.* Environment And Genetics in Lung cancer Etiology (EAGLE) study: an integrative population-based case-control study of lung cancer. *BMC Public Health* **8**, 203 (2008).
18. Glasheen, C. *et al.* Is the Fagerstrom test for nicotine dependence invariant across secular trends in smoking? A question for cross-birth cohort analysis of nicotine dependence. *Drug Alcohol Depend* **185**, 127-132 (2018).

19. Borodulin, K. *et al.* Forty-year trends in cardiovascular risk factors in Finland. *Eur J Public Health* **25**, 539-46 (2015).
20. Lim, E.T. *et al.* Distribution and medical impact of loss-of-function variants in the Finnish founder population. *PLoS Genet* **10**, e1004494 (2014).
21. Hallfors, J. *et al.* Genome-wide association study in Finnish twins highlights the connection between nicotine addiction and neurotrophin signaling pathway. *Addict Biol* (2018).
22. Kaprio, J. The Finnish Twin Cohort Study: an update. *Twin Res Hum Genet* **16**, 157-62 (2013).
23. Kaprio, J. Twin studies in Finland 2006. *Twin Res Hum Genet* **9**, 772-7 (2006).
24. Das, S. *et al.* Next-generation genotype imputation service and methods. *Nat Genet* **48**, 1284-1287 (2016).
25. Manolio, T.A. *et al.* New models of collaboration in genome-wide association studies: the Genetic Association Information Network. *Nat Genet* **39**, 1045-51 (2007).
26. Lindenberg, A. *et al.* The German multi-centre study on smoking-related behavior-description of a population-based case-control study. *Addict Biol* **16**, 638-53 (2011).
27. Sempos, C.T., Bild, D.E. & Manolio, T.A. Overview of the Jackson Heart Study: a study of cardiovascular diseases in African American men and women. *Am J Med Sci* **317**, 142-6 (1999).
28. The Atherosclerosis Risk in Communities (ARIC) Study: design and objectives. The ARIC investigators. *Am J Epidemiol* **129**, 687-702 (1989).
29. Miller, M.B. *et al.* The Minnesota Center for Twin and Family Research genome-wide association study. *Twin Res Hum Genet* **15**, 767-74 (2012).
30. Vrieze, S.I. *et al.* Rare nonsynonymous exonic variants in addiction and behavioral disinhibition. *Biol Psychiatry* **75**, 783-9 (2014).
31. Boomsma, D.I. *et al.* Netherlands Twin Register: from twins to twin families. *Twin Res Hum Genet* **9**, 849-57 (2006).
32. Willemsen, G. *et al.* The Netherlands Twin Register biobank: a resource for genetic epidemiological studies. *Twin Res Hum Genet* **13**, 231-45 (2010).
33. Minica, C.C. *et al.* Heritability, SNP- and Gene-Based Analyses of Cannabis Use Initiation and Age at Onset. *Behav Genet* **45**, 503-13 (2015).
34. Vink, J.M., Willemsen, G., Beem, A.L. & Boomsma, D.I. The Fagerstrom Test for Nicotine Dependence in a Dutch sample of daily smokers and ex-smokers. *Addict Behav* **30**, 575-9 (2005).
35. Lessov, C.N. *et al.* Defining nicotine dependence for genetic research: evidence from Australian twins. *Psychol Med* **34**, 865-79 (2004).
36. Edenberg, H.J. The collaborative study on the genetics of alcoholism: an update. *Alcohol Res Health* **26**, 214-8 (2002).
37. Bierut, L.J., Strickland, J.R., Thompson, J.R., Afful, S.E. & Cottler, L.B. Drug use and dependence in cocaine dependent subjects, community-based individuals, and their siblings. *Drug Alcohol Depend* **95**, 14-22 (2008).
38. Dick, D.M. *et al.* Spit for Science: launching a longitudinal study of genetic and environmental influences on substance use and emotional health at a large US university. *Front Genet* **5**, 47 (2014).

39. Harris, P.A. *et al.* Research electronic data capture (REDCap)--a metadata-driven methodology and workflow process for providing translational research informatics support. *J Biomed Inform* **42**, 377-81 (2009).
40. Peterson, R.E. *et al.* The utility of empirically assigning ancestry groups in cross-population genetic studies of addiction. *Am J Addict* **26**, 494-501 (2017).
41. Gelernter, J. *et al.* Genome-wide association study of alcohol dependence:significant findings in African- and European-Americans including novel risk loci. *Mol Psychiatry* **19**, 41-9 (2014).
42. Gelernter, J. *et al.* Genome-wide association study of cocaine dependence and related traits: FAM53B identified as a risk gene. *Mol Psychiatry* **19**, 717-723 (2014).
43. Gelernter, J. *et al.* Genome-wide association study of opioid dependence: multiple associations mapped to calcium and potassium pathways. *Biol Psychiatry* **76**, 66-74 (2014).
44. Gelernter, J. *et al.* Genome-wide association study of nicotine dependence in American populations: identification of novel risk loci in both African-Americans and European-Americans. *Biol Psychiatry* **77**, 493-503 (2015).
45. Hendershot, T. *et al.* Using the PhenX Toolkit to Add Standard Measures to a Study. *Curr Protoc Hum Genet* **86**, 1 21 1-1 21 17 (2015).
46. Gordon, D., Finch, S.J., Nothnagel, M. & Ott, J. Power and sample size calculations for case-control genetic association tests when errors are present: application to single nucleotide polymorphisms. *Hum Hered* **54**, 22-33 (2002).
47. Sinnott, J.A. *et al.* Improving the power of genetic association tests with imperfect phenotype derived from electronic medical records. *Hum Genet* **133**, 1369-82 (2014).
48. Liu, M. *et al.* Association studies of up to 1.2 million individuals yield new insights into the genetic etiology of tobacco and alcohol use. *Nat Genet* **51**, 237-244 (2019).
49. Boyles, A.L., Harris, S.F., Rooney, A.A. & Thayer, K.A. Forest Plot Viewer: a new graphing tool. *Epidemiology* **22**, 746-7 (2011).
50. American Society of Clinical Oncology. Heaviness of Smoking Index (HSI).
51. Machiela, M.J. & Chanock, S.J. LDlink: a web-based application for exploring population-specific haplotype structure and linking correlated alleles of possible functional variants. *Bioinformatics* **31**, 3555-7 (2015).
52. Yang, J. *et al.* Conditional and joint multiple-SNP analysis of GWAS summary statistics identifies additional variants influencing complex traits. *Nat Genet* **44**, 369-75, S1-3 (2012).
53. Yang, J., Lee, S.H., Goddard, M.E. & Visscher, P.M. GCTA: a tool for genome-wide complex trait analysis. *Am J Hum Genet* **88**, 76-82 (2011).
54. Barbeira, A.N. *et al.* Integrating predicted transcriptome from multiple tissues improves association detection. *PLoS Genet* **15**, e1007889 (2019).
55. Hibar, D.P. *et al.* Common genetic variants influence human subcortical brain structures. *Nature* **520**, 224-9 (2015).
56. McKay, J.D. *et al.* Large-scale association analysis identifies new lung cancer susceptibility loci and heterogeneity in genetic susceptibility across histological subtypes. *Nat Genet* **49**, 1126-1132 (2017).
57. Dastani, Z. *et al.* Novel loci for adiponectin levels and their influence on type 2 diabetes and metabolic traits: a multi-ethnic meta-analysis of 45,891 individuals. *PLoS Genet* **8**, e1002607 (2012).

58. Nikpay, M. *et al.* A comprehensive 1,000 Genomes-based genome-wide association meta-analysis of coronary artery disease. *Nat Genet* **47**, 1121-1130 (2015).
59. Ware, J.J. *et al.* Genome-Wide Meta-Analysis of Cotinine Levels in Cigarette Smokers Identifies Locus at 4q13.2. *Sci Rep* **6**, 20092 (2016).
60. Benyamin, B. *et al.* Childhood intelligence is heritable, highly polygenic and associated with FNBP1L. *Mol Psychiatry* **19**, 253-8 (2014).
61. Rietveld, C.A. *et al.* GWAS of 126,559 individuals identifies genetic variants associated with educational attainment. *Science* **340**, 1467-71 (2013).
62. Sniekers, S. *et al.* Genome-wide association meta-analysis of 78,308 individuals identifies new loci and genes influencing human intelligence. *Nat Genet* **49**, 1107-1112 (2017).
63. Lee, J.J. *et al.* Gene discovery and polygenic prediction from a genome-wide association study of educational attainment in 1.1 million individuals. *Nat Genet* **50**, 1112-1121 (2018).
64. Walters, R.K. *et al.* Transancestral GWAS of alcohol dependence reveals common genetic underpinnings with psychiatric disorders. *Nat Neurosci* **21**, 1656-1669 (2018).
65. Demontis, D. *et al.* Genome-wide association study implicates CHRNA2 in cannabis use disorder. *Nat Neurosci* **22**, 1066-1074 (2019).
66. Pasman, J.A. *et al.* GWAS of lifetime cannabis use reveals new risk loci, genetic overlap with psychiatric traits, and a causal influence of schizophrenia. *Nat Neurosci* **21**, 1161-1170 (2018).
67. Lambert, J.C. *et al.* Meta-analysis of 74,046 individuals identifies 11 new susceptibility loci for Alzheimer's disease. *Nat Genet* **45**, 1452-8 (2013).
68. van Rheenen, W. *et al.* Genome-wide association analyses identify new risk variants and the genetic architecture of amyotrophic lateral sclerosis. *Nat Genet* **48**, 1043-8 (2016).
69. Simon-Sanchez, J. *et al.* Genome-wide association study reveals genetic risk underlying Parkinson's disease. *Nat Genet* **41**, 1308-12 (2009).
70. de Moor, M.H. *et al.* Meta-analysis of genome-wide association studies for personality. *Mol Psychiatry* **17**, 337-49 (2012).
71. Okbay, A. *et al.* Genetic variants associated with subjective well-being, depressive symptoms, and neuroticism identified through genome-wide analyses. *Nat Genet* **48**, 624-33 (2016).
72. Watson, H.J. *et al.* Genome-wide association study identifies eight risk loci and implicates metabo-psychiatric origins for anorexia nervosa. *Nat Genet* **51**, 1207-1214 (2019).
73. Demontis, D. *et al.* Discovery of the first genome-wide significant risk loci for attention deficit/hyperactivity disorder. *Nat Genet* **51**, 63-75 (2019).
74. Grove, J. *et al.* Identification of common genetic risk variants for autism spectrum disorder. *Nat Genet* **51**, 431-444 (2019).
75. Stahl, E.A. *et al.* Genome-wide association study identifies 30 loci associated with bipolar disorder. *Nat Genet* **51**, 793-803 (2019).
76. Howard, D.M. *et al.* Genome-wide meta-analysis of depression identifies 102 independent variants and highlights the importance of the prefrontal brain regions. *Nat Neurosci* **22**, 343-352 (2019).

77. Nievergelt, C.M. *et al.* International meta-analysis of PTSD genome-wide association studies identifies sex- and ancestry-specific genetic risk loci. *Nat Commun* **10**, 4558 (2019).
78. Smoller, J.W. *et al.* Identification of risk loci with shared effects on five major psychiatric disorders: a genome-wide analysis. *Lancet* **381**, 1371-9 (2013).
79. Schizophrenia Working Group of the Psychiatric Genomics Consortium. Biological insights from 108 schizophrenia-associated genetic loci. *Nature* **511**, 421-7 (2014).
80. Hobbs, B.D. *et al.* Genetic loci associated with chronic obstructive pulmonary disease overlap with loci for lung function and pulmonary fibrosis. *Nat Genet* **49**, 426-432 (2017).
81. Wain, L.V. *et al.* Genome-wide association analyses for lung function and chronic obstructive pulmonary disease identify new loci and potential druggable targets. *Nat Genet* **49**, 416-425 (2017).
82. Wellcome Trust Case Control ConsortiumG. *et al.* Bayesian refinement of association signals for 14 loci in 3 common diseases. *Nat Genet* **44**, 1294-301 (2012).
83. Pruim, R.J. *et al.* LocusZoom: regional visualization of genome-wide association scan results. *Bioinformatics* **26**, 2336-7 (2010).
84. GTEx Consortium. Human genomics. The Genotype-Tissue Expression (GTEx) pilot analysis: multitissue gene regulation in humans. *Science* **348**, 648-60 (2015).
85. BrainSeq: A Human Brain Genomics Consortium. BrainSeq: Neurogenomics to Drive Novel Target Discovery for Neuropsychiatric Disorders. *Neuron* **88**, 1078-1083 (2015).
86. Ward, L.D. & Kellis, M. HaploReg: a resource for exploring chromatin states, conservation, and regulatory motif alterations within sets of genetically linked variants. *Nucleic Acids Res* **40**, D930-4 (2012).
87. Welter, D. *et al.* The NHGRI GWAS Catalog, a curated resource of SNP-trait associations. *Nucleic Acids Res* **42**, D1001-6 (2014).
88. Karlsson Linner, R. *et al.* Genome-wide association analyses of risk tolerance and risky behaviors in over 1 million individuals identify hundreds of loci and shared genetic influences. *Nat Genet* **51**, 245-257 (2019).
89. Erzurumluoglu, A.M. *et al.* Meta-analysis of up to 622,409 individuals identifies 40 novel smoking behaviour associated genetic loci. *Mol Psychiatry* (2019).
90. Kichaev, G. *et al.* Leveraging Polygenic Functional Enrichment to Improve GWAS Power. *Am J Hum Genet* **104**, 65-75 (2019).
91. Lutz, S.M. *et al.* A genome-wide association study identifies risk loci for spirometric measures among smokers of European and African ancestry. *BMC Genet* **16**, 138 (2015).
92. Nagel, M. *et al.* Meta-analysis of genome-wide association studies for neuroticism in 449,484 individuals identifies novel genetic loci and pathways. *Nat Genet* **50**, 920-927 (2018).
93. Wray, N.R. *et al.* Genome-wide association analyses identify 44 risk variants and refine the genetic architecture of major depression. *Nat Genet* **50**, 668-681 (2018).
94. Perry, J.R. *et al.* Parent-of-origin-specific allelic associations among 106 genomic loci for age at menarche. *Nature* **514**, 92-97 (2014).
95. Buchwald, J. *et al.* Genome-wide association meta-analysis of nicotine metabolism and cigarette consumption measures in smokers of European descent. *Mol Psychiatry* (2020).
96. Day, F.R., Ong, K.K. & Perry, J.R.B. Elucidating the genetic basis of social interaction and isolation. *Nat Commun* **9**, 2457 (2018).
